# Large scale genomic and evolutionary study reveals SARS-CoV-2 virus isolates from Bangladesh strongly correlate with European origin and not with China

**DOI:** 10.1101/2021.01.17.425424

**Authors:** Mohammad Fazle Alam Rabbi, Md. Imran Khan, Saam Hasan, Mauricio Chalita, Kazi Nadim Hasan, Abu Sufian, Md. Bayejid Hosen, Mohammed Nafiz Imtiaz Polol, Jannatun Naima, Kihyun Lee, Yeong Ouk Kim, Mamudul Hasan Razu, Mala khan, Md. Mizanur Rahman, Jongsik Chun, Md. Abdul Khaleque, Nur A. Hasan, Rita R Colwell, Sharif Akhteruzzaman

**Author notes:** Corresponding author email address: Sharif Akhteruzzaman.

## Abstract

**Rationale:** The global public health is in serious crisis due to emergence of SARS-CoV-2 virus. Studies are ongoing to reveal the genomic variants of the virus circulating in various parts of the world. However, data generated from low- and middle-income countries are scarce due to resource limitation. This study was focused to perform whole genome sequencing of 151 SARS-CoV-2 isolates from COVID-19 positive Bangladeshi patients. The goal of this study was to identify the genomic variants among the SARS-CoV-2 virus isolates in Bangladesh, to determine the molecular epidemiology and to develop a relationship between host clinical trait with the virus genomic variants.

**Method:** Suspected patients were tested for COVID-19 using one step commercial qPCR kit for SARS-CoV-2 Virus. Viral RNA was extracted from positive patients, converted to cDNA which was amplified using Ion AmpliSeq™ SARS-CoV-2 Research Panel. Massive parallel sequencing was carried out using Ion AmpliSeq™ Library Kit Plus. Assembly of raw data is done by aligning the reads to a pre-defined reference genome (NC_045512.2) while retaining the unique variations of the input raw data by creating a consensus genome. A random forest-based association analysis was carried out to correlate the viral genomic variants with the clinical traits present in the host.

**Result:** Among the 151 viral isolates, we observed the 413 unique variants. Among these 8 variants occurred in more than 80 % of cases which include 241C to T, 1163A to T, 3037C to T,14408C to T, 23403A to G, 28881G to A, 28882 G to A, and finally the 28883G to C. Phylogenetic analysis revealed a predominance of variants belonging to GR clade, which have a strong geographical presence in Europe, indicating possible introduction of the SARS-CoV-2 virus into Bangladesh through a European channel. However, other possibilities like a route of entry from China cannot be ruled out as viral isolate belonging to L clade with a close relationship to Wuhan reference genome was also detected. We observed a total of 37 genomic variants to be strongly associated with clinical symptoms such as fever, sore throat, overall symptomatic status, etc. (Fisher’s Exact Test p-value<0.05). The most mention-worthy among those were the 3916CtoT (associated with causing sore throat, p-value 0.0005), the 14408C to T (associated with protection from developing cough, p-value= 0.027), and the 28881G to A, 28882G to A, and 28883G to C variant (associated with causing chest pain, p-value 0.025).

**Conclusion:** To our knowledge, this study is the first large scale phylogenomic studies of SARS-CoV-2 virus circulating in Bangladesh. The observed epidemiological and genomic features may inform future research platform for disease management, vaccine development and epidemiological study.

## 1. Introduction

In December 2019, several cases of unknown pneumonia were reported in the Hubei province of China which raised concerns among world health experts^1^. The aetiology was later diagnosed as a novel coronavirus and was dubbed by Chinese authorities as “COVID-19” or “2019-nCoV”^2,3^. The virus was later designated as Severe Acute Respiratory Syndrome Coronavirus 2 (SARS-CoV-2) based on taxonomic and genetic relationship with the previously identified SARS-CoV virus^4^. It’s high rate of transmissibility ^5,6^ allowed the virus to achieve global transmission rapidly^7,8^. The World Health Organization (WHO) announced COVID-19 as a pandemic on March 11, 2020^9^. As of December 14, 2020 nearly 70 million infections have been reported with over 1.6 million deaths^10^.

SARS-CoV-2 is a positive-sense single-stranded RNA virus believed to be transmitted in aerosols and common surfaces^11^. It has proven adept at transmission and possesses a strong pathogenic capacity, heightening need for in-depth understanding of its genetic characteristics. A significant effort is ongoing to develop an effective vaccine against this virus. Hence understanding it’s key genomic features and variants is important^12,13^. A large amount of information has accumulated with regard to genomic and proteomic variants found subtypes of virus mainly circulating in developed parts of the world^5,14,15^. However, information on molecular variants of the SARS-CoV-2 virus circulating in low- and middle-income countries (LMIC) is sparse.

Bangladesh, a developing country in South Asia, is one of the most densely populated (over 1000 people/km^2^)^16^. The first COVID-19 case in Bangladesh was reported on March 08, 2020^17^. Since then, the country has suffered a steeply rising number of new COVID-19 cases. As of October 19, 2020, nearly 400,000 COVID-19 cases have been reported, and more than 5,500 people have died^18^. The country also has a large population who are settled and work abroad, especially in the Middle East and Europe^19^. These workers tend to visit families in Bangladesh during the summer season, March to June. Furthermore, China is a strategic partner in economic development of Bangladesh, with significant traffic between these two countries^20^. These factors indicate possible routes by which the virus entered Bangladesh.

Whole-genome sequences of the SARS-CoV-2 virus have been publicly deposited, the majority of which can be accessed from the Global Initiative on Sharing All Influenza Data (GISAID). As of October 19^th^, 2020, more than 130,000 viral sequences have been uploaded. GISAID classifies them into 7 clades based on single nucleotide polymorphism profiles. These compose clade S (C8782T, T28144C includes NS8-L84S), clade L (C241, C3037, A23403, C8782, G11083, G25563, G26144, T28144, G28882), clade V (G11083T, G26144T, NSP6-L37F + NS3-G251V), clade G (C241T, C3037T, A23403G includes S-D614G), clade GH (C241T, C3037T, A23403G, G25563T includes S-D614G + NS3-Q57H) and clade GR (C241T, C3037T, A23403G, G28882A includes S-D614G + N-G204R) ^21^.

A large body of SARS-CoV-2 genomic information is available on strains from the developed world, but comparatively less information is available from resource-poor countries like Bangladesh^22^. The published literature focusing on SARS-CoV-2 genome biology from Bangladesh is comparatively limited. In this study, we sequenced and analysed comprehensively the whole genomes of 151 SARS-CoV-2 strains from patients suffering from varying degrees of disease severity. To compare Bangladeshi isolates with those from elsewhere, a comparative analysis that involved additional SARS-CoV-2 genomes of strains concurrently circulated in other parts of the world. A comparative study of this large number of samples has not yet been published from Bangladesh to the best of our knowledge. Finally, we conducted machine learning based analysis focused on the association of SARS-CoV-2 individual variants and clinically relevant parameters such as symptomatic status, fever, etc.

## 2. Methodology

### 2.1 Ethical approval

The cross-sectional study comprised 151 Bangladeshi patients diagnosed COVID-19 positive, based on real-time reverse transcriptase PCR (rRT-PCR). Samples were collected from the outdoor patient department (OPD) of the Central Police Hospital (CPH) and other tertiary medical centers in Dhaka, Bangladesh from April 28, 2020 to July 21, 2020. DNA Solution Ltd. (DNAS), in collaboration with the Designated Reference Institute for Chemical Measurements (DRICM) in Dhaka, Bangladesh provided the COVID-19 diagnosis and carried out subsequent whole-genome sequencing. All procedures in the study were according to ethical standards of the Helsinki Declaration of 1975, as revised in 2000^23^. Informed consent was obtained from each individual providing samples. The study protocol was approved by Bangladesh Council of Scientific and Industrial Research’s (BCSIR) ethics review committee (Ref No# 5600.8400.02.037.20).

### 2.2 Sample collection and real-time PCR

Oro-pharyngeal swabs from suspected patients were collected in virus transport medium (VTM) and transported in a cool box to DNA Solution Ltd. The samples were tested for SARS-CoV-2 RNA using a commercial one-step real-time COVID-19 PCR kit (Sansure Biotech Inc., Changsha, China) following manufacturer’s instruction. The real-time PCR kit uses PCR-Fluorescence probing technology and targets two genes, ORF 1 ab and conserved coding regions of the nucleocapsid protein N gene. Positive internal control of human Ribonuclease P (RNAase P), along with positive and negative control were used to nullify presence of PCR inhibitors. Real-time PCR was carried out in ABI7500 Fast DX instrument (Thermo Fisher Scientific, Massachusetts, USA). Samples with ct value of less than 30 for both target genes were selected for subsequent viral RNA isolation and whole-genome sequencing.

### 2.3 RNA extraction and cDNA preparation

RNA extraction was carried out using QIAamp® DSP Virus Spin Kit (Qiagen, Hilden, Germany) according to the instruction manual. Briefly, 200 μl of VTM containing the oropharyngeal swab was employed as starting material for viral RNA extraction using Silica-membrane technology. The samples were lysed, binding to the silica-membrane column, washed to remove contaminants, and eluted with RNase-free elution buffer. cDNA was prepared the same day, both the random hexamers and oligo dT primers were used, by using ProtoScript® II First Strand cDNA Synthesis Kit (NEB, Ipswich, MA, USA). Prepared cDNA was stored at −20◻C until further use.

### 2.4 Library preparation and whole-genome sequencing

Ion AmpliSeq™ SARS-CoV-2 Research Panel (Thermo Fisher Scientific, Massachusetts, USA) was employed to amplify the SARS-CoV-2 genome using prepared cDNA as template. The panel contains 237 pairs of specific primers covering more than 99% of the SARS-CoV-2 genome. Amplified fragments were carried forward to prepare libraries for massive parallel sequencing using Ion AmpliSeq™ Library Kit Plus (Thermo Fisher Scientific, Massachusetts, USA), following manufacturer’s instructions. Each of the prepared libraries was diluted to 100pM and pooled together for clonal amplification on the Ion One Touch 2 instrument. Clonally amplified libraries were enriched on using Ion One Touch ES followed by loading the enriched libraries on an Ion 530 chip. 15-20 samples were multiplexed simultaneously on the 530 chip during each run.

### 2.5 Bioinformatic Analysis

Genome assembly of the raw data was performed by the EzCOVID19^24^ cloud service provided on the EzBioCloud website^25^. Assembly is done by aligning reads to a pre-defined reference genome (NC_045512.2), while retaining the unique variations of the input raw data by creating a consensus genome. The consensus genome was then compared against the same reference genome to calculate single nucleotide variations (SNV) and positions. The SNVs were compared against GISAID clade variation markers^26^. **Figure 1** was generated by extracting all SNVs provided by EzCOVID19 and using RAxML ^27^using all default parameters.

**Figure 1:**
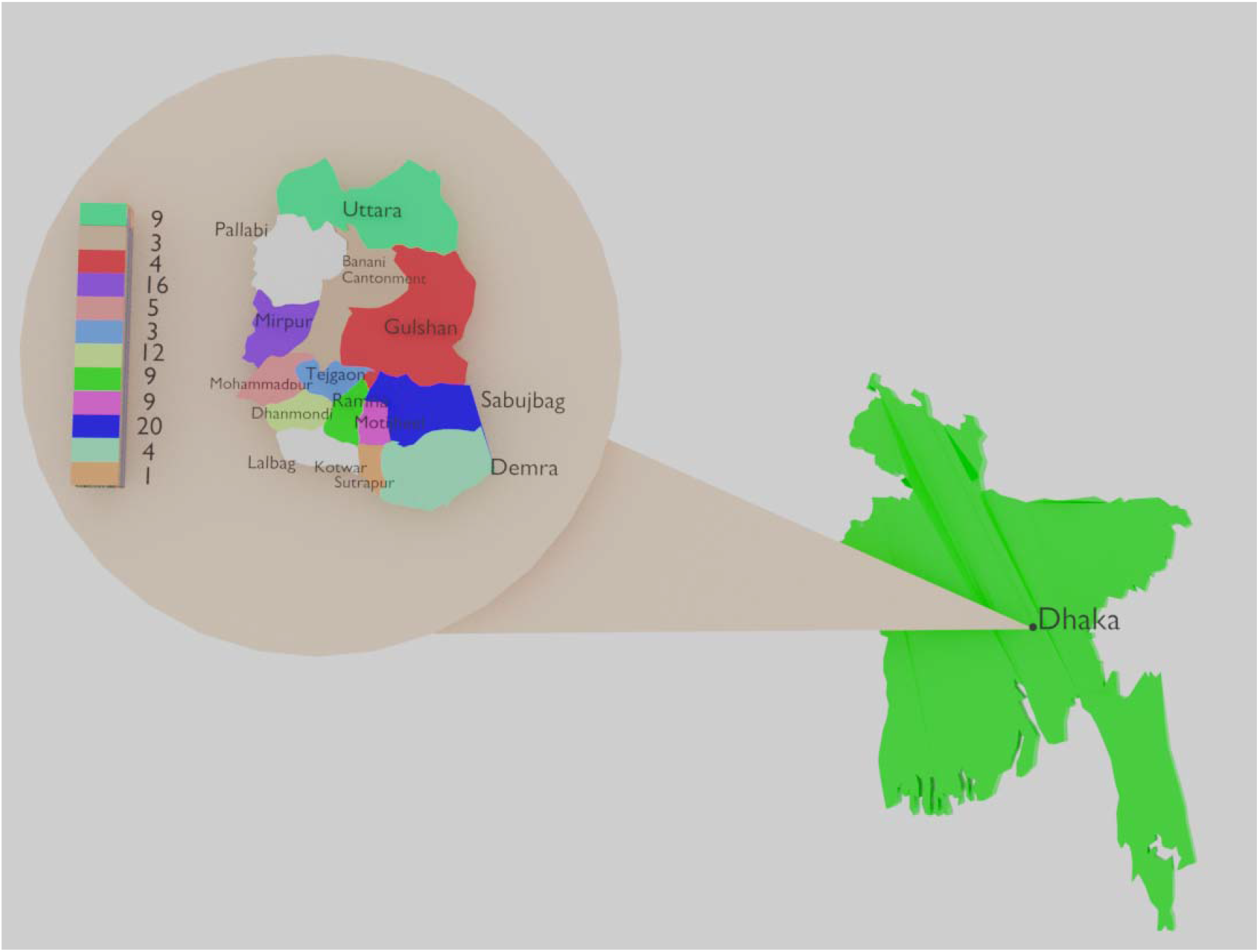
Schematic diagram of geographical distribution of COVID-19 samples. A total of 151 COVID-19 positive patients were considered in this study. Among them, 95 individuals gave consent to reveal their geographical location. Among them, it is clear the majority of samples came from the communities in or around the Sabujbag, Dhanmondi, Mirpur, Uttara and Ramna areas. A relatively fewer numbers of samples came from the central and southern most regions of Dhaka.

The phylogenetic and group typing analysis was accomplished using the EzCOVID19 cloud service, where a pre-build type grouping system based on occurrence of signature variants was provided. In brief, EzCOVID19 considered 2,761 SARS-CoV-2 genomes available at GISAID until April 01, 2020. Using a pairwise alignment approach (Myer-Miller’s method) each/all 2,761 genomes matched against Wuhan-Hu-1 reference genome (NC_045512.2) were aligned. From the resulting alignment, homopolymeric stretches of bases that cause frameshift errors were manually removed. The alignment matrix was then searched for variant sites and, in this process, sites at which >= 99% genomes showed a valid nucleotide character (not gap or ambiguous) were used. Among the variant positions, sites with >= 1% minor allele frequency (to avoid using sites that only have infrequent/spurious mutations) were selected. This resulted in 41 SNV sites (T514, C1059, G1397, G1440, C2416, A2480,C2558, G2891, C3037, C8782, T9477, C9962, A10323, G11083, C14408, C14724, C14805, C15324, T17247, C17747, A17858, C18060, C18877, A20268, T21584, A23403, G25563, G25979, G26144, A26530, C27046, T28144, C28657, T28688, G28851, C28863, G28881, G28882, G28883, C29095, G29553) which resulted in 88 unique allele combinations considered types. All isolates were typed according to typing system and isolates belonging to the same group were considered a similar study group.

The clinical importance of each of the variant present in our samples was assessed. To do this clinical metadata for 104 of the 151 individuals providing samples were collected Clinical parameters included fever, skin rash, diarrhoea, sore throat, chest pain, pneumonia, cough, anorexia, redness and itching of eyes, and overall symptomatic status of each individual (i.e. asymptomatic, mildly symptomatic, or severely symptomatic).

A random forest model was implemented to determine association between all variants and each clinical factor^28^. Variants classified as important determinants of the category variable (clinical trait in question) were selected for further statistical analysis. Chi square and fisher’s exact test were performed for each variant to establish whether effect of the presence of each variant was significant, with respect to the selected clinical factor. The p-value threshold for significance was set to 0.05.

## 3. Results

### 3.1 Data quality

A total of 151 individuals positive for SARS-CoV-2 participated in this study. Patient gender, age, and geographical region were requested for all positive individuals. Although all positive individuals provide written consent to participate in the study, only 104 individuals provided metadata information. Among those 104 individuals, sixty-five (65) were male and 39 were female. The mean age was 41.09 ± 1.75 (SEM). Geographical data analysis showed that the samples considered in this study were scattered throughout Dhaka city (**Figure 1)**, which is the capital and most densely populated city of Bangladesh^29^.

For all isolates from patients (n=151) more than 98% of the sequence reads generated aligned successfully to the reference genome (NC_045512.2). Similarly, coverage for each genome was determined showing that their range was between 800X to more than 6000X, with a mean of 3000X. These indicated that all data generated in this study were high quality and could be further analysed with confidence.

### 3.2 Phylogenetic distribution of Bangladeshi isolates into distinct GISAID clades

Phylogenetic analysis of 151 SARS-CoV-2 genome sequences decoded in this study along with concurrent reference isolates, classified Bangladeshi isolates into three paraphyletic clades according to GISAID clade classification system (**Figure 2**). This system classifies all SARS-CoV-2 isolates into 7 different clades defined by the presence of unique SNP signatures.. Among the genome of 151 SARS-CoV-2 Bangladeshi isolates, 132 genomes (87.4%) were placed into the GR clade followed by 13 genomes (8.7%) to G, and 6 genomes (3.9%) to GH clade. This appearance coincided with the recent deposition of genome sequences from Asia, where the majority of newly deposited genomes also were in the GR clade, followed by GH, G, and O^30^. An additional randomly selected 20 viral genomes were deposited in the GISAID database by various laboratories in Bangladesh. The clade distribution pattern of these other Bangladeshi isolates was similar to our sample sets (17 belong to GR clade, and each of other 3 belonging to G, S and L clade). Among them one of the isolate (Gene Accession No. MT5664683.1) belonging to the L clade was an interesting observation, mostly due to the fact that it shared very high similarity with the Wuhan reference genome (NC_045512.2). GH, GR and G are the clades with the most member isolates worldwide. The V clade isolates are rarer but have discovered in Asia both previously and more recently. However, the L clade has been very rare in Asia recently and the scenario coincides with the Bangladesh isolates except the one isolate considered in our study. The sequencing data of 151 SARS-CoV-2 virus genome sequences used in this study were uploaded in NCBI Gene Bank database and the associated accession number are summarized in **Supplementary Table 1**.

**Figure 2:**
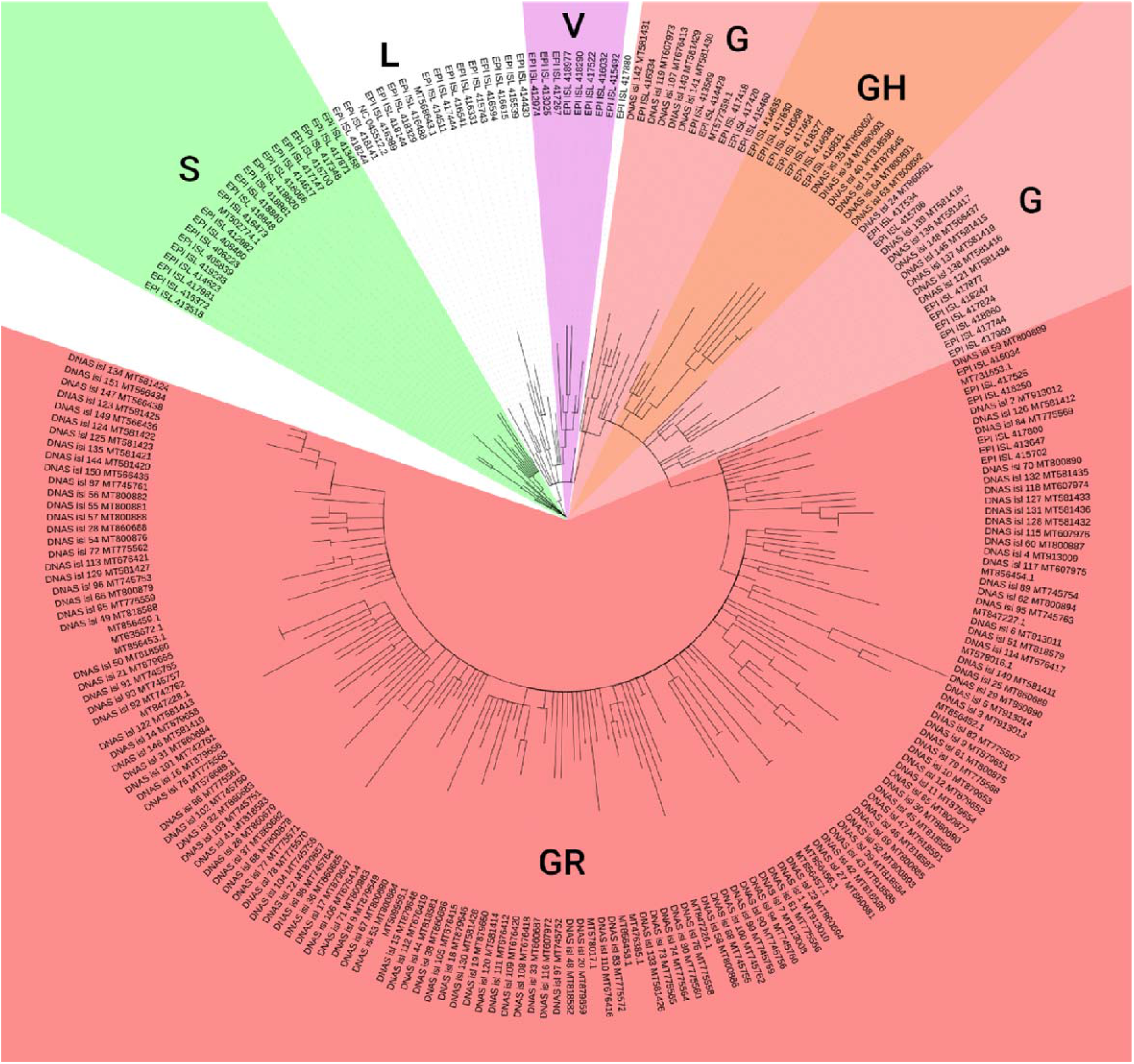
Radial SNP based phylogenetic tree displaying the classification of 151 SARS-CoV-2 virus isolates according to GISAID clade classification system. 132 out of 151 isolates considered in this study belonged to GR clade, 13 belonged to G clade and rest belonged to GH clade. Among the 20 randomly picked isolates from other Bangladeshi laboratories, 17 belonged to GR clade, 1 belonged to G clade, 1 to S clade and another to L clade which is where the Wuhan reference (NC_045512.2) genome is placed. This is in line with the recent identification and deposition of genomes from Asia. Majority of recently deposited Asian isolates have belonged to the GR clade.

### 3.3 Variant analysis

The 151 isolates contained a combined total of 1753 single nucleotide variants (SNVs); minus those which occurred within ambiguous codons and were not considered for further analysis (**Table 1**). These variants were spread across the 412 positions in the SARS-CoV-2 genome. Eight of these variants occurred in more than 79% of the isolates (**Figure 3A**), namely C to T change at 241, the A to T change at 1163, the C to T change at 3037 and at 14408, the A to G change at 23403, G to A change at 28881 and 28882, and finally the G to C change at 28883. Among these, 241 C to T is a 5’ UTR SNP mutation, whereas 23403 A to G is a synonymous and all others are nonsynonymous. As further validation, all these variants were also present in the other Bangladeshi isolates we included in our analysis (**Figure 3B)**. 1600 (91.27%) of our variants occurred within the coding regions, while the rest occurred in UTR regions. Among the variants that occur within genes on 1179 (73.68%) of them were nonsynonymous, resulting in amino acid changes. This included 244 unique variants and 242 unique positions where base substitutions led to said variants. Among one hundred and fifty one (151) SARS-CoV-2 virus isolates, DNAS_isl_29_MT860690 had the highest number of variants (total eighteen; 18), whereas, DNAS_isl_136_MT581417, DNAS_isl_138_MT581416 and DNAS_isl_148_MT566437 had the least number of variants (only six; 6). Among our isolates, DNAS_isl_29_MT860690 had the highest number of non-synonymous mutation (total fifteen; 15). The amino acid variants with the highest occurrence were the D to G change in the S protein caused by the variant at nucleotide position 23403, the R to K and G to R in the N protein caused by the multiple nucleotide variant from nucleotide position 28881 to 28883, the P to L in ORF1ab caused by the 14408 variant, and the I to F change also in ORF1ab resulting from the variant at nucleotide position 1163. All of these variants were in over 120 (79%) of our samples. There was one position with more than one kind of base substitution among our samples. This was position 28727 where we found a G to A change as well as a G to T change, occurring in different samples. For amino acid variants, there were two positions in the genome where multiple variants led to different amino acid changes. First was the aforementioned 28727, where different variants led either to an A to S or an A to T change in the N protein. The other was the amino acid substitution caused by the 28883 variant. While this variant occurs alongside those at 28881 and 28882 as an MNV, the 28883 base is part of a different codon than the other two. In most cases the G to C variant at this position led to a G to R substitution in the N gene product. However there was one isolate which also contained a variant at position 28884. This altered the resultant amino acid substitution to a G to Q change instead (**Supplementary Table 2**).

**Table 1.** Variant frequency observed in SARS-CoV-2 virus isolates. A total of 151 SARS-CoV-2 virus isolates carried 1753 total variants. Among these, 1179 were nonsynonymous mutations, 421 were synonymous mutations and 153 were 5'UTR SNP mutations. DNAS_isl_29_MT860690 had the highest number of variants (18), whereas, DNAS_isl_136_MT581417, DNAS_isl_138_MT581416 and DNAS_isl_148_MT566437 had the least number of variants (6). DNAS_isl_29_MT860690 had the highest number of non-synonymous mutation (15).

| Isolate Name | Variant Type |  |  | Total Variants |
| --- | --- | --- | --- | --- |
|  | Nonsynonymous | Synonymous | 5'UTR SNP |  |
| DNAS_isl_1_MT913010 | 8 | 4 | 2 | 14 |
| DNAS_isl_2_MT913012 | 8 | 3 | 1 | 12 |
| DNAS_isl_3_MT913013 | 13 | 3 | 1 | 17 |
| DNAS_isl_4_MT913009 | 8 | 2 | 1 | 11 |
| DNAS_isl_5_MT913014 | 13 | 3 | 1 | 17 |
| DNAS_isl_6_MT913011 | 8 | 4 | 1 | 13 |
| DNAS_isl_7_MT913008 | 6 | 1 | 1 | 8 |
| DNAS_isl_8_MT879649 | 7 | 3 | 1 | 11 |
| DNAS_isl_9_MT879651 | 9 | 2 | 1 | 12 |
| DNAS_isl_10_MT879653 | 9 | 2 | 1 | 12 |
| DNAS_isl_11_MT879654 | 11 | 3 | 1 | 15 |
| DNAS_isl_12_MT879652 | 10 | 3 | 1 | 14 |
| DNAS_isl_13_MT879645 | 5 | 9 | 1 | 15 |
| DNAS_isl_14_MT879658 | 7 | 7 | 1 | 15 |
| DNAS_isl_15_MT879648 | 6 | 2 | 1 | 9 |
| DNAS_isl_16_MT879656 | 8 | 2 | 1 | 11 |
| DNAS_isl_17_MT879647 | 8 | 1 | 1 | 10 |
| DNAS_isl_18_MT879646 | 6 | 1 | 1 | 8 |
| DNAS_isl_19_MT879650 | 7 | 1 | 1 | 9 |
| DNAS_isl_20_MT879659 | 10 | 1 | 1 | 12 |
| DNAS_isl_21_MT879655 | 6 | 2 | 1 | 9 |
| DNAS_isl_22_MT879657 | 11 | 4 | 1 | 16 |
| DNAS_isl_23_MT860694 | 11 | 4 | 1 | 16 |
| DNAS_isl_24_MT860691 | 4 | 4 | 1 | 9 |
| DNAS_isl_25_MT860689 | 9 | 3 | 1 | 13 |
| DNAS_isl_26_MT860679 | 4 | 6 | 3 | 13 |
| DNAS_isl_27_MT860681 | 10 | 3 | 1 | 14 |
| DNAS_isl_28_MT860688 | 8 | 3 | 1 | 12 |
| DNAS_isl_29_MT860690 | 15 | 2 | 1 | 18 |
| DNAS_isl_30_MT860680 | 10 | 2 | 1 | 13 |
| DNAS_isl_31_MT860684 | 8 | 4 | 1 | 13 |
| DNAS_isl_32_MT860683 | 6 | 4 | 1 | 11 |
| DNAS_isl_33_MT860687 | 8 | 1 | 1 | 10 |
| DNAS_isl_34_MT860693 | 8 | 6 | 1 | 15 |
| DNAS_isl_35_MT860692 | 6 | 5 | 1 | 12 |
| DNAS_isl_36_MT860685 | 9 | 1 | 1 | 11 |
| DNAS_isl_37_MT860682 | 12 | 3 | 1 | 16 |
| DNAS_isl_38_MT860686 | 7 | 4 | 1 | 12 |
| DNAS_isl_39_MT818584 | 6 | 2 | 1 | 9 |
| DNAS_isl_40_MT818590 | 6 | 8 | 1 | 15 |
| DNAS_isl_41_MT818583 | 7 | 4 | 1 | 12 |
| DNAS_isl_42_MT818586 | 13 | 2 | 1 | 16 |
| DNAS_isl_43_MT818585 | 13 | 2 | 1 | 16 |
| DNAS_isl_44_MT818581 | 6 | 1 | 1 | 8 |
| DNAS_isl_45_MT818589 | 10 | 4 | 1 | 15 |
| DNAS_isl_46_MT818587 | 11 | 3 | 1 | 15 |
| DNAS_isl_47_MT818591 | 12 | 3 | 1 | 16 |
| DNAS_isl_48_MT818582 | 7 | 2 | 1 | 10 |
| DNAS_isl_49_MT818588 | 8 | 4 | 1 | 13 |
| DNAS_isl_50_MT818580 | 7 | 1 | 1 | 9 |
| DNAS_isl_51_MT818579 | 11 | 4 | 1 | 16 |
| DNAS_isl_52_MT800893 | 10 | 3 | 1 | 14 |
| DNAS_isl_53_MT800884 | 6 | 1 | 1 | 8 |
| DNAS_isl_54_MT800876 | 7 | 3 | 1 | 11 |
| DNAS_isl_55_MT800881 | 8 | 2 | 1 | 11 |
| DNAS_isl_56_MT800882 | 9 | 1 | 1 | 11 |
| DNAS_isl_57_MT800888 | 8 | 3 | 1 | 12 |
| DNAS_isl_58_MT800886 | 11 | 2 | 1 | 14 |
| DNAS_isl_59_MT800889 | 8 | 1 | 1 | 10 |
| DNAS_isl_60_MT800887 | 9 | 3 | 1 | 13 |
| DNAS_isl_61_MT800875 | 10 | 2 | 1 | 13 |
| DNAS_isl_62_MT800894 | 10 | 4 | 1 | 15 |
| DNAS_isl_63_MT800892 | 7 | 5 | 1 | 13 |
| DNAS_isl_64_MT800891 | 6 | 5 | 1 | 12 |
| DNAS_isl_65_MT800877 | 9 | 2 | 1 | 12 |
| DNAS_isl_66_MT800879 | 7 | 3 | 1 | 11 |
| DNAS_isl_67_MT800880 | 7 | 2 | 1 | 10 |
| DNAS_isl_68_MT800878 | 10 | 1 | 1 | 12 |
| DNAS_isl_69_MT800885 | 8 | 3 | 1 | 12 |
| DNAS_isl_70_MT800890 | 6 | 2 | 1 | 9 |
| DNAS_isl_71_MT800883 | 6 | 5 | 1 | 12 |
| DNAS_isl_72_MT775562 | 7 | 2 | 1 | 10 |
| DNAS_isl_73_MT775565 | 9 | 2 | 1 | 12 |
| DNAS_isl_74_MT775564 | 6 | 2 | 1 | 9 |
| DNAS_isl_75_MT775558 | 8 | 3 | 1 | 12 |
| DNAS_isl_76_MT775563 | 7 | 2 | 1 | 10 |
| DNAS_isl_77_MT775571 | 9 | 1 | 1 | 11 |
| DNAS_isl_78_MT775570 | 9 | 1 | 1 | 11 |
| DNAS_isl_79_MT775568 | 10 | 2 | 1 | 13 |
| DNAS_isl_80_MT775560 | 7 | 3 | 1 | 11 |
| DNAS_isl_81_MT775566 | 6 | 1 | 1 | 8 |
| DNAS_isl_82_MT775567 | 8 | 5 | 2 | 15 |
| DNAS_isl_83_MT775572 | 12 | 2 | 1 | 15 |
| DNAS_isl_84_MT775569 | 8 | 3 | 1 | 12 |
| DNAS_isl_85_MT775559 | 7 | 4 | 1 | 12 |
| DNAS_isl_86_MT775561 | 8 | 5 | 1 | 14 |
| DNAS_isl_87_MT745761 | 9 | 2 | 1 | 12 |
| DNAS_isl_88_MT745755 | 7 | 5 | 1 | 13 |
| DNAS_isl_89_MT745754 | 9 | 3 | 1 | 13 |
| DNAS_isl_90_MT745756 | 7 | 3 | 1 | 11 |
| DNAS_isl_91_MT745765 | 9 | 5 | 1 | 15 |
| DNAS_isl_92_MT742762 | 9 | 5 | 1 | 15 |
| DNAS_isl_93_MT745757 | 9 | 5 | 1 | 15 |
| DNAS_isl_94_MT745760 | 6 | 1 | 1 | 8 |
| DNAS_isl_95_MT745763 | 6 | 1 | 1 | 8 |
| DNAS_isl_96_MT745753 | 7 | 2 | 1 | 10 |
| DNAS_isl_97_MT745752 | 9 | 3 | 1 | 13 |
| DNAS_isl_98_MT745764 | 9 | 1 | 1 | 11 |
| DNAS_isl_99_MT745759 | 6 | 2 | 1 | 9 |
| DNAS_isl_100_MT745762 | 6 | 2 | 1 | 9 |
| DNAS_isl_101_MT742761 | 8 | 3 | 1 | 12 |
| DNAS_isl_102_MT745750 | 9 | 2 | 1 | 12 |
| DNAS_isl_103_MT745751 | 10 | 3 | 1 | 14 |
| DNAS_isl_104_MT745758 | 10 | 1 | 1 | 12 |
| DNAS_isl_105_MT676415 | 6 | 4 | 1 | 11 |
| DNAS_isl_106_MT676414 | 6 | 1 | 1 | 8 |
| DNAS_isl_107_MT676413 | 6 | 1 | 1 | 8 |
| DNAS_isl_108_MT676418 | 5 | 3 | 1 | 9 |
| DNAS_isl_109_MT676420 | 6 | 2 | 1 | 9 |
| DNAS_isl_110_MT676416 | 8 | 2 | 1 | 11 |
| DNAS_isl_111_MT676412 | 6 | 2 | 1 | 9 |
| DNAS_isl_112_MT676419 | 7 | 2 | 1 | 10 |
| DNAS_isl_113_MT676421 | 9 | 2 | 1 | 12 |
| DNAS_isl_114_MT676417 | 6 | 1 | 1 | 8 |
| DNAS_isl_115_MT607976 | 6 | 4 | 1 | 11 |
| DNAS_isl_116_MT607972 | 8 | 3 | 1 | 12 |
| DNAS_isl_117_MT607975 | 9 | 2 | 1 | 12 |
| DNAS_isl_118_MT607974 | 7 | 6 | 1 | 14 |
| DNAS_isl_119_MT607973 | 6 | 3 | 1 | 10 |
| DNAS_isl_120_MT581414 | 9 | 1 | 0 | 10 |
| DNAS_isl_121_MT581434 | 5 | 4 | 1 | 10 |
| DNAS_isl_122_MT581413 | 6 | 4 | 1 | 11 |
| DNAS_isl_123_MT581425 | 12 | 2 | 1 | 15 |
| DNAS_isl_124_MT581422 | 9 | 2 | 1 | 12 |
| DNAS_isl_125_MT581423 | 8 | 2 | 1 | 11 |
| DNAS_isl_126_MT581412 | 7 | 1 | 0 | 8 |
| DNAS_isl_127_MT581433 | 7 | 4 | 1 | 12 |
| DNAS_isl_128_MT581432 | 6 | 4 | 1 | 11 |
| DNAS_isl_129_MT581427 | 8 | 2 | 1 | 11 |
| DNAS_isl_130_MT581428 | 6 | 2 | 1 | 9 |
| DNAS_isl_131_MT581436 | 7 | 3 | 1 | 11 |
| DNAS_isl_132_MT581435 | 7 | 4 | 1 | 12 |
| DNAS_isl_133_MT581426 | 8 | 2 | 1 | 11 |
| DNAS_isl_134_MT581424 | 11 | 2 | 1 | 14 |
| DNAS_isl_135_MT581421 | 8 | 2 | 1 | 11 |
| DNAS_isl_136_MT581417 | 2 | 3 | 1 | 6 |
| DNAS_isl_137_MT581419 | 2 | 4 | 1 | 7 |
| DNAS_isl_138_MT581416 | 2 | 3 | 1 | 6 |
| DNAS_isl_139_MT581418 | 3 | 4 | 1 | 8 |
| DNAS_isl_140_MT581411 | 7 | 2 | 1 | 10 |
| DNAS_isl_141_MT581430 | 6 | 2 | 1 | 9 |
| DNAS_isl_142_MT581431 | 4 | 4 | 1 | 9 |
| DNAS_isl_143_MT581429 | 5 | 2 | 1 | 8 |
| DNAS_isl_144_MT581420 | 8 | 2 | 1 | 11 |
| DNAS_isl_145_MT581415 | 4 | 3 | 1 | 8 |
| DNAS_isl_146_MT581410 | 6 | 2 | 1 | 9 |
| DNAS_isl_147_MT566438 | 10 | 2 | 1 | 13 |
| DNAS_isl_148_MT566437 | 2 | 3 | 1 | 6 |
| DNAS_isl_149_MT566436 | 9 | 3 | 1 | 13 |
| DNAS_isl_150_MT566435 | 9 | 3 | 1 | 13 |
| DNAS_isl_151_MT566434 | 11 | 2 | 1 | 14 |
| **Total Variants** | 1179 | 421 | 153 | 1753 |

**Figure 3:**
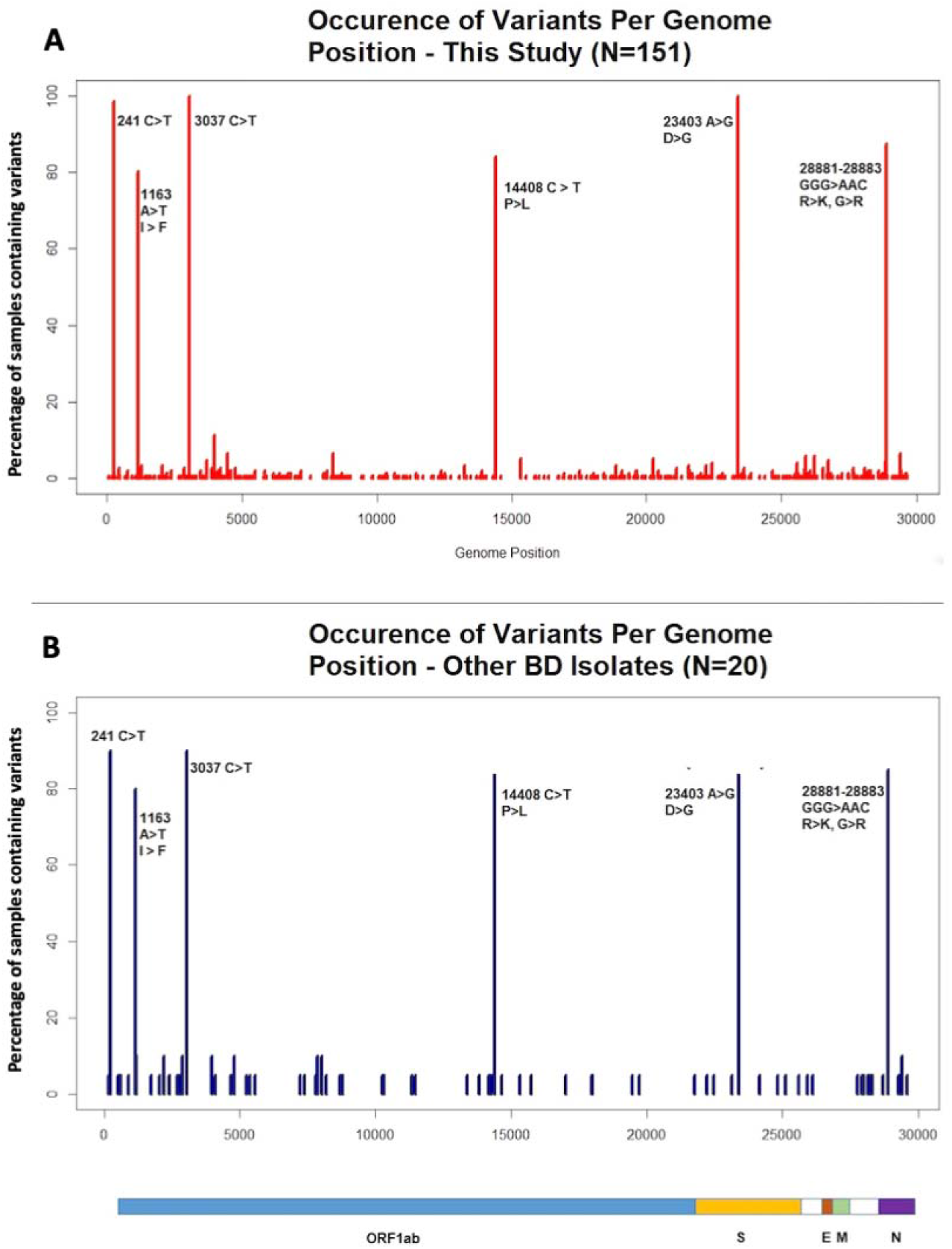
Distribution of variants across the SARS-CoV-2 Genome, as compared to the Wuhan Reference Sequence (NC_045512.2) A total of 1753 variants across 151 isolates spreading over 412 positions (3A in red). 8 of these variants occurred in over 120 isolates. These were the C to T change at 241, the A to T change at 1163, the C to T change at 3037 and at 14408, the A to G change at 23403, G to A change at 28881 and 28882, and finally the G to C change at 28883. The 241 C to T variant is the only one among 8 to occur in a non-coding region (it is a 5’ UTR SNP). The most common among the variants was the 23403 A to G change, which results in the D to G mutation at position 614 of the spike glycoprotein. The 8 major variants were also found in a number of other Bangladeshi isolates submitted in GISAID (3B in blue).

### 3.4 Gene Distribution of Variants

*ORF1ab* contained the highest number of mutations, which was expected considering- it is the largest among the SARS-CoV-2 genes. The nucleocapsid phosphoprotein encoded by *N* gene had the second highest number of mutations, followed by the surface glycoprotein encoded by the *S* gene. The remaining genes had fewer variants compared to these three. Lowest number of mutations was in *ORF6*, *ORF7b* and envelope protein encoded by *E* gene. Distribution of unique variants among the genes followed a similar pattern as the distribution of total variants. *ORF1ab* contained highest number of unique variants followed by *S* and *N* genes (**Table 2)**.

**Table 2:** Total gene variants observed in SARS-CoV-2 virus isolates. The *ORF1ab* contained highest number of mutations and is the longest among eleven (11) genes in SARS-CoV-2 virus. The nucleocapsid phosphoprotein encoded by *N* gene had the second highest number of mutations, followed by the surface glycoprotein encoded by the *S* gene.

| Gene | Total Variants | Unique Variants | Common Variants<br>(with other BD samples) |
| --- | --- | --- | --- |
| envelope protein | 2 | 2 | 0 |
| membrane glycoprotein | 23 | 12 | 0 |
| nucleocapsid phosphoprotein | 447 | 33 | 3 |
| ORF10 protein | 9 | 7 | 1 |
| ORF1ab polyprotein | 788 | 251 | 9 |
| ORF3a protein | 64 | 30 | 0 |
| ORF6 protein | 3 | 2 | 0 |
| ORF7a protein | 12 | 7 | 0 |
| ORF7b | 2 | 2 | 0 |
| ORF8 protein | 15 | 9 | 1 |
| surface glycoprotein | 235 | 53 | 3 |

The number of times each possible amino acid change occurred with a given gene was determined. **Figure 4** a heatmap displays frequency of occurrence for each type of change for all 11 genes, with a total of 76 types of amino acid changes were found in our analysis. Overall, D to G change was the most common (161 of 1175 amino acid variants), R to K (132), I to F (121), P to L (133) and G to R (132) were most frequent. Other common amino acid substitutions included A to V (15 for ORF1ab, ORF3a and S proteins), T to I (34), L to F (25), Q to H (28), and S to L (22). It should be noted that most amino acid changes are the same variant occurring in a large number of samples.

**Figure 4:**
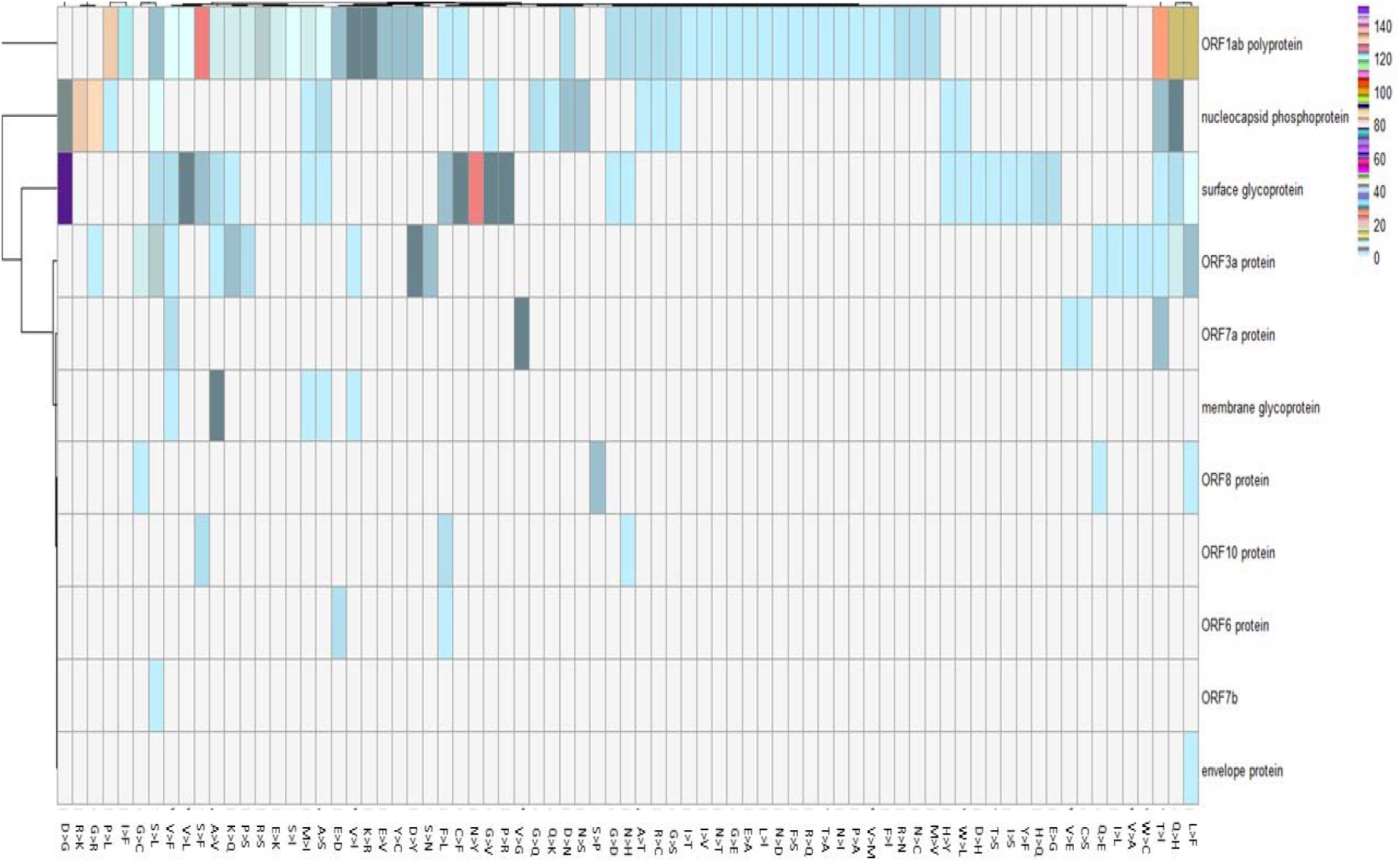
Heatmap showing the amino acid change per gene in our SARS-CoV-2 virus isolates. A total of 76 different kinds of amino acid changes could be observed across the 11 SARS-CoV-2 genes. Overall, D to G changes were the most abundant (161 out of 1175 amino acid variants). Aside from that, R to K (132 times), I to F (121 times), P to L (133 times) and G to R (132 times) occurred most frequently among the samples. Other common amino acid substitutions included A to V (occurring 15 times across the ORF1ab, ORF3a and S proteins), T to I (34 times), L to F (25 times), Q to H (28 times), and S to L (22 times). It should be noted that the majority of occurrences for each amino acid change are in fact the same variant occurring in a large number of samples. The majority of changes occurred only once, as indicated by the boxes with a light blue colour. As a result majority of these were only found in 1 of the genes. The white colour indicates that (i.e. the corresponding amino acid change for each of the white cells did not occur at all in the concerned gene).

### 3.5 Epidemiological sub-typing of Bangladeshi isolates

The isolates were classified into groups based on the EzCOVID19 SNP profile subtyping system as described in the methodology section^24^. The 151 isolates comprised eight groups based on type assignment according to the EzCOVID19 algorithm (**Figure 5**).Group 1 was most closely related to type 9. Both shared a common horizontal distance from the root and grouped together in the same branch. No mismatch was observed in the 41 SNP sites between this group and type 9. Nine virus isolates belonged to this group. Type 9 are most prevalent in Europe. Groups 2 and 3 were most closely related to type 2. Group 3 was identical to this type with regards to 41 SNP profile. However group 2 contained one SNP difference at position 14408. The base at this position was G, whereas for type 2 it was T. Three isolates belonged to group 2, while one belonged to group 3. This subtype is most prevalent in Europe, and found in parts of Asia, Africa and North America as shown is **Figure 5**. For group 4, the closest subtype was type 61. Only 1 fall into group 4 and was not an exact match with type 61, which contains the 11083 G to T variant and 27046 C to T variant. Both were absent in the group 4. It is worth noting that type 61 occurs exclusively in Europe. Study Group 5 closest related subtype was 15. The SNP profile was identical between these two. This type has a far more global distribution and occurs in North America, Europe and Asia. They group together with two member branches. Six belong to this study group. Group 6 shared highest similarity with type 26 with regards to the branch distance. Only one isolate joined group 6 and also has a SNP mismatch with type 26. Type 26 contained 25563 G to T variant, while our isolate did not. Isolates belonging to type 26 has been seen mostly in Europe and to a lesser degree, in South and North America. The closest matching subtype for Group 7 was type 64. Only one 1 isolate from our study belonged to study group 7 and it has contained a SNP mismatch with its most closely related type. Type 64 contained 11083 G to T variant, group 7 isolate did not. This is also a predominantly European subtype, while also occurring in Asia and South America. Lastly, group 8 shared closest common ancestor with type 4, with key SNP profiles exact match and comprised the majority of the SARS-CoV-2 isolates (130 of 151). Virus isolates belonging to type 4 are also predominantly in Europe; with limited presence in Asia, Oceania, and North America.

**Figure 5:**
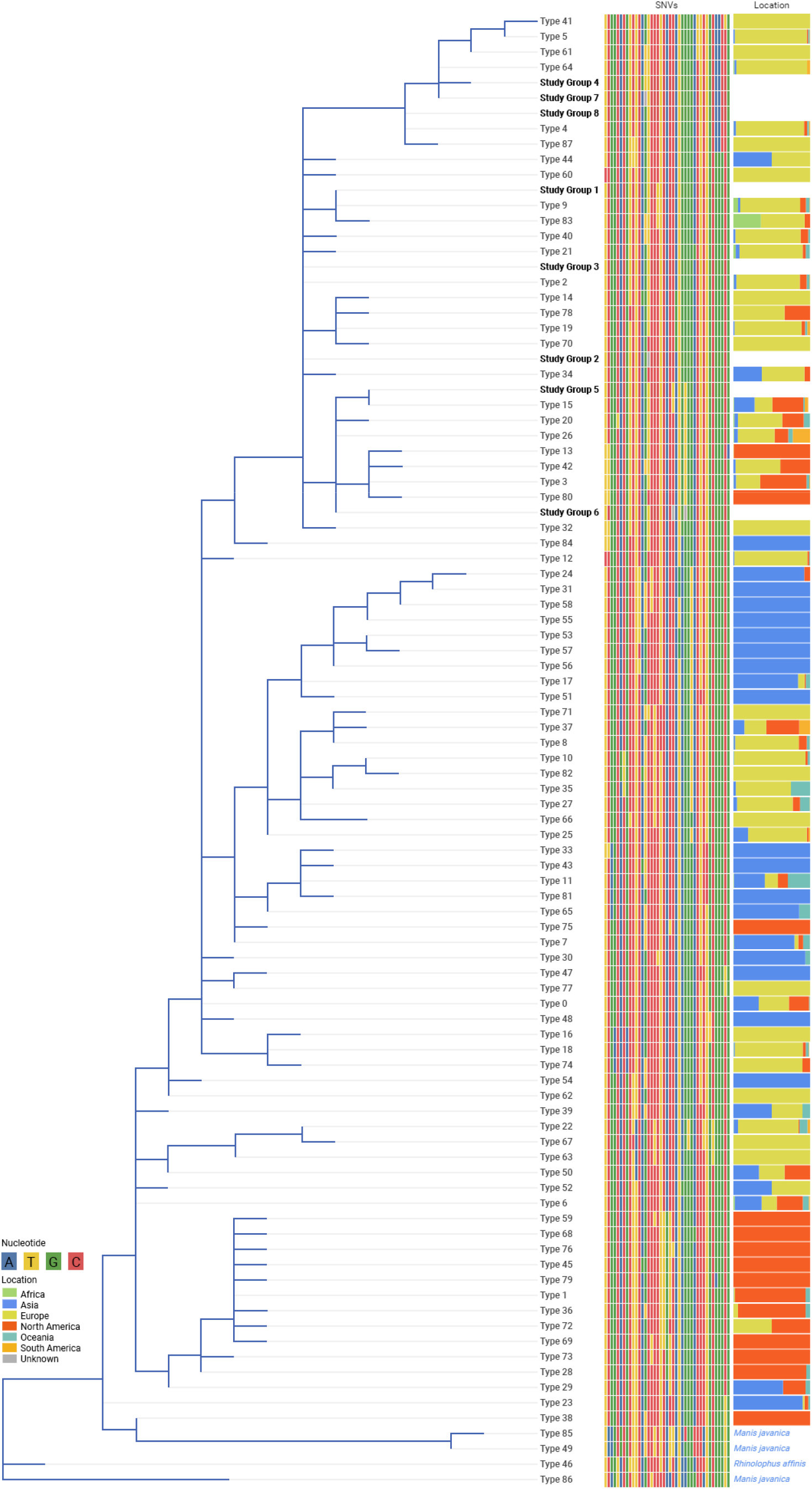
SNP based phylogenetic tree displaying the classification of 151 SARS-CoV-2 virus isolates according to EZCovid19 classification system for SARS-CoV-2 virus. Study group 1 (N=9) closely relates to type 9 with an exact match to SNP profile. Study group 2 (N=3) and Study group 3 (N=1) closely related to type 2. Study group 2 had a mismatch with type 2 at 14408 position (14408 T to G). Study Group 4 (N=1) closely matched with type 61 with two mismatch at 11083 G to T and 27046 C to T variant. Study Group 5 (N=6) has an exact match with type 15 in terms of branch distance and SNP profile. Study Group 6 (N=6) closely matched with Type 26 with a mismatch at 25563 G to T variant. Study Group 7 (N=1) has close match with type 64 with a SNP mismatch at 11083 G to T variant. Lastly, study group 8 (N=130) shared its closest common ancestor with type 4 with an exact match. All the close related types of our virus isolates has a string presence in Europe.

### 3.6 Clinical Importance of Variants

A total of 37 variants gave *p-*-values less than 0.05 selected parameters **(Supplementary Table 3)**, one of which was significantly associated with more than one disease symptoms, mainly 3961 C to T variant, shown an important determinant for patients developing sore throat and diarrhoea. The 14408 C to T, significantly associated with coughing, where individuals infected with a subtype with the variant appeared to suffer less from coughing. The particular variant was also linked with a host of other parameter determined usnig the random forest model. However, neither of the statistical test returned significant *p*-value for them. Variants 22199 G to T, 19593 C to T, 13902 T to C, 774 C to T, and 21597 C to T were significantly associated with development of sore throat, 28881 G to A, 28882 G to A, and 28883 G to C with chest pain, 21123 G to T with anorexia and 29118 C to T, 28178 G to T, 29262 G to T with pneumonia.

There were other variants which coincided with individuals lacking specific disease characteristics for example, 4105 G to T, 3456 A to G, 28305 A to G, 26051 G to A, 24685 T to C not loss of taste and smell.

Some variants, despite significant association with certain clinical factors, were not consistent with respect to specific factors. 28292 C to A, 4300 G to T, 26526 G to T, 17193 G to T, 2731 G to A, 98264 A to G, 12025 C to T, 8311 C to T, 714 G to A, 8366 G to A, 18859 G to T, 9416 G to A, 20808 G to A were all significantly related to onset or protection from skin rash. 28079 G to T and 3053 G to T were associated with itching and redness of eyes. Therefore, in all cases, the nature of the relationship could not be established based on the observations.

Finally, three variants were significantly associated with overall symptomatic status of the patients, namely 28580 G to A, 2363 C to T, and the 3871 G to T, and were significantly more likely to be asymptomatic. **Table 3** summarizes a few of these clinically relevant variants and their associated symptoms based on significance.

**Table 3:** Clinical Importance of Variants. A total of 37 variants returned p values less than 0.05 for various parameters. The 8 most significant variants is summarized here. 3961 C to T variant showed association with sore throat and diarrhoea development. The 14408 C to T inversely associated with cough development. The 28881 G to A, 28882 G to A, and 28883 G to C were associated with chest pain, and 29118 C to T, 28178 G to T, 29262 G to T were associated with pneumonia.

| Variant(s) | Variant Type | Disease State Association | Effect on Disease Phenotype | Fisher's Test P-value of Significance | Extra Comment |
| --- | --- | --- | --- | --- | --- |
| 14408 C to T | SNV | Cough | Protects from Cough | 0.02691376 |  |
| 3961C to T | SNV | Sore Throat | Causes Sore Throat | 0.000570637 |  |
|  | SNV | Diarrhoea | Protects from Diarrhoea | 0.03658523 |  |
| 28881G to A,<br>28882G to A,<br>28883G to C | SNV | Chest Pain | Causes Chest Pain | 0.02478889 | Co-variants |
| 29118C to T,<br>28178G to T,<br>29262G to T | Co-Variants (SNVs) | Pneumonia | Causes Pneumonia | 0.009615385 | Co-variants. Only 1 patient developed pneumonia however they were infected with a subtype containing unique variants. |

## 4. Discussion

In this large cross-sectional study, we have observed that SARS-CoV-2 virus isolates are related European subtypes, concurrent with other studies of Bangladeshi SARS-CoV-2 ^31^. SARS-CoV-2 virus isolated from neighbouring country, India, also were predominantly European sub-type^32^. This would suggest the virus entered Bangladesh via Europe, but multiple points of entry remain a very real possibility, reinforced by the observation that many of our isolates belonged to found in Africa, South America and North America. The GISAID clade classification offers a new perspective and most of our isolates belonged to clades that have recently seen high prevalence in Asia. However, possibilities that the virus may simply have entered these Asian countries through European sources earlier. Based on the data presented here, there is the possibility of SARS-CoV-2 entered Bangladesh through European countries, Asian countries, or both. A completely decisive conclusion is difficult to draw from SNP phylogeny. However, there is sufficient evidence for informed guess of European or Asian entry, with confidence inclining to the former.

The variants provided several referral points of insight. The eight most common variants were 241 C to T, 1163 A to T, 3037 C to T, 14408 C to T, 23403 G to A, 28881 G to A, 28882 G to A, and 28883 G to T, and were also present in the majority of other Bangladeshi isolates we included in our analysis. The 23403 G to A variant, which results in D to G substitution in the S protein, is one of the most prevalent amino acid mutations of the virus. In particular, it is a defining signature of the L, GH, G and GR clades. The majority of recently sequenced European isolates belonge to the latter three, lending further credibility to European origin claim since a large portion of recently sequenced Asian isolates have been classified as belonging to other clades (not the most common six).

Presence of an L clade member among the Bangladeshi isolates opens up an interesting perspective. The 28882 variant, with those at 241, 3037, and 23403 are characteristic markers for the L clade which contains the Wuhan reference genome and other early Chinese isolates. An evolutionary timeline would suggest one possible sequence would be SARS-CoV-2 entering Bangladesh via a Chinese source for example, several students had to be evacuated from Wuhan, arriving in Bangladesh midway into the outbreak. Hence, the aforementioned four variants in our samples. The virus responded to local selection pressures and incorporated the other common variants, such as 14408, 1163, and those variants at 28881 and 28883, which appear to have become co-variants. One reason why this may have sound more credible is the complete absence of a number of common European variants in our samples. Variants such as 14805 C to T, 1440 G to A, 2558 C to T and others were not present in any of our isolates, nor were they present in the other Bangladeshi samples analysed. Further 11083 G to T were also rare, occurring in only one or a few more samples. An issue with this logic could be that recently sequenced isolates from Europe have generally belonged to the G clade, i.e. containing variants at 241, 3037, and 23403 and not 11083, the other most common European variant of the clade typing scheme. This raises question whether the major route of transmission arose when the more recent European isolates arrived in Bangladesh. However, the possibility of virus subtypes in different parts of the world accumulating similar mutations independently must be considered.

A few of the amino acid variants showed clear prevalence compared to others. Six occurred in more than 80 of the isolates. The most common were 614 D to G for the S protein, which occurred in all samples, followed by, in decreasing order of frequency, 203 R to K and the 204 G to R also in N protein (132), P to L at 323 in RNA Polymerase of ORF1ab (127), and I to F in NSP2 of ORF1ab (121). The N protein R to K variant in particular is curious. One of the nucleotide variants, 28881 G to A, was an earlier variant been reported. It was found predominantly in European isolates, but nor two other co-variants, 28882 G to A and 28883 G to C. Subsequently, 28882 was detected and used as a marker in the GISAID’s clade classification. It’s perplexing when these three variants became associated (association is presumed since they occur together whenever they have been found in a genome). One possibility be that the 28881 and 28882 variants arose separately. The 28882 genotype perhaps accumulated the 28881 and 28883 variants, as it spread across Asia (28881 variant was not observed in Asia before). While the 28881 genotype isolate did the same and incorporated the 28882 and 28883 variants, completing the worldwide distribution pattern of these three variants that observed today ^33^. From the Bangladesh standpoint, both European and Asian origins remain a possibility, with regard to arrival of strains with this genotype. The European origin theory remains possible as other common European variants are largely absent, meaings less likely Europe to Bangladesh transmission. Finally, in one isolate there is a 28884 G to A variant eliminating the change of 204 G to R variants. Instead, the four nucleotide MNV leads to G to Q at 204 of the N protein. While seemingly similar, glutamine is not positively charged in its side chain, unlike arginine. Thus far it is indicated, that there is a lower mortality rate in Bangladesh compared many other parts of the world, the majority of the viral strains active in Bangladesh have to date harboured G to R change. Here an isolate incorporating a new mutation that, while not reversing the glycine to arginine substitution, does nonetheless alter it from a positively charged amino acid to one that is neutral (though polar).

There has not been significant focus this far on linking between individual SARS-CoV-2 variants and the disease manifestation associated with the virus. By employing machine learning and statistics, it was possible to report a significant association between variant and a specific clinical factors. Two variants, by using random forest model, were 14408 C to T and 3916 C to T. The second is a synonymous change. The first however results in a proline to leucine mutation in the RNA polymerase protein. Although it’s fisher’s test p value was only significant for the development of cough, the potential importance for overall symptomatic status deserves further attention. In general, individuals infected with the virus subtype containing this variant were asymptomatic. The RNA polymerase is critical for viral replication and alteration in function may well affect a function and consequently prevent onset of symptoms. In addition the 28881-28883 multiple nucleotide variant showed strong association with onset of chest pain. Heart failure (with other organ failures) has been suggested as a major causes of fatality in SARS-CoV-2^34^. These three variants appeared to be correlated with the onset of chest pain, suggesting a key pathogenic determinant.

## 5. Conclusion

Our goal in this study was to identify the phylogenetic origin and genetic variation among the SARS-CoV-2 isolates in Bangladesh. The findings indicate the 151 virus isolates from our study has strongest phylogenetic link with European isolates; although this link cannot be completely verified with this data alone. In addition, we identified 8 variants that are very common among the Bangladeshi variants considered in this study. These variants have been found in most parts of the world and have been validated by other studies carried out in Bangladesh. Finally, an association analysis between variants and clinical metadata showed one of the variants as being negatively correlated with the onset of cough, while another 3, an MNV, being positively correlated with the onset of chest pain among patients. This study is first large-scale genomic study to our knowledge revealing the phylogenetic relationship of SARS-CoV-2 virus circulating in Bangladesh which opens up new area of research to combat with this pandemic efficiently.

## 7. Author Contribution

MFR designed the experiment. AS, MBH, MHR identified the positive COVID-19 samples and MFR, MIK carried out the sequencing experiments. MNP and JN collected the metadata. SH, MC, KL, YOK and JC carried out the bioinformatics analysis. MIK and SH wrote the first draft manuscript. MFR, KNH, MMR, MK, MAK, NAH, RRC and SA reviewed the manuscript and provide comments. All authors approved the final submission.

## Notes

### Competing Interest Statement

The authors have declared no competing interest.

