## Supplemental Table 1 for "Large scale genomic and evolutionary study reveals SARS-CoV-2 virus isolates from Bangladesh strongly correlate with European origin and not with China"

**Supplementary Table 1: Summary of Isolate names and associated Gene Bank Accession Number**. We had sequenced in total one hundred and fifty one (151) SARS-CoV-2 virus isolated collected from Bangladeshi individuals. The virus isolates names and their respective Gene Bank Accession number are summarized here. All the sequences were uploaded under the NCBI Bio project# PRJNA643654.

| Isolate Name | Gene Bank Accession No | Isolate Name | Gene Bank Accession No | Isolate Name | Gene Bank Accession No |
| --- | --- | --- | --- | --- | --- |
| DNAS_isl_1_MT913010 | MT913010 | DNAS_isl_52_MT800893 | MT800893 | DNAS_isl_103_MT745751 | MT745751 |
| DNAS_isl_2_MT913012 | MT913012 | DNAS_isl_53_MT800884 | MT800884 | DNAS_isl_104_MT745758 | MT745758 |
| DNAS_isl_3_MT913013 | MT913013 | DNAS_isl_54_MT800876 | MT800876 | DNAS_isl_105_MT676415 | MT676415 |
| DNAS_isl_4_MT913009 | MT913009 | DNAS_isl_55_MT800881 | MT800881 | DNAS_isl_106_MT676414 | MT676414 |
| DNAS_isl_5_MT913014 | MT913014 | DNAS_isl_56_MT800882 | MT800882 | DNAS_isl_107_MT676413 | MT676413 |
| DNAS_isl_6_MT913011 | MT913011 | DNAS_isl_57_MT800888 | MT800888 | DNAS_isl_108_MT676418 | MT676418 |
| DNAS_isl_7_MT913008 | MT913008 | DNAS_isl_58_MT800886 | MT800886 | DNAS_isl_109_MT676420 | MT676420 |
| DNAS_isl_8_MT879649 | MT879649 | DNAS_isl_59_MT800889 | MT800889 | DNAS_isl_110_MT676416 | MT676416 |
| DNAS_isl_9_MT879651 | MT879651 | DNAS_isl_60_MT800887 | MT800887 | DNAS_isl_111_MT676412 | MT676412 |
| DNAS_isl_10_MT879653 | MT879653 | DNAS_isl_61_MT800875 | MT800875 | DNAS_isl_112_MT676419 | MT676419 |
| DNAS_isl_11_MT879654 | MT879654 | DNAS_isl_62_MT800894 | MT800894 | DNAS_isl_113_MT676421 | MT676421 |
| DNAS_isl_12_MT879652 | MT879652 | DNAS_isl_63_MT800892 | MT800892 | DNAS_isl_114_MT676417 | MT676417 |
| DNAS_isl_13_MT879645 | MT879645 | DNAS_isl_64_MT800891 | MT800891 | DNAS_isl_115_MT607976 | MT607976 |
| DNAS_isl_14_MT879658 | MT879658 | DNAS_isl_65_MT800877 | MT800877 | DNAS_isl_116_MT607972 | MT607972 |
| DNAS_isl_15_MT879648 | MT879648 | DNAS_isl_66_MT800879 | MT800879 | DNAS_isl_117_MT607975 | MT607975 |
| DNAS_isl_16_MT879656 | MT879656 | DNAS_isl_67_MT800880 | MT800880 | DNAS_isl_118_MT607974 | MT607974 |
| DNAS_isl_17_MT879647 | MT879647 | DNAS_isl_68_MT800878 | MT800878 | DNAS_isl_119_MT607973 | MT607973 |
| DNAS_isl_18_MT879646 | MT879646 | DNAS_isl_69_MT800885 | MT800885 | DNAS_isl_120_MT581414 | MT581414 |
| DNAS_isl_19_MT879650 | MT879650 | DNAS_isl_70_MT800890 | MT800890 | DNAS_isl_121_MT581434 | MT581434 |
| DNAS_isl_20_MT879659 | MT879659 | DNAS_isl_71_MT800883 | MT800883 | DNAS_isl_122_MT581413 | MT581413 |
| DNAS_isl_21_MT879655 | MT879655 | DNAS_isl_72_MT775562 | MT775562 | DNAS_isl_123_MT581425 | MT581425 |
| DNAS_isl_22_MT879657 | MT879657 | DNAS_isl_73_MT775565 | MT775565 | DNAS_isl_124_MT581422 | MT581422 |
| DNAS_isl_23_MT860694 | MT860694 | DNAS_isl_74_MT775564 | MT775564 | DNAS_isl_125_MT581423 | MT581423 |
| DNAS_isl_24_MT860691 | MT860691 | DNAS_isl_75_MT775558 | MT775558 | DNAS_isl_126_MT581412 | MT581412 |
| DNAS_isl_25_MT860689 | MT860689 | DNAS_isl_76_MT775563 | MT775563 | DNAS_isl_127_MT581433 | MT581433 |
| DNAS_isl_26_MT860679 | MT860679 | DNAS_isl_77_MT775571 | MT775571 | DNAS_isl_128_MT581432 | MT581432 |
| DNAS_isl_27_MT860681 | MT860681 | DNAS_isl_78_MT775570 | MT775570 | DNAS_isl_129_MT581427 | MT581427 |
| DNAS_isl_28_MT860688 | MT860688 | DNAS_isl_79_MT775568 | MT775568 | DNAS_isl_130_MT581428 | MT581428 |
| DNAS_isl_29_MT860690 | MT860690 | DNAS_isl_80_MT775560 | MT775560 | DNAS_isl_131_MT581436 | MT581436 |
| DNAS_isl_30_MT860680 | MT860680 | DNAS_isl_81_MT775566 | MT775566 | DNAS_isl_132_MT581435 | MT581435 |
| DNAS_isl_31_MT860684 | MT860684 | DNAS_isl_82_MT775567 | MT775567 | DNAS_isl_133_MT581426 | MT581426 |
| DNAS_isl_32_MT860683 | MT860683 | DNAS_isl_83_MT775572 | MT775572 | DNAS_isl_134_MT581424 | MT581424 |
| DNAS_isl_33_MT860687 | MT860687 | DNAS_isl_84_MT775569 | MT775569 | DNAS_isl_135_MT581421 | MT581421 |
| DNAS_isl_34_MT860693 | MT860693 | DNAS_isl_85_MT775559 | MT775559 | DNAS_isl_136_MT581417 | MT581417 |
| DNAS_isl_35_MT860692 | MT860692 | DNAS_isl_86_MT775561 | MT775561 | DNAS_isl_137_MT581419 | MT581419 |
| DNAS_isl_36_MT860685 | MT860685 | DNAS_isl_87_MT745761 | MT745761 | DNAS_isl_138_MT581416 | MT581416 |
| DNAS_isl_37_MT860682 | MT860682 | DNAS_isl_88_MT745755 | MT745755 | DNAS_isl_139_MT581418 | MT581418 |
| DNAS_isl_38_MT860686 | MT860686 | DNAS_isl_89_MT745754 | MT745754 | DNAS_isl_140_MT581411 | MT581411 |
| DNAS_isl_39_MT818584 | MT818584 | DNAS_isl_90_MT745756 | MT745756 | DNAS_isl_141_MT581430 | MT581430 |
| DNAS_isl_40_MT818590 | MT818590 | DNAS_isl_91_MT745765 | MT745765 | DNAS_isl_142_MT581431 | MT581431 |
| DNAS_isl_41_MT818583 | MT818583 | DNAS_isl_92_MT742762 | MT742762 | DNAS_isl_143_MT581429 | MT581429 |
| DNAS_isl_42_MT818586 | MT818586 | DNAS_isl_93_MT745757 | MT745757 | DNAS_isl_144_MT581420 | MT581420 |
| DNAS_isl_43_MT818585 | MT818585 | DNAS_isl_94_MT745760 | MT745760 | DNAS_isl_145_MT581415 | MT581415 |
| DNAS_isl_44_MT818581 | MT818581 | DNAS_isl_95_MT745763 | MT745763 | DNAS_isl_146_MT581410 | MT581410 |
| DNAS_isl_45_MT818589 | MT818589 | DNAS_isl_96_MT745753 | MT745753 | DNAS_isl_147_MT566438 | MT566438 |
| DNAS_isl_46_MT818587 | MT818587 | DNAS_isl_97_MT745752 | MT745752 | DNAS_isl_148_MT566437 | MT566437 |
| DNAS_isl_47_MT818591 | MT818591 | DNAS_isl_98_MT745764 | MT745764 | DNAS_isl_149_MT566436 | MT566436 |
| DNAS_isl_48_MT818582 | MT818582 | DNAS_isl_99_MT745759 | MT745759 | DNAS_isl_150_MT566435 | MT566435 |
| DNAS_isl_49_MT818588 | MT818588 | DNAS_isl_100_MT745762 | MT745762 | DNAS_isl_151_MT566434 | MT566434 |
| DNAS_isl_50_MT818580 | MT818580 | DNAS_isl_101_MT742761 | MT742761 |  |  |
| DNAS_isl_51_MT818579 | MT818579 | DNAS_isl_102_MT745750 | MT745750 |  |  |
