## Supplemental Table 2 for "Large scale genomic and evolutionary study reveals SARS-CoV-2 virus isolates from Bangladesh strongly correlate with European origin and not with China"

**Supplementary Table 2.: Summary of Single nucleotide variants and amino acid change among the 151 SARS-COV-2 virus isolates.**

All variants observed in individual isolates are summarized.

| **Isolate** | **Nucleotide Change** | **Reference amino acid** | **Query amino acid** | **Type** | **Coverage** |
| --- | --- | --- | --- | --- | --- |
| DNAS_isl_1_MT913010 | 23536CtoT | N | N | Synonymous | 8007 |
| DNAS_isl_1_MT913010 | 6352GtoT | K | N | Nonsynonymous | 2480 |
| DNAS_isl_1_MT913010 | 23403AtoG | D | G | Nonsynonymous | 7963 |
| DNAS_isl_1_MT913010 | 8026AtoG | A | A | Synonymous | 4205 |
| DNAS_isl_1_MT913010 | 28909CtoT | G | G | Synonymous | 7933 |
| DNAS_isl_1_MT913010 | 14408CtoT | P | L | Nonsynonymous | 1477 |
| DNAS_isl_1_MT913010 | 28883GtoC | G | R | Nonsynonymous | 2188 |
| DNAS_isl_1_MT913010 | 22118TtoC | F | L | Nonsynonymous | 4336 |
| DNAS_isl_1_MT913010 | 28882GtoA | R | K | Nonsynonymous | 2187 |
| DNAS_isl_1_MT913010 | 241CtoT | R | C | 5'UTR SNP | 3130 |
| DNAS_isl_1_MT913010 | 28881GtoA | R | K | Nonsynonymous | 2186 |
| DNAS_isl_1_MT913010 | 1163AtoT | I | F | Nonsynonymous | 7079 |
| DNAS_isl_1_MT913010 | 3037CtoT | F | F | Synonymous | 6030 |
| DNAS_isl_1_MT913010 | 203CtoT | S | L | 5'UTR SNP | 5928 |
| DNAS_isl_2_MT913012 | 3037CtoT | F | F | Synonymous | 5194 |
| DNAS_isl_2_MT913012 | 14408CtoT | P | L | Nonsynonymous | 5305 |
| DNAS_isl_2_MT913012 | 3163TtoC | S | S | Synonymous | 7926 |
| DNAS_isl_2_MT913012 | 5826CtoT | T | I | Nonsynonymous | 3108 |
| DNAS_isl_2_MT913012 | 23403AtoG | D | G | Nonsynonymous | 7934 |
| DNAS_isl_2_MT913012 | 241CtoT | R | C | 5'UTR SNP | 4087 |
| DNAS_isl_2_MT913012 | 17350GtoA | V | I | Nonsynonymous | 4564 |
| DNAS_isl_2_MT913012 | 28883GtoC | G | R | Nonsynonymous | 1606 |
| DNAS_isl_2_MT913012 | 11078TtoC | F | L | Nonsynonymous | 7981 |
| DNAS_isl_2_MT913012 | 28881GtoA | R | K | Nonsynonymous | 1593 |
| DNAS_isl_2_MT913012 | 28882GtoA | R | K | Nonsynonymous | 1601 |
| DNAS_isl_2_MT913012 | 29614CtoT | C | C | Synonymous | 7996 |
| DNAS_isl_3_MT913013 | 26051GtoA | S | N | Nonsynonymous | 2484 |
| DNAS_isl_3_MT913013 | 24685TtoC | D | D | Synonymous | 4295 |
| DNAS_isl_3_MT913013 | 28305AtoG | N | S | Nonsynonymous | 7981 |
| DNAS_isl_3_MT913013 | 26801CtoT | L | L | Synonymous | 7664 |
| DNAS_isl_3_MT913013 | 14408CtoT | P | L | Nonsynonymous | 2100 |
| DNAS_isl_3_MT913013 | 3871GtoT | K | N | Nonsynonymous | 2262 |
| DNAS_isl_3_MT913013 | 1163AtoT | I | F | Nonsynonymous | 7172 |
| DNAS_isl_3_MT913013 | 3456AtoG | Y | C | Nonsynonymous | 3012 |
| DNAS_isl_3_MT913013 | 27240GtoT | E | D | Nonsynonymous | 7337 |
| DNAS_isl_3_MT913013 | 3037CtoT | F | F | Synonymous | 5639 |
| DNAS_isl_3_MT913013 | 4105GtoT | K | N | Nonsynonymous | 5446 |
| DNAS_isl_3_MT913013 | 23403AtoG | D | G | Nonsynonymous | 7885 |
| DNAS_isl_3_MT913013 | 25855GtoT | D | Y | Nonsynonymous | 3807 |
| DNAS_isl_3_MT913013 | 28883GtoC | G | R | Nonsynonymous | 3860 |
| DNAS_isl_3_MT913013 | 241CtoT | R | C | 5'UTR SNP | 2802 |
| DNAS_isl_3_MT913013 | 28882GtoA | R | K | Nonsynonymous | 3860 |
| DNAS_isl_3_MT913013 | 28881GtoA | R | K | Nonsynonymous | 3852 |
| DNAS_isl_4_MT913009 | 19962GtoA | T | T | Synonymous | 6696 |
| DNAS_isl_4_MT913009 | 25855GtoT | D | Y | Nonsynonymous | 6697 |
| DNAS_isl_4_MT913009 | 241CtoT | R | C | 5'UTR SNP | 1302 |
| DNAS_isl_4_MT913009 | 28882GtoA | R | K | Nonsynonymous | 2406 |
| DNAS_isl_4_MT913009 | 1163AtoT | I | F | Nonsynonymous | 6498 |
| DNAS_isl_4_MT913009 | 28881GtoA | R | K | Nonsynonymous | 2403 |
| DNAS_isl_4_MT913009 | 23403AtoG | D | G | Nonsynonymous | 7961 |
| DNAS_isl_4_MT913009 | 14408CtoT | P | L | Nonsynonymous | 4303 |
| DNAS_isl_4_MT913009 | 3037CtoT | F | F | Synonymous | 5284 |
| DNAS_isl_4_MT913009 | 28883GtoC | G | R | Nonsynonymous | 2407 |
| DNAS_isl_4_MT913009 | 1563GtoT | C | F | Nonsynonymous | 7908 |
| DNAS_isl_5_MT913014 | 23403AtoG | D | G | Nonsynonymous | 7943 |
| DNAS_isl_5_MT913014 | 4105GtoT | K | N | Nonsynonymous | 7318 |
| DNAS_isl_5_MT913014 | 3037CtoT | F | F | Synonymous | 4807 |
| DNAS_isl_5_MT913014 | 14408CtoT | P | L | Nonsynonymous | 4324 |
| DNAS_isl_5_MT913014 | 241CtoT | R | C | 5'UTR SNP | 1032 |
| DNAS_isl_5_MT913014 | 28881GtoA | R | K | Nonsynonymous | 1726 |
| DNAS_isl_5_MT913014 | 25855GtoT | D | Y | Nonsynonymous | 2275 |
| DNAS_isl_5_MT913014 | 28882GtoA | R | K | Nonsynonymous | 1731 |
| DNAS_isl_5_MT913014 | 28883GtoC | G | R | Nonsynonymous | 1732 |
| DNAS_isl_5_MT913014 | 24685TtoC | D | D | Synonymous | 6328 |
| DNAS_isl_5_MT913014 | 26051GtoA | S | N | Nonsynonymous | 7520 |
| DNAS_isl_5_MT913014 | 27240GtoT | E | D | Nonsynonymous | 8001 |
| DNAS_isl_5_MT913014 | 3871GtoT | K | N | Nonsynonymous | 3151 |
| DNAS_isl_5_MT913014 | 28305AtoG | N | S | Nonsynonymous | 7979 |
| DNAS_isl_5_MT913014 | 1163AtoT | I | F | Nonsynonymous | 6519 |
| DNAS_isl_5_MT913014 | 3456AtoG | Y | C | Nonsynonymous | 5150 |
| DNAS_isl_5_MT913014 | 16377GtoT | P | P | Synonymous | 8000 |
| DNAS_isl_6_MT913011 | 3037CtoT | F | F | Synonymous | 3828 |
| DNAS_isl_6_MT913011 | 23403AtoG | D | G | Nonsynonymous | 5236 |
| DNAS_isl_6_MT913011 | 28882GtoA | R | K | Nonsynonymous | 4324 |
| DNAS_isl_6_MT913011 | 14408CtoT | P | L | Nonsynonymous | 1385 |
| DNAS_isl_6_MT913011 | 28881GtoA | R | K | Nonsynonymous | 4319 |
| DNAS_isl_6_MT913011 | 29245CtoT | V | V | Synonymous | 5807 |
| DNAS_isl_6_MT913011 | 8937CtoT | A | V | Nonsynonymous | 4321 |
| DNAS_isl_6_MT913011 | 241CtoT | R | C | 5'UTR SNP | 1319 |
| DNAS_isl_6_MT913011 | 28883GtoC | G | R | Nonsynonymous | 4325 |
| DNAS_isl_6_MT913011 | 26681CtoT | F | F | Synonymous | 5043 |
| DNAS_isl_6_MT913011 | 1163AtoT | I | F | Nonsynonymous | 6560 |
| DNAS_isl_6_MT913011 | 5051CtoT | P | S | Nonsynonymous | 1753 |
| DNAS_isl_6_MT913011 | 22432CtoT | D | D | Synonymous | 4276 |
| DNAS_isl_7_MT913008 | 28882GtoA | R | K | Nonsynonymous | 1522 |
| DNAS_isl_7_MT913008 | 241CtoT | R | C | 5'UTR SNP | 1212 |
| DNAS_isl_7_MT913008 | 28883GtoC | G | R | Nonsynonymous | 1523 |
| DNAS_isl_7_MT913008 | 28881GtoA | R | K | Nonsynonymous | 1518 |
| DNAS_isl_7_MT913008 | 3037CtoT | F | F | Synonymous | 5804 |
| DNAS_isl_7_MT913008 | 1163AtoT | I | F | Nonsynonymous | 5806 |
| DNAS_isl_7_MT913008 | 23403AtoG | D | G | Nonsynonymous | 7951 |
| DNAS_isl_7_MT913008 | 14408CtoT | P | L | Nonsynonymous | 5090 |
| DNAS_isl_8_MT879649 | 26882CtoT | L | L | Synonymous | 1580 |
| DNAS_isl_8_MT879649 | 10279CtoT | L | L | Synonymous | 2272 |
| DNAS_isl_8_MT879649 | 28883GtoC | G | R | Nonsynonymous | 1317 |
| DNAS_isl_8_MT879649 | 27752CtoT | T | I | Nonsynonymous | 963 |
| DNAS_isl_8_MT879649 | 23403AtoG | D | G | Nonsynonymous | 1908 |
| DNAS_isl_8_MT879649 | 241CtoT | R | C | 5'UTR SNP | 358 |
| DNAS_isl_8_MT879649 | 14408CtoT | P | L | Nonsynonymous | 362 |
| DNAS_isl_8_MT879649 | 3037CtoT | F | F | Synonymous | 1356 |
| DNAS_isl_8_MT879649 | 1163AtoT | I | F | Nonsynonymous | 1917 |
| DNAS_isl_8_MT879649 | 28881GtoA | R | K | Nonsynonymous | 1313 |
| DNAS_isl_8_MT879649 | 28882GtoA | R | K | Nonsynonymous | 1316 |
| DNAS_isl_9_MT879651 | 1163AtoT | I | F | Nonsynonymous | 7255 |
| DNAS_isl_9_MT879651 | 3037CtoT | F | F | Synonymous | 5827 |
| DNAS_isl_9_MT879651 | 14408CtoT | P | L | Nonsynonymous | 3766 |
| DNAS_isl_9_MT879651 | 6706CtoT | N | N | Synonymous | 7988 |
| DNAS_isl_9_MT879651 | 25904CtoT | S | L | Nonsynonymous | 3457 |
| DNAS_isl_9_MT879651 | 241CtoT | R | C | 5'UTR SNP | 2919 |
| DNAS_isl_9_MT879651 | 28883GtoC | G | R | Nonsynonymous | 3796 |
| DNAS_isl_9_MT879651 | 28882GtoA | R | K | Nonsynonymous | 3796 |
| DNAS_isl_9_MT879651 | 29160CtoT | T | I | Nonsynonymous | 7968 |
| DNAS_isl_9_MT879651 | 28881GtoA | R | K | Nonsynonymous | 3787 |
| DNAS_isl_9_MT879651 | 20537TtoC | I | T | Nonsynonymous | 7859 |
| DNAS_isl_9_MT879651 | 23403AtoG | D | G | Nonsynonymous | 7967 |
| DNAS_isl_10_MT879653 | 28882GtoA | R | K | Nonsynonymous | 2230 |
| DNAS_isl_10_MT879653 | 28883GtoC | G | R | Nonsynonymous | 2231 |
| DNAS_isl_10_MT879653 | 14408CtoT | P | L | Nonsynonymous | 1832 |
| DNAS_isl_10_MT879653 | 28881GtoA | R | K | Nonsynonymous | 2220 |
| DNAS_isl_10_MT879653 | 1163AtoT | I | F | Nonsynonymous | 6023 |
| DNAS_isl_10_MT879653 | 241CtoT | R | C | 5'UTR SNP | 3713 |
| DNAS_isl_10_MT879653 | 2363CtoT | L | F | Nonsynonymous | 2036 |
| DNAS_isl_10_MT879653 | 28580GtoA | D | N | Nonsynonymous | 7132 |
| DNAS_isl_10_MT879653 | 20238GtoT | R | S | Nonsynonymous | 4131 |
| DNAS_isl_10_MT879653 | 26211GtoT | V | V | Synonymous | 8009 |
| DNAS_isl_10_MT879653 | 3037CtoT | F | F | Synonymous | 6237 |
| DNAS_isl_10_MT879653 | 23403AtoG | D | G | Nonsynonymous | 7963 |
| DNAS_isl_11_MT879654 | 25904CtoT | S | L | Nonsynonymous | 6784 |
| DNAS_isl_11_MT879654 | 29578CtoA | F | L | Nonsynonymous | 4215 |
| DNAS_isl_11_MT879654 | 2363CtoT | L | F | Nonsynonymous | 230 |
| DNAS_isl_11_MT879654 | 1163AtoT | I | F | Nonsynonymous | 3319 |
| DNAS_isl_11_MT879654 | 3037CtoT | F | F | Synonymous | 7278 |
| DNAS_isl_11_MT879654 | 26211GtoT | V | V | Synonymous | 8011 |
| DNAS_isl_11_MT879654 | 28580GtoA | D | N | Nonsynonymous | 7120 |
| DNAS_isl_11_MT879654 | 20238GtoT | R | S | Nonsynonymous | 481 |
| DNAS_isl_11_MT879654 | 23403AtoG | D | G | Nonsynonymous | 7937 |
| DNAS_isl_11_MT879654 | 29274CtoT | T | I | Nonsynonymous | 2458 |
| DNAS_isl_11_MT879654 | 26828GtoT | L | L | Synonymous | 614 |
| DNAS_isl_11_MT879654 | 28881GtoA | R | K | Nonsynonymous | 5490 |
| DNAS_isl_11_MT879654 | 28882GtoA | R | K | Nonsynonymous | 5508 |
| DNAS_isl_11_MT879654 | 28883GtoC | G | R | Nonsynonymous | 5509 |
| DNAS_isl_11_MT879654 | 241CtoT | R | C | 5'UTR SNP | 2399 |
| DNAS_isl_12_MT879652 | 20238GtoT | R | S | Nonsynonymous | 613 |
| DNAS_isl_12_MT879652 | 26828GtoT | L | L | Synonymous | 691 |
| DNAS_isl_12_MT879652 | 26211GtoT | V | V | Synonymous | 1726 |
| DNAS_isl_12_MT879652 | 1163AtoT | I | F | Nonsynonymous | 617 |
| DNAS_isl_12_MT879652 | 3037CtoT | F | F | Synonymous | 790 |
| DNAS_isl_12_MT879652 | 28883GtoC | G | R | Nonsynonymous | 1072 |
| DNAS_isl_12_MT879652 | 14408CtoT | P | L | Nonsynonymous | 212 |
| DNAS_isl_12_MT879652 | 28882GtoA | R | K | Nonsynonymous | 1072 |
| DNAS_isl_12_MT879652 | 241CtoT | R | C | 5'UTR SNP | 272 |
| DNAS_isl_12_MT879652 | 28580GtoA | D | N | Nonsynonymous | 1395 |
| DNAS_isl_12_MT879652 | 23403AtoG | D | G | Nonsynonymous | 1056 |
| DNAS_isl_12_MT879652 | 28881GtoA | R | K | Nonsynonymous | 1071 |
| DNAS_isl_12_MT879652 | 29578CtoA | F | L | Nonsynonymous | 1719 |
| DNAS_isl_12_MT879652 | 2363CtoT | L | F | Nonsynonymous | 278 |
| DNAS_isl_13_MT879645 | 2836CtoT | C | C | Synonymous | 7819 |
| DNAS_isl_13_MT879645 | 22444CtoT | D | D | Synonymous | 5408 |
| DNAS_isl_13_MT879645 | 21000TtoC | D | D | Synonymous | 3984 |
| DNAS_isl_13_MT879645 | 23403AtoG | D | G | Nonsynonymous | 7973 |
| DNAS_isl_13_MT879645 | 14408CtoT | P | L | Nonsynonymous | 2806 |
| DNAS_isl_13_MT879645 | 241CtoT | R | C | 5'UTR SNP | 2903 |
| DNAS_isl_13_MT879645 | 13819CtoT | L | F | Nonsynonymous | 2423 |
| DNAS_isl_13_MT879645 | 16746TtoC | G | G | Synonymous | 5446 |
| DNAS_isl_13_MT879645 | 18189CtoA | T | T | Synonymous | 5837 |
| DNAS_isl_13_MT879645 | 28854CtoT | S | L | Nonsynonymous | 7785 |
| DNAS_isl_13_MT879645 | 18877CtoT | L | L | Synonymous | 7951 |
| DNAS_isl_13_MT879645 | 26735CtoT | Y | Y | Synonymous | 6729 |
| DNAS_isl_13_MT879645 | 3037CtoT | F | F | Synonymous | 5602 |
| DNAS_isl_13_MT879645 | 25563GtoT | Q | H | Nonsynonymous | 7987 |
| DNAS_isl_13_MT879645 | 3589CtoT | H | H | Synonymous | 1935 |
| DNAS_isl_14_MT879658 | 28882GtoA | R | K | Nonsynonymous | 4320 |
| DNAS_isl_14_MT879658 | 28883GtoC | G | R | Nonsynonymous | 4319 |
| DNAS_isl_14_MT879658 | 10357TtoC | Y | Y | Synonymous | 676 |
| DNAS_isl_14_MT879658 | 25522GtoA | G | R | Nonsynonymous | 5850 |
| DNAS_isl_14_MT879658 | 28881GtoA | R | K | Nonsynonymous | 4310 |
| DNAS_isl_14_MT879658 | 28079GtoT | V | V | Synonymous | 7875 |
| DNAS_isl_14_MT879658 | 2005CtoT | L | L | Synonymous | 2231 |
| DNAS_isl_14_MT879658 | 3961CtoT | I | I | Synonymous | 761 |
| DNAS_isl_14_MT879658 | 487GtoA | S | S | Synonymous | 3385 |
| DNAS_isl_14_MT879658 | 1163AtoT | I | F | Nonsynonymous | 2759 |
| DNAS_isl_14_MT879658 | 3037CtoT | F | F | Synonymous | 4614 |
| DNAS_isl_14_MT879658 | 241CtoT | R | C | 5'UTR SNP | 1036 |
| DNAS_isl_14_MT879658 | 1480CtoT | A | A | Synonymous | 5451 |
| DNAS_isl_14_MT879658 | 25450AtoC | I | L | Nonsynonymous | 3093 |
| DNAS_isl_14_MT879658 | 23403AtoG | D | G | Nonsynonymous | 2271 |
| DNAS_isl_15_MT879648 | 25617GtoT | K | N | Nonsynonymous | 6672 |
| DNAS_isl_15_MT879648 | 28882GtoA | R | K | Nonsynonymous | 6212 |
| DNAS_isl_15_MT879648 | 28881GtoA | R | K | Nonsynonymous | 6196 |
| DNAS_isl_15_MT879648 | 28883GtoC | G | R | Nonsynonymous | 6213 |
| DNAS_isl_15_MT879648 | 241CtoT | R | C | 5'UTR SNP | 2874 |
| DNAS_isl_15_MT879648 | 1909CtoT | F | F | Synonymous | 2260 |
| DNAS_isl_15_MT879648 | 3037CtoT | F | F | Synonymous | 7495 |
| DNAS_isl_15_MT879648 | 23403AtoG | D | G | Nonsynonymous | 7752 |
| DNAS_isl_15_MT879648 | 1163AtoT | I | F | Nonsynonymous | 4571 |
| DNAS_isl_16_MT879656 | 14408CtoT | P | L | Nonsynonymous | 80 |
| DNAS_isl_16_MT879656 | 241CtoT | R | C | 5'UTR SNP | 108 |
| DNAS_isl_16_MT879656 | 28881GtoA | R | K | Nonsynonymous | 380 |
| DNAS_isl_16_MT879656 | 28883GtoC | G | R | Nonsynonymous | 380 |
| DNAS_isl_16_MT879656 | 28882GtoA | R | K | Nonsynonymous | 380 |
| DNAS_isl_16_MT879656 | 1163AtoT | I | F | Nonsynonymous | 544 |
| DNAS_isl_16_MT879656 | 3037CtoT | F | F | Synonymous | 439 |
| DNAS_isl_16_MT879656 | 19813CtoT | P | S | Nonsynonymous | 142 |
| DNAS_isl_16_MT879656 | 23403AtoG | D | G | Nonsynonymous | 669 |
| DNAS_isl_16_MT879656 | 28727GtoA | A | T | Nonsynonymous | 1016 |
| DNAS_isl_16_MT879656 | 3961CtoT | I | I | Synonymous | 187 |
| DNAS_isl_17_MT879647 | 14408CtoT | P | L | Nonsynonymous | 2217 |
| DNAS_isl_17_MT879647 | 28882GtoA | R | K | Nonsynonymous | 3453 |
| DNAS_isl_17_MT879647 | 28883GtoC | G | R | Nonsynonymous | 3453 |
| DNAS_isl_17_MT879647 | 241CtoT | R | C | 5'UTR SNP | 3274 |
| DNAS_isl_17_MT879647 | 28881GtoA | R | K | Nonsynonymous | 3444 |
| DNAS_isl_17_MT879647 | 1163AtoT | I | F | Nonsynonymous | 7660 |
| DNAS_isl_17_MT879647 | 28326GtoT | G | V | Nonsynonymous | 7949 |
| DNAS_isl_17_MT879647 | 4503AtoT | E | V | Nonsynonymous | 4164 |
| DNAS_isl_17_MT879647 | 3037CtoT | F | F | Synonymous | 5944 |
| DNAS_isl_17_MT879647 | 23403AtoG | D | G | Nonsynonymous | 7968 |
| DNAS_isl_18_MT879646 | 28881GtoA | R | K | Nonsynonymous | 3535 |
| DNAS_isl_18_MT879646 | 29527GtoT | Q | H | Nonsynonymous | 7986 |
| DNAS_isl_18_MT879646 | 23403AtoG | D | G | Nonsynonymous | 7932 |
| DNAS_isl_18_MT879646 | 1163AtoT | I | F | Nonsynonymous | 7800 |
| DNAS_isl_18_MT879646 | 28882GtoA | R | K | Nonsynonymous | 3539 |
| DNAS_isl_18_MT879646 | 28883GtoC | G | R | Nonsynonymous | 3539 |
| DNAS_isl_18_MT879646 | 3037CtoT | F | F | Synonymous | 6592 |
| DNAS_isl_18_MT879646 | 241CtoT | R | C | 5'UTR SNP | 4308 |
| DNAS_isl_19_MT879650 | 1163AtoT | I | F | Nonsynonymous | 7769 |
| DNAS_isl_19_MT879650 | 28883GtoC | G | R | Nonsynonymous | 6197 |
| DNAS_isl_19_MT879650 | 28882GtoA | R | K | Nonsynonymous | 6197 |
| DNAS_isl_19_MT879650 | 23403AtoG | D | G | Nonsynonymous | 3655 |
| DNAS_isl_19_MT879650 | 28881GtoA | R | K | Nonsynonymous | 6188 |
| DNAS_isl_19_MT879650 | 3037CtoT | F | F | Synonymous | 7382 |
| DNAS_isl_19_MT879650 | 241CtoT | R | C | 5'UTR SNP | 1882 |
| DNAS_isl_19_MT879650 | 19543AtoG | I | V | Nonsynonymous | 5760 |
| DNAS_isl_19_MT879650 | 2210GtoT | V | F | Nonsynonymous | 3315 |
| DNAS_isl_20_MT879659 | 1163AtoT | I | F | Nonsynonymous | 4998 |
| DNAS_isl_20_MT879659 | 23403AtoG | D | G | Nonsynonymous | 6065 |
| DNAS_isl_20_MT879659 | 28883GtoC | G | R | Nonsynonymous | 3946 |
| DNAS_isl_20_MT879659 | 26730GtoT | V | F | Nonsynonymous | 3817 |
| DNAS_isl_20_MT879659 | 28882GtoA | R | K | Nonsynonymous | 3946 |
| DNAS_isl_20_MT879659 | 14408CtoT | P | L | Nonsynonymous | 751 |
| DNAS_isl_20_MT879659 | 28881GtoA | R | K | Nonsynonymous | 3935 |
| DNAS_isl_20_MT879659 | 241CtoT | R | C | 5'UTR SNP | 1731 |
| DNAS_isl_20_MT879659 | 6541CtoT | H | H | Synonymous | 2819 |
| DNAS_isl_20_MT879659 | 8760CtoT | A | V | Nonsynonymous | 1587 |
| DNAS_isl_20_MT879659 | 3037CtoT | F | F | Synonymous | 6727 |
| DNAS_isl_20_MT879659 | 12535TtoG | T | T | Synonymous | 3608 |
| DNAS_isl_21_MT879655 | 1163AtoT | I | F | Nonsynonymous | 7068 |
| DNAS_isl_21_MT879655 | 14408CtoT | P | L | Nonsynonymous | 2693 |
| DNAS_isl_21_MT879655 | 28882GtoA | R | K | Nonsynonymous | 1378 |
| DNAS_isl_21_MT879655 | 28881GtoA | R | K | Nonsynonymous | 1374 |
| DNAS_isl_21_MT879655 | 20703CtoT | Y | Y | Synonymous | 5996 |
| DNAS_isl_21_MT879655 | 241CtoT | R | C | 5'UTR SNP | 2588 |
| DNAS_isl_21_MT879655 | 28883GtoC | G | R | Nonsynonymous | 1378 |
| DNAS_isl_21_MT879655 | 3037CtoT | F | F | Synonymous | 5580 |
| DNAS_isl_21_MT879655 | 23403AtoG | D | G | Nonsynonymous | 7982 |
| DNAS_isl_22_MT879657 | 23403AtoG | D | G | Nonsynonymous | 7959 |
| DNAS_isl_22_MT879657 | 3037CtoT | F | F | Synonymous | 6012 |
| DNAS_isl_22_MT879657 | 241CtoT | R | C | 5'UTR SNP | 3859 |
| DNAS_isl_22_MT879657 | 9826AtoG | K | K | Synonymous | 2076 |
| DNAS_isl_22_MT879657 | 1163AtoT | I | F | Nonsynonymous | 7043 |
| DNAS_isl_22_MT879657 | 12025CtoT | S | S | Synonymous | 3872 |
| DNAS_isl_22_MT879657 | 8311CtoT | N | N | Synonymous | 7326 |
| DNAS_isl_22_MT879657 | 714GtoA | G | D | Nonsynonymous | 3275 |
| DNAS_isl_22_MT879657 | 28882GtoA | R | K | Nonsynonymous | 3133 |
| DNAS_isl_22_MT879657 | 8366GtoA | A | T | Nonsynonymous | 6654 |
| DNAS_isl_22_MT879657 | 18859GtoT | A | S | Nonsynonymous | 8003 |
| DNAS_isl_22_MT879657 | 14408CtoT | P | L | Nonsynonymous | 1906 |
| DNAS_isl_22_MT879657 | 28881GtoA | R | K | Nonsynonymous | 3121 |
| DNAS_isl_22_MT879657 | 9416GtoA | G | S | Nonsynonymous | 4454 |
| DNAS_isl_22_MT879657 | 28883GtoC | G | R | Nonsynonymous | 3133 |
| DNAS_isl_22_MT879657 | 20808GtoA | L | L | Synonymous | 7879 |
| DNAS_isl_23_MT860694 | 23401GtoT | Q | H | Nonsynonymous | 7995 |
| DNAS_isl_23_MT860694 | 23403AtoG | D | G | Nonsynonymous | 7986 |
| DNAS_isl_23_MT860694 | 8026AtoG | A | A | Synonymous | 4277 |
| DNAS_isl_23_MT860694 | 25455GtoT | K | N | Nonsynonymous | 7956 |
| DNAS_isl_23_MT860694 | 25777CtoT | L | F | Nonsynonymous | 7938 |
| DNAS_isl_23_MT860694 | 25904CtoT | S | L | Nonsynonymous | 5102 |
| DNAS_isl_23_MT860694 | 25521CtoT | F | F | Synonymous | 7846 |
| DNAS_isl_23_MT860694 | 28881GtoA | R | K | Nonsynonymous | 2408 |
| DNAS_isl_23_MT860694 | 241CtoT | R | C | 5'UTR SNP | 1076 |
| DNAS_isl_23_MT860694 | 28883GtoC | G | R | Nonsynonymous | 2412 |
| DNAS_isl_23_MT860694 | 28882GtoA | R | K | Nonsynonymous | 2411 |
| DNAS_isl_23_MT860694 | 3037CtoT | F | F | Synonymous | 5371 |
| DNAS_isl_23_MT860694 | 9502CtoT | A | A | Synonymous | 4305 |
| DNAS_isl_23_MT860694 | 1163AtoT | I | F | Nonsynonymous | 7034 |
| DNAS_isl_23_MT860694 | 27496TtoA | C | S | Nonsynonymous | 3821 |
| DNAS_isl_23_MT860694 | 14408CtoT | P | L | Nonsynonymous | 3654 |
| DNAS_isl_24_MT860691 | 13255CtoT | C | C | Synonymous | 2438 |
| DNAS_isl_24_MT860691 | 14408CtoT | P | L | Nonsynonymous | 2771 |
| DNAS_isl_24_MT860691 | 13458CtoT | S | L | Nonsynonymous | 1730 |
| DNAS_isl_24_MT860691 | 241CtoT | R | C | 5'UTR SNP | 2092 |
| DNAS_isl_24_MT860691 | 28854CtoT | S | L | Nonsynonymous | 7914 |
| DNAS_isl_24_MT860691 | 22747CtoT | V | V | Synonymous | 1436 |
| DNAS_isl_24_MT860691 | 26735CtoT | Y | Y | Synonymous | 6734 |
| DNAS_isl_24_MT860691 | 23403AtoG | D | G | Nonsynonymous | 7989 |
| DNAS_isl_24_MT860691 | 3037CtoT | F | F | Synonymous | 6623 |
| DNAS_isl_25_MT860689 | 9022TtoC | C | C | Synonymous | 2038 |
| DNAS_isl_25_MT860689 | 14408CtoT | P | L | Nonsynonymous | 832 |
| DNAS_isl_25_MT860689 | 12067GtoT | M | I | Nonsynonymous | 3393 |
| DNAS_isl_25_MT860689 | 28883GtoC | G | R | Nonsynonymous | 2594 |
| DNAS_isl_25_MT860689 | 10305AtoC | N | T | Nonsynonymous | 5187 |
| DNAS_isl_25_MT860689 | 241CtoT | R | C | 5'UTR SNP | 881 |
| DNAS_isl_25_MT860689 | 3871GtoT | K | N | Nonsynonymous | 1337 |
| DNAS_isl_25_MT860689 | 23403AtoG | D | G | Nonsynonymous | 4368 |
| DNAS_isl_25_MT860689 | 1163AtoT | I | F | Nonsynonymous | 3549 |
| DNAS_isl_25_MT860689 | 3037CtoT | F | F | Synonymous | 2969 |
| DNAS_isl_25_MT860689 | 28881GtoA | R | K | Nonsynonymous | 2581 |
| DNAS_isl_25_MT860689 | 28882GtoA | R | K | Nonsynonymous | 2594 |
| DNAS_isl_25_MT860689 | 24370CtoT | D | D | Synonymous | 772 |
| DNAS_isl_26_MT860679 | 3037CtoT | F | F | Synonymous | 5789 |
| DNAS_isl_26_MT860679 | 14408CtoT | P | L | Nonsynonymous | 2326 |
| DNAS_isl_26_MT860679 | 18828CtoT | V | V | Synonymous | 8003 |
| DNAS_isl_26_MT860679 | 16743TtoC | V | V | Synonymous | 4913 |
| DNAS_isl_26_MT860679 | 9856GtoT | V | V | Synonymous | 2515 |
| DNAS_isl_26_MT860679 | 28883GtoC | G | R | Nonsynonymous | 4184 |
| DNAS_isl_26_MT860679 | 28882GtoA | R | K | Nonsynonymous | 4184 |
| DNAS_isl_26_MT860679 | 1163AtoT | I | F | Nonsynonymous | 7778 |
| DNAS_isl_26_MT860679 | 3961CtoT | I | I | Synonymous | 3737 |
| DNAS_isl_26_MT860679 | 23403AtoG | D | G | Nonsynonymous | 7947 |
| DNAS_isl_26_MT860679 | 28881GtoA | R | K | Nonsynonymous | 4173 |
| DNAS_isl_26_MT860679 | 241CtoT | R | C | 5'UTR SNP | 2312 |
| DNAS_isl_26_MT860679 | 10816TtoC | S | S | Synonymous | 5865 |
| DNAS_isl_27_MT860681 | 199GtoT | V | F | 5'UTR SNP | 4730 |
| DNAS_isl_27_MT860681 | 28883GtoC | G | R | Nonsynonymous | 4252 |
| DNAS_isl_27_MT860681 | 61GtoT | V | F | 5'UTR SNP | 4675 |
| DNAS_isl_27_MT860681 | 4936GtoA | K | K | Synonymous | 3683 |
| DNAS_isl_27_MT860681 | 16750CtoT | P | S | Nonsynonymous | 3170 |
| DNAS_isl_27_MT860681 | 23403AtoG | D | G | Nonsynonymous | 6215 |
| DNAS_isl_27_MT860681 | 1163AtoT | I | F | Nonsynonymous | 6627 |
| DNAS_isl_27_MT860681 | 3037CtoT | F | F | Synonymous | 3767 |
| DNAS_isl_27_MT860681 | 18183CtoT | D | D | Synonymous | 4663 |
| DNAS_isl_27_MT860681 | 14408CtoT | P | L | Nonsynonymous | 1644 |
| DNAS_isl_27_MT860681 | 28882GtoA | R | K | Nonsynonymous | 4252 |
| DNAS_isl_27_MT860681 | 29621AtoC | N | H | Nonsynonymous | 7975 |
| DNAS_isl_27_MT860681 | 241CtoT | R | C | 5'UTR SNP | 1363 |
| DNAS_isl_27_MT860681 | 28881GtoA | R | K | Nonsynonymous | 4247 |
| DNAS_isl_28_MT860688 | 241CtoT | R | C | 5'UTR SNP | 1048 |
| DNAS_isl_28_MT860688 | 434GtoA | E | K | Nonsynonymous | 7609 |
| DNAS_isl_28_MT860688 | 3037CtoT | F | F | Synonymous | 4986 |
| DNAS_isl_28_MT860688 | 23403AtoG | D | G | Nonsynonymous | 7970 |
| DNAS_isl_28_MT860688 | 14408CtoT | P | L | Nonsynonymous | 3730 |
| DNAS_isl_28_MT860688 | 1163AtoT | I | F | Nonsynonymous | 6932 |
| DNAS_isl_28_MT860688 | 23604CtoG | P | R | Nonsynonymous | 7988 |
| DNAS_isl_28_MT860688 | 28881GtoA | R | K | Nonsynonymous | 2934 |
| DNAS_isl_28_MT860688 | 18981CtoT | H | H | Synonymous | 7318 |
| DNAS_isl_28_MT860688 | 28882GtoA | R | K | Nonsynonymous | 2941 |
| DNAS_isl_28_MT860688 | 28079GtoT | V | V | Synonymous | 7956 |
| DNAS_isl_28_MT860688 | 28883GtoC | G | R | Nonsynonymous | 2943 |
| DNAS_isl_29_MT860690 | 21614CtoT | L | F | Nonsynonymous | 6629 |
| DNAS_isl_29_MT860690 | 1408GtoA | E | K | Nonsynonymous | 7950 |
| DNAS_isl_29_MT860690 | 1163AtoT | I | F | Nonsynonymous | 7515 |
| DNAS_isl_29_MT860690 | 1406GtoA | E | K | Nonsynonymous | 7933 |
| DNAS_isl_29_MT860690 | 4105GtoT | K | N | Nonsynonymous | 6163 |
| DNAS_isl_29_MT860690 | 3456AtoG | Y | C | Nonsynonymous | 2321 |
| DNAS_isl_29_MT860690 | 28882GtoA | R | K | Nonsynonymous | 3118 |
| DNAS_isl_29_MT860690 | 28883GtoC | G | R | Nonsynonymous | 3118 |
| DNAS_isl_29_MT860690 | 26550GtoA | V | I | Nonsynonymous | 3224 |
| DNAS_isl_29_MT860690 | 241CtoT | R | C | 5'UTR SNP | 1542 |
| DNAS_isl_29_MT860690 | 3037CtoT | F | F | Synonymous | 5252 |
| DNAS_isl_29_MT860690 | 16943GtoT | S | I | Nonsynonymous | 7888 |
| DNAS_isl_29_MT860690 | 14408CtoT | P | L | Nonsynonymous | 4666 |
| DNAS_isl_29_MT860690 | 28881GtoA | R | K | Nonsynonymous | 3110 |
| DNAS_isl_29_MT860690 | 28305AtoG | N | S | Nonsynonymous | 7982 |
| DNAS_isl_29_MT860690 | 23403AtoG | D | G | Nonsynonymous | 7957 |
| DNAS_isl_29_MT860690 | 26051GtoA | S | N | Nonsynonymous | 6998 |
| DNAS_isl_29_MT860690 | 24685TtoC | D | D | Synonymous | 3966 |
| DNAS_isl_30_MT860680 | 3037CtoT | F | F | Synonymous | 6187 |
| DNAS_isl_30_MT860680 | 14408CtoT | P | L | Nonsynonymous | 2087 |
| DNAS_isl_30_MT860680 | 1163AtoT | I | F | Nonsynonymous | 6634 |
| DNAS_isl_30_MT860680 | 241CtoT | R | C | 5'UTR SNP | 2422 |
| DNAS_isl_30_MT860680 | 23403AtoG | D | G | Nonsynonymous | 7956 |
| DNAS_isl_30_MT860680 | 28883GtoC | G | R | Nonsynonymous | 4337 |
| DNAS_isl_30_MT860680 | 20238GtoT | R | S | Nonsynonymous | 3478 |
| DNAS_isl_30_MT860680 | 5886CtoT | T | I | Nonsynonymous | 3514 |
| DNAS_isl_30_MT860680 | 28881GtoA | R | K | Nonsynonymous | 4327 |
| DNAS_isl_30_MT860680 | 26211GtoT | V | V | Synonymous | 8009 |
| DNAS_isl_30_MT860680 | 25904CtoT | S | L | Nonsynonymous | 2213 |
| DNAS_isl_30_MT860680 | 28882GtoA | R | K | Nonsynonymous | 4337 |
| DNAS_isl_30_MT860680 | 27205TtoC | F | L | Nonsynonymous | 2149 |
| DNAS_isl_31_MT860684 | 23403AtoG | D | G | Nonsynonymous | 7977 |
| DNAS_isl_31_MT860684 | 4456CtoT | A | A | Synonymous | 7339 |
| DNAS_isl_31_MT860684 | 241CtoT | R | C | 5'UTR SNP | 3787 |
| DNAS_isl_31_MT860684 | 1163AtoT | I | F | Nonsynonymous | 7559 |
| DNAS_isl_31_MT860684 | 27476CtoT | T | I | Nonsynonymous | 2307 |
| DNAS_isl_31_MT860684 | 24782AtoC | N | H | Nonsynonymous | 5842 |
| DNAS_isl_31_MT860684 | 3961CtoT | I | I | Synonymous | 5236 |
| DNAS_isl_31_MT860684 | 3037CtoT | F | F | Synonymous | 5420 |
| DNAS_isl_31_MT860684 | 28881GtoA | R | K | Nonsynonymous | 4132 |
| DNAS_isl_31_MT860684 | 19374CtoT | F | F | Synonymous | 4813 |
| DNAS_isl_31_MT860684 | 28883GtoC | G | R | Nonsynonymous | 4136 |
| DNAS_isl_31_MT860684 | 14408CtoT | P | L | Nonsynonymous | 2464 |
| DNAS_isl_31_MT860684 | 28882GtoA | R | K | Nonsynonymous | 4136 |
| DNAS_isl_32_MT860683 | 6616AtoG | L | L | Synonymous | 2857 |
| DNAS_isl_32_MT860683 | 3037CtoT | F | F | Synonymous | 5517 |
| DNAS_isl_32_MT860683 | 1163AtoT | I | F | Nonsynonymous | 4368 |
| DNAS_isl_32_MT860683 | 3961CtoT | I | I | Synonymous | 4060 |
| DNAS_isl_32_MT860683 | 241CtoT | R | C | 5'UTR SNP | 1243 |
| DNAS_isl_32_MT860683 | 14408CtoT | P | L | Nonsynonymous | 2305 |
| DNAS_isl_32_MT860683 | 23403AtoG | D | G | Nonsynonymous | 6746 |
| DNAS_isl_32_MT860683 | 28883GtoC | G | R | Nonsynonymous | 5085 |
| DNAS_isl_32_MT860683 | 28881GtoA | R | K | Nonsynonymous | 5078 |
| DNAS_isl_32_MT860683 | 28882GtoA | R | K | Nonsynonymous | 5085 |
| DNAS_isl_32_MT860683 | 18495TtoC | L | L | Synonymous | 749 |
| DNAS_isl_33_MT860687 | 241CtoT | R | C | 5'UTR SNP | 1555 |
| DNAS_isl_33_MT860687 | 23403AtoG | D | G | Nonsynonymous | 7997 |
| DNAS_isl_33_MT860687 | 2351GtoA | G | S | Nonsynonymous | 7964 |
| DNAS_isl_33_MT860687 | 14408CtoT | P | L | Nonsynonymous | 3623 |
| DNAS_isl_33_MT860687 | 25555GtoT | V | F | Nonsynonymous | 6451 |
| DNAS_isl_33_MT860687 | 28881GtoA | R | K | Nonsynonymous | 3112 |
| DNAS_isl_33_MT860687 | 1163AtoT | I | F | Nonsynonymous | 6700 |
| DNAS_isl_33_MT860687 | 28882GtoA | R | K | Nonsynonymous | 3120 |
| DNAS_isl_33_MT860687 | 28883GtoC | G | R | Nonsynonymous | 3120 |
| DNAS_isl_33_MT860687 | 3037CtoT | F | F | Synonymous | 4506 |
| DNAS_isl_34_MT860693 | 3037CtoT | F | F | Synonymous | 5041 |
| DNAS_isl_34_MT860693 | 25494GtoT | T | T | Synonymous | 4484 |
| DNAS_isl_34_MT860693 | 26735CtoT | Y | Y | Synonymous | 7941 |
| DNAS_isl_34_MT860693 | 1068GtoA | G | E | Nonsynonymous | 7068 |
| DNAS_isl_34_MT860693 | 25563GtoT | Q | H | Nonsynonymous | 7971 |
| DNAS_isl_34_MT860693 | 23403AtoG | D | G | Nonsynonymous | 7956 |
| DNAS_isl_34_MT860693 | 28854CtoT | S | L | Nonsynonymous | 7967 |
| DNAS_isl_34_MT860693 | 241CtoT | R | C | 5'UTR SNP | 1465 |
| DNAS_isl_34_MT860693 | 27092CtoT | D | D | Synonymous | 7990 |
| DNAS_isl_34_MT860693 | 2940CtoT | P | L | Nonsynonymous | 3838 |
| DNAS_isl_34_MT860693 | 14408CtoT | P | L | Nonsynonymous | 3697 |
| DNAS_isl_34_MT860693 | 22444CtoT | D | D | Synonymous | 7902 |
| DNAS_isl_34_MT860693 | 25273GtoC | M | I | Nonsynonymous | 8002 |
| DNAS_isl_34_MT860693 | 1593CtoT | S | F | Nonsynonymous | 6391 |
| DNAS_isl_34_MT860693 | 18877CtoT | L | L | Synonymous | 7978 |
| DNAS_isl_35_MT860692 | 14055GtoT | L | L | Synonymous | 6113 |
| DNAS_isl_35_MT860692 | 26735CtoT | Y | Y | Synonymous | 7888 |
| DNAS_isl_35_MT860692 | 23403AtoG | D | G | Nonsynonymous | 7959 |
| DNAS_isl_35_MT860692 | 25563GtoT | Q | H | Nonsynonymous | 7976 |
| DNAS_isl_35_MT860692 | 28854CtoT | S | L | Nonsynonymous | 7978 |
| DNAS_isl_35_MT860692 | 241CtoT | R | C | 5'UTR SNP | 4147 |
| DNAS_isl_35_MT860692 | 3037CtoT | F | F | Synonymous | 5581 |
| DNAS_isl_35_MT860692 | 14408CtoT | P | L | Nonsynonymous | 3853 |
| DNAS_isl_35_MT860692 | 22444CtoT | D | D | Synonymous | 7350 |
| DNAS_isl_35_MT860692 | 25494GtoT | T | T | Synonymous | 5165 |
| DNAS_isl_35_MT860692 | 21304CtoA | R | N | Nonsynonymous | 4248 |
| DNAS_isl_35_MT860692 | 21305GtoA | R | N | Nonsynonymous | 4268 |
| DNAS_isl_36_MT860685 | 5794AtoT | E | D | Nonsynonymous | 2880 |
| DNAS_isl_36_MT860685 | 3037CtoT | F | F | Synonymous | 2942 |
| DNAS_isl_36_MT860685 | 241CtoT | R | C | 5'UTR SNP | 806 |
| DNAS_isl_36_MT860685 | 1163AtoT | I | F | Nonsynonymous | 4182 |
| DNAS_isl_36_MT860685 | 14408CtoT | P | L | Nonsynonymous | 695 |
| DNAS_isl_36_MT860685 | 4503AtoT | E | V | Nonsynonymous | 488 |
| DNAS_isl_36_MT860685 | 23403AtoG | D | G | Nonsynonymous | 3756 |
| DNAS_isl_36_MT860685 | 28883GtoC | G | R | Nonsynonymous | 3395 |
| DNAS_isl_36_MT860685 | 28882GtoA | R | K | Nonsynonymous | 3395 |
| DNAS_isl_36_MT860685 | 28881GtoA | R | K | Nonsynonymous | 3383 |
| DNAS_isl_36_MT860685 | 6633CtoT | A | V | Nonsynonymous | 2142 |
| DNAS_isl_37_MT860682 | 26305CtoT | L | F | Nonsynonymous | 5748 |
| DNAS_isl_37_MT860682 | 241CtoT | R | C | 5'UTR SNP | 3328 |
| DNAS_isl_37_MT860682 | 430AtoC | E | D | Nonsynonymous | 7983 |
| DNAS_isl_37_MT860682 | 3961CtoT | I | I | Synonymous | 3411 |
| DNAS_isl_37_MT860682 | 3037CtoT | F | F | Synonymous | 5801 |
| DNAS_isl_37_MT860682 | 1163AtoT | I | F | Nonsynonymous | 7218 |
| DNAS_isl_37_MT860682 | 17790GtoT | Q | H | Nonsynonymous | 7320 |
| DNAS_isl_37_MT860682 | 14408CtoT | P | L | Nonsynonymous | 2102 |
| DNAS_isl_37_MT860682 | 11235CtoT | A | V | Nonsynonymous | 4776 |
| DNAS_isl_37_MT860682 | 6701CtoT | L | F | Nonsynonymous | 4873 |
| DNAS_isl_37_MT860682 | 10396GtoT | V | V | Synonymous | 7685 |
| DNAS_isl_37_MT860682 | 28883GtoC | G | R | Nonsynonymous | 4067 |
| DNAS_isl_37_MT860682 | 28882GtoA | R | K | Nonsynonymous | 4066 |
| DNAS_isl_37_MT860682 | 28881GtoA | R | K | Nonsynonymous | 4052 |
| DNAS_isl_37_MT860682 | 13966GtoA | A | T | Nonsynonymous | 3401 |
| DNAS_isl_37_MT860682 | 23403AtoG | D | G | Nonsynonymous | 7959 |
| DNAS_isl_38_MT860686 | 16014GtoT | R | R | Synonymous | 1477 |
| DNAS_isl_38_MT860686 | 23403AtoG | D | G | Nonsynonymous | 3172 |
| DNAS_isl_38_MT860686 | 1163AtoT | I | F | Nonsynonymous | 3764 |
| DNAS_isl_38_MT860686 | 28883GtoC | G | R | Nonsynonymous | 3015 |
| DNAS_isl_38_MT860686 | 28881GtoA | R | K | Nonsynonymous | 3003 |
| DNAS_isl_38_MT860686 | 28882GtoA | R | K | Nonsynonymous | 3015 |
| DNAS_isl_38_MT860686 | 28076CtoT | C | C | Synonymous | 7780 |
| DNAS_isl_38_MT860686 | 3037CtoT | F | F | Synonymous | 2435 |
| DNAS_isl_38_MT860686 | 13060TtoC | T | T | Synonymous | 1416 |
| DNAS_isl_38_MT860686 | 18105GtoT | Q | H | Nonsynonymous | 3880 |
| DNAS_isl_38_MT860686 | 241CtoT | R | C | 5'UTR SNP | 812 |
| DNAS_isl_38_MT860686 | 6990CtoT | S | F | Nonsynonymous | 1612 |
| DNAS_isl_39_MT818584 | 28883GtoC | G | R | Nonsynonymous | 4532 |
| DNAS_isl_39_MT818584 | 28882GtoA | R | K | Nonsynonymous | 4532 |
| DNAS_isl_39_MT818584 | 23403AtoG | D | G | Nonsynonymous | 7931 |
| DNAS_isl_39_MT818584 | 14408CtoT | P | L | Nonsynonymous | 1259 |
| DNAS_isl_39_MT818584 | 1163AtoT | I | F | Nonsynonymous | 7513 |
| DNAS_isl_39_MT818584 | 241CtoT | R | C | 5'UTR SNP | 3199 |
| DNAS_isl_39_MT818584 | 28881GtoA | R | K | Nonsynonymous | 4518 |
| DNAS_isl_39_MT818584 | 23797TtoC | D | D | Synonymous | 2905 |
| DNAS_isl_39_MT818584 | 3037CtoT | F | F | Synonymous | 6291 |
| DNAS_isl_40_MT818590 | 16721AtoC | E | A | Nonsynonymous | 7934 |
| DNAS_isl_40_MT818590 | 22444CtoT | D | D | Synonymous | 5143 |
| DNAS_isl_40_MT818590 | 18877CtoT | L | L | Synonymous | 7970 |
| DNAS_isl_40_MT818590 | 3037CtoT | F | F | Synonymous | 6180 |
| DNAS_isl_40_MT818590 | 2836CtoT | C | C | Synonymous | 5385 |
| DNAS_isl_40_MT818590 | 25563GtoT | Q | H | Nonsynonymous | 7990 |
| DNAS_isl_40_MT818590 | 28854CtoT | S | L | Nonsynonymous | 7958 |
| DNAS_isl_40_MT818590 | 23403AtoG | D | G | Nonsynonymous | 7954 |
| DNAS_isl_40_MT818590 | 12514GtoT | T | T | Synonymous | 5051 |
| DNAS_isl_40_MT818590 | 241CtoT | R | C | 5'UTR SNP | 3124 |
| DNAS_isl_40_MT818590 | 29353CtoT | Y | Y | Synonymous | 7944 |
| DNAS_isl_40_MT818590 | 26164CtoT | P | S | Nonsynonymous | 4046 |
| DNAS_isl_40_MT818590 | 14408CtoT | P | L | Nonsynonymous | 1639 |
| DNAS_isl_40_MT818590 | 26735CtoT | Y | Y | Synonymous | 6547 |
| DNAS_isl_40_MT818590 | 6967AtoG | L | L | Synonymous | 4334 |
| DNAS_isl_41_MT818583 | 241CtoT | R | C | 5'UTR SNP | 1157 |
| DNAS_isl_41_MT818583 | 28882GtoA | R | K | Nonsynonymous | 4693 |
| DNAS_isl_41_MT818583 | 28883GtoC | G | R | Nonsynonymous | 4694 |
| DNAS_isl_41_MT818583 | 25842TtoC | H | H | Synonymous | 2126 |
| DNAS_isl_41_MT818583 | 2334CtoT | A | V | Nonsynonymous | 2412 |
| DNAS_isl_41_MT818583 | 8956CtoT | Y | Y | Synonymous | 3345 |
| DNAS_isl_41_MT818583 | 3961CtoT | I | I | Synonymous | 1624 |
| DNAS_isl_41_MT818583 | 3037CtoT | F | F | Synonymous | 3342 |
| DNAS_isl_41_MT818583 | 1163AtoT | I | F | Nonsynonymous | 4139 |
| DNAS_isl_41_MT818583 | 23403AtoG | D | G | Nonsynonymous | 3515 |
| DNAS_isl_41_MT818583 | 28881GtoA | R | K | Nonsynonymous | 4680 |
| DNAS_isl_41_MT818583 | 14408CtoT | P | L | Nonsynonymous | 545 |
| DNAS_isl_42_MT818586 | 28883GtoC | G | R | Nonsynonymous | 4092 |
| DNAS_isl_42_MT818586 | 28882GtoA | R | K | Nonsynonymous | 4092 |
| DNAS_isl_42_MT818586 | 29625CtoT | S | F | Nonsynonymous | 7906 |
| DNAS_isl_42_MT818586 | 23403AtoG | D | G | Nonsynonymous | 7921 |
| DNAS_isl_42_MT818586 | 241CtoT | R | C | 5'UTR SNP | 3594 |
| DNAS_isl_42_MT818586 | 23520CtoT | A | V | Nonsynonymous | 7997 |
| DNAS_isl_42_MT818586 | 28881GtoA | R | K | Nonsynonymous | 4081 |
| DNAS_isl_42_MT818586 | 11462AtoG | M | V | Nonsynonymous | 6982 |
| DNAS_isl_42_MT818586 | 3037CtoT | F | F | Synonymous | 4971 |
| DNAS_isl_42_MT818586 | 25947GtoT | Q | H | Nonsynonymous | 6891 |
| DNAS_isl_42_MT818586 | 5826CtoT | T | I | Nonsynonymous | 2574 |
| DNAS_isl_42_MT818586 | 1163AtoT | I | F | Nonsynonymous | 7502 |
| DNAS_isl_42_MT818586 | 10369CtoT | R | R | Synonymous | 3757 |
| DNAS_isl_42_MT818586 | 14408CtoT | P | L | Nonsynonymous | 2829 |
| DNAS_isl_42_MT818586 | 10658GtoA | V | I | Nonsynonymous | 6180 |
| DNAS_isl_42_MT818586 | 18040GtoT | A | S | Nonsynonymous | 7997 |
| DNAS_isl_43_MT818585 | 5826CtoT | T | I | Nonsynonymous | 3504 |
| DNAS_isl_43_MT818585 | 29625CtoT | S | F | Nonsynonymous | 7925 |
| DNAS_isl_43_MT818585 | 11462AtoG | M | V | Nonsynonymous | 7973 |
| DNAS_isl_43_MT818585 | 3037CtoT | F | F | Synonymous | 5368 |
| DNAS_isl_43_MT818585 | 18040GtoT | A | S | Nonsynonymous | 8008 |
| DNAS_isl_43_MT818585 | 25947GtoT | Q | H | Nonsynonymous | 8001 |
| DNAS_isl_43_MT818585 | 10658GtoA | V | I | Nonsynonymous | 7379 |
| DNAS_isl_43_MT818585 | 28881GtoA | R | K | Nonsynonymous | 3541 |
| DNAS_isl_43_MT818585 | 14408CtoT | P | L | Nonsynonymous | 3298 |
| DNAS_isl_43_MT818585 | 28883GtoC | G | R | Nonsynonymous | 3552 |
| DNAS_isl_43_MT818585 | 23403AtoG | D | G | Nonsynonymous | 7942 |
| DNAS_isl_43_MT818585 | 28882GtoA | R | K | Nonsynonymous | 3552 |
| DNAS_isl_43_MT818585 | 10369CtoT | R | R | Synonymous | 4399 |
| DNAS_isl_43_MT818585 | 241CtoT | R | C | 5'UTR SNP | 3265 |
| DNAS_isl_43_MT818585 | 23520CtoT | A | V | Nonsynonymous | 8002 |
| DNAS_isl_43_MT818585 | 1163AtoT | I | F | Nonsynonymous | 7060 |
| DNAS_isl_44_MT818581 | 1163AtoT | I | F | Nonsynonymous | 1915 |
| DNAS_isl_44_MT818581 | 241CtoT | R | C | 5'UTR SNP | 364 |
| DNAS_isl_44_MT818581 | 3037CtoT | F | F | Synonymous | 1998 |
| DNAS_isl_44_MT818581 | 28883GtoC | G | R | Nonsynonymous | 1537 |
| DNAS_isl_44_MT818581 | 23403AtoG | D | G | Nonsynonymous | 2809 |
| DNAS_isl_44_MT818581 | 3096CtoT | S | L | Nonsynonymous | 1085 |
| DNAS_isl_44_MT818581 | 28882GtoA | R | K | Nonsynonymous | 1536 |
| DNAS_isl_44_MT818581 | 28881GtoA | R | K | Nonsynonymous | 1527 |
| DNAS_isl_45_MT818589 | 4582CtoT | N | N | Synonymous | 7995 |
| DNAS_isl_45_MT818589 | 241CtoT | R | C | 5'UTR SNP | 3418 |
| DNAS_isl_45_MT818589 | 25904CtoT | S | L | Nonsynonymous | 5151 |
| DNAS_isl_45_MT818589 | 1163AtoT | I | F | Nonsynonymous | 7710 |
| DNAS_isl_45_MT818589 | 23403AtoG | D | G | Nonsynonymous | 7967 |
| DNAS_isl_45_MT818589 | 26211GtoT | V | V | Synonymous | 8005 |
| DNAS_isl_45_MT818589 | 3037CtoT | F | F | Synonymous | 4959 |
| DNAS_isl_45_MT818589 | 14408CtoT | P | L | Nonsynonymous | 1562 |
| DNAS_isl_45_MT818589 | 29440GtoT | Q | H | Nonsynonymous | 7967 |
| DNAS_isl_45_MT818589 | 21606GtoT | C | F | Nonsynonymous | 6388 |
| DNAS_isl_45_MT818589 | 20238GtoT | R | S | Nonsynonymous | 4189 |
| DNAS_isl_45_MT818589 | 28883GtoC | G | R | Nonsynonymous | 4183 |
| DNAS_isl_45_MT818589 | 28882GtoA | R | K | Nonsynonymous | 4182 |
| DNAS_isl_45_MT818589 | 5467CtoT | Y | Y | Synonymous | 6972 |
| DNAS_isl_45_MT818589 | 28881GtoA | R | K | Nonsynonymous | 4174 |
| DNAS_isl_46_MT818587 | 21606GtoT | C | F | Nonsynonymous | 4025 |
| DNAS_isl_46_MT818587 | 3037CtoT | F | F | Synonymous | 6129 |
| DNAS_isl_46_MT818587 | 23403AtoG | D | G | Nonsynonymous | 7912 |
| DNAS_isl_46_MT818587 | 4582CtoT | N | N | Synonymous | 3834 |
| DNAS_isl_46_MT818587 | 1163AtoT | I | F | Nonsynonymous | 7113 |
| DNAS_isl_46_MT818587 | 14408CtoT | P | L | Nonsynonymous | 1135 |
| DNAS_isl_46_MT818587 | 28883GtoC | G | R | Nonsynonymous | 4490 |
| DNAS_isl_46_MT818587 | 28882GtoA | R | K | Nonsynonymous | 4490 |
| DNAS_isl_46_MT818587 | 22023AtoG | E | G | Nonsynonymous | 3709 |
| DNAS_isl_46_MT818587 | 28881GtoA | R | K | Nonsynonymous | 4481 |
| DNAS_isl_46_MT818587 | 25904CtoT | S | L | Nonsynonymous | 2397 |
| DNAS_isl_46_MT818587 | 29440GtoT | Q | H | Nonsynonymous | 7935 |
| DNAS_isl_46_MT818587 | 26211GtoT | V | V | Synonymous | 8006 |
| DNAS_isl_46_MT818587 | 241CtoT | R | C | 5'UTR SNP | 3372 |
| DNAS_isl_46_MT818587 | 20238GtoT | R | S | Nonsynonymous | 2629 |
| DNAS_isl_47_MT818591 | 1163AtoT | I | F | Nonsynonymous | 5520 |
| DNAS_isl_47_MT818591 | 23403AtoG | D | G | Nonsynonymous | 7966 |
| DNAS_isl_47_MT818591 | 28882GtoA | R | K | Nonsynonymous | 5384 |
| DNAS_isl_47_MT818591 | 3037CtoT | F | F | Synonymous | 6665 |
| DNAS_isl_47_MT818591 | 28883GtoC | G | R | Nonsynonymous | 5384 |
| DNAS_isl_47_MT818591 | 20238GtoT | R | S | Nonsynonymous | 2700 |
| DNAS_isl_47_MT818591 | 28881GtoA | R | K | Nonsynonymous | 5377 |
| DNAS_isl_47_MT818591 | 22023AtoG | E | G | Nonsynonymous | 2966 |
| DNAS_isl_47_MT818591 | 20762CtoT | T | I | Nonsynonymous | 1266 |
| DNAS_isl_47_MT818591 | 25904CtoT | S | L | Nonsynonymous | 2839 |
| DNAS_isl_47_MT818591 | 26211GtoT | V | V | Synonymous | 7998 |
| DNAS_isl_47_MT818591 | 21606GtoT | C | F | Nonsynonymous | 2500 |
| DNAS_isl_47_MT818591 | 29440GtoT | Q | H | Nonsynonymous | 7960 |
| DNAS_isl_47_MT818591 | 14408CtoT | P | L | Nonsynonymous | 1096 |
| DNAS_isl_47_MT818591 | 241CtoT | R | C | 5'UTR SNP | 1929 |
| DNAS_isl_47_MT818591 | 4582CtoT | N | N | Synonymous | 3518 |
| DNAS_isl_48_MT818582 | 3037CtoT | F | F | Synonymous | 5370 |
| DNAS_isl_48_MT818582 | 14408CtoT | P | L | Nonsynonymous | 1576 |
| DNAS_isl_48_MT818582 | 1163AtoT | I | F | Nonsynonymous | 7625 |
| DNAS_isl_48_MT818582 | 5184CtoT | P | L | Nonsynonymous | 3950 |
| DNAS_isl_48_MT818582 | 29209AtoT | S | S | Synonymous | 3772 |
| DNAS_isl_48_MT818582 | 241CtoT | R | C | 5'UTR SNP | 2294 |
| DNAS_isl_48_MT818582 | 28881GtoA | R | K | Nonsynonymous | 4343 |
| DNAS_isl_48_MT818582 | 28882GtoA | R | K | Nonsynonymous | 4354 |
| DNAS_isl_48_MT818582 | 23403AtoG | D | G | Nonsynonymous | 7781 |
| DNAS_isl_48_MT818582 | 28883GtoC | G | R | Nonsynonymous | 4354 |
| DNAS_isl_49_MT818588 | 8146CtoT | D | D | Synonymous | 7083 |
| DNAS_isl_49_MT818588 | 28881GtoA | R | K | Nonsynonymous | 3574 |
| DNAS_isl_49_MT818588 | 3037CtoT | F | F | Synonymous | 4581 |
| DNAS_isl_49_MT818588 | 28882GtoA | R | K | Nonsynonymous | 3583 |
| DNAS_isl_49_MT818588 | 23403AtoG | D | G | Nonsynonymous | 7964 |
| DNAS_isl_49_MT818588 | 706CtoT | D | D | Synonymous | 1533 |
| DNAS_isl_49_MT818588 | 28883GtoC | G | R | Nonsynonymous | 3583 |
| DNAS_isl_49_MT818588 | 25599GtoT | W | C | Nonsynonymous | 3817 |
| DNAS_isl_49_MT818588 | 14408CtoT | P | L | Nonsynonymous | 4270 |
| DNAS_isl_49_MT818588 | 1163AtoT | I | F | Nonsynonymous | 6443 |
| DNAS_isl_49_MT818588 | 28342AtoT | S | S | Synonymous | 4936 |
| DNAS_isl_49_MT818588 | 17518CtoT | L | F | Nonsynonymous | 7993 |
| DNAS_isl_49_MT818588 | 241CtoT | R | C | 5'UTR SNP | 1412 |
| DNAS_isl_50_MT818580 | 23403AtoG | D | G | Nonsynonymous | 7374 |
| DNAS_isl_50_MT818580 | 14408CtoT | P | L | Nonsynonymous | 2011 |
| DNAS_isl_50_MT818580 | 1163AtoT | I | F | Nonsynonymous | 7802 |
| DNAS_isl_50_MT818580 | 241CtoT | R | C | 5'UTR SNP | 1911 |
| DNAS_isl_50_MT818580 | 3037CtoT | F | F | Synonymous | 5564 |
| DNAS_isl_50_MT818580 | 28882GtoA | R | K | Nonsynonymous | 3826 |
| DNAS_isl_50_MT818580 | 28883GtoC | G | R | Nonsynonymous | 3826 |
| DNAS_isl_50_MT818580 | 20274GtoT | M | I | Nonsynonymous | 3169 |
| DNAS_isl_50_MT818580 | 28881GtoA | R | K | Nonsynonymous | 3819 |
| DNAS_isl_51_MT818579 | 1163AtoT | I | F | Nonsynonymous | 7342 |
| DNAS_isl_51_MT818579 | 6807CtoT | T | I | Nonsynonymous | 7611 |
| DNAS_isl_51_MT818579 | 9451CtoT | Y | Y | Synonymous | 7978 |
| DNAS_isl_51_MT818579 | 19662CtoT | H | H | Synonymous | 6327 |
| DNAS_isl_51_MT818579 | 28883GtoC | G | R | Nonsynonymous | 3763 |
| DNAS_isl_51_MT818579 | 28881GtoA | R | K | Nonsynonymous | 3757 |
| DNAS_isl_51_MT818579 | 3037CtoT | F | F | Synonymous | 5665 |
| DNAS_isl_51_MT818579 | 28882GtoA | R | K | Nonsynonymous | 3762 |
| DNAS_isl_51_MT818579 | 24812GtoC | D | H | Nonsynonymous | 2467 |
| DNAS_isl_51_MT818579 | 12747CtoT | T | I | Nonsynonymous | 3709 |
| DNAS_isl_51_MT818579 | 26681CtoT | F | F | Synonymous | 7345 |
| DNAS_isl_51_MT818579 | 23403AtoG | D | G | Nonsynonymous | 7972 |
| DNAS_isl_51_MT818579 | 241CtoT | R | C | 5'UTR SNP | 2910 |
| DNAS_isl_51_MT818579 | 25460CtoT | A | V | Nonsynonymous | 7942 |
| DNAS_isl_51_MT818579 | 14408CtoT | P | L | Nonsynonymous | 3510 |
| DNAS_isl_51_MT818579 | 13308AtoG | K | R | Nonsynonymous | 7975 |
| DNAS_isl_52_MT800893 | 23593GtoT | Q | H | Nonsynonymous | 8003 |
| DNAS_isl_52_MT800893 | 23403AtoG | D | G | Nonsynonymous | 7948 |
| DNAS_isl_52_MT800893 | 241CtoT | R | C | 5'UTR SNP | 1534 |
| DNAS_isl_52_MT800893 | 2000AtoC | R | R | Synonymous | 7462 |
| DNAS_isl_52_MT800893 | 4138TtoC | V | V | Synonymous | 1008 |
| DNAS_isl_52_MT800893 | 1747GtoT | L | F | Nonsynonymous | 3437 |
| DNAS_isl_52_MT800893 | 11222GtoT | V | F | Nonsynonymous | 2706 |
| DNAS_isl_52_MT800893 | 3264CtoT | T | I | Nonsynonymous | 3266 |
| DNAS_isl_52_MT800893 | 14408CtoT | P | L | Nonsynonymous | 654 |
| DNAS_isl_52_MT800893 | 3037CtoT | F | F | Synonymous | 6322 |
| DNAS_isl_52_MT800893 | 28881GtoA | R | K | Nonsynonymous | 3990 |
| DNAS_isl_52_MT800893 | 28883GtoC | G | R | Nonsynonymous | 3995 |
| DNAS_isl_52_MT800893 | 28882GtoA | R | K | Nonsynonymous | 3995 |
| DNAS_isl_52_MT800893 | 1163AtoT | I | F | Nonsynonymous | 3385 |
| DNAS_isl_53_MT800884 | 1163AtoT | I | F | Nonsynonymous | 4654 |
| DNAS_isl_53_MT800884 | 241CtoT | R | C | 5'UTR SNP | 1857 |
| DNAS_isl_53_MT800884 | 14408CtoT | P | L | Nonsynonymous | 672 |
| DNAS_isl_53_MT800884 | 3037CtoT | F | F | Synonymous | 5551 |
| DNAS_isl_53_MT800884 | 23403AtoG | D | G | Nonsynonymous | 7656 |
| DNAS_isl_53_MT800884 | 28882GtoA | R | K | Nonsynonymous | 4848 |
| DNAS_isl_53_MT800884 | 28883GtoC | G | R | Nonsynonymous | 4848 |
| DNAS_isl_53_MT800884 | 28881GtoA | R | K | Nonsynonymous | 4843 |
| DNAS_isl_54_MT800876 | 28079GtoT | V | V | Synonymous | 7950 |
| DNAS_isl_54_MT800876 | 3037CtoT | F | F | Synonymous | 7728 |
| DNAS_isl_54_MT800876 | 28883GtoC | G | R | Nonsynonymous | 2827 |
| DNAS_isl_54_MT800876 | 1163AtoT | I | F | Nonsynonymous | 7588 |
| DNAS_isl_54_MT800876 | 28882GtoA | R | K | Nonsynonymous | 2827 |
| DNAS_isl_54_MT800876 | 23403AtoG | D | G | Nonsynonymous | 7940 |
| DNAS_isl_54_MT800876 | 28881GtoA | R | K | Nonsynonymous | 2822 |
| DNAS_isl_54_MT800876 | 241CtoT | R | C | 5'UTR SNP | 2915 |
| DNAS_isl_54_MT800876 | 3053GtoT | D | Y | Nonsynonymous | 5539 |
| DNAS_isl_54_MT800876 | 17592GtoT | V | V | Synonymous | 5556 |
| DNAS_isl_54_MT800876 | 14408CtoT | P | L | Nonsynonymous | 2933 |
| DNAS_isl_55_MT800881 | 28882GtoA | R | K | Nonsynonymous | 4616 |
| DNAS_isl_55_MT800881 | 14408CtoT | P | L | Nonsynonymous | 2406 |
| DNAS_isl_55_MT800881 | 28881GtoA | R | K | Nonsynonymous | 4607 |
| DNAS_isl_55_MT800881 | 241CtoT | R | C | 5'UTR SNP | 2579 |
| DNAS_isl_55_MT800881 | 1163AtoT | I | F | Nonsynonymous | 7577 |
| DNAS_isl_55_MT800881 | 7177AtoG | L | L | Synonymous | 2783 |
| DNAS_isl_55_MT800881 | 434GtoA | E | K | Nonsynonymous | 7595 |
| DNAS_isl_55_MT800881 | 3037CtoT | F | F | Synonymous | 5965 |
| DNAS_isl_55_MT800881 | 23403AtoG | D | G | Nonsynonymous | 7961 |
| DNAS_isl_55_MT800881 | 23604CtoG | P | R | Nonsynonymous | 7364 |
| DNAS_isl_55_MT800881 | 28883GtoC | G | R | Nonsynonymous | 4616 |
| DNAS_isl_56_MT800882 | 1163AtoT | I | F | Nonsynonymous | 3867 |
| DNAS_isl_56_MT800882 | 23604CtoG | P | R | Nonsynonymous | 6548 |
| DNAS_isl_56_MT800882 | 7177AtoG | L | L | Synonymous | 1162 |
| DNAS_isl_56_MT800882 | 28883GtoC | G | R | Nonsynonymous | 4037 |
| DNAS_isl_56_MT800882 | 3037CtoT | F | F | Synonymous | 4575 |
| DNAS_isl_56_MT800882 | 241CtoT | R | C | 5'UTR SNP | 1961 |
| DNAS_isl_56_MT800882 | 14408CtoT | P | L | Nonsynonymous | 1212 |
| DNAS_isl_56_MT800882 | 28881GtoA | R | K | Nonsynonymous | 4027 |
| DNAS_isl_56_MT800882 | 28882GtoA | R | K | Nonsynonymous | 4037 |
| DNAS_isl_56_MT800882 | 434GtoA | E | K | Nonsynonymous | 4770 |
| DNAS_isl_56_MT800882 | 23403AtoG | D | G | Nonsynonymous | 7361 |
| DNAS_isl_57_MT800888 | 3037CtoT | F | F | Synonymous | 5734 |
| DNAS_isl_57_MT800888 | 23403AtoG | D | G | Nonsynonymous | 7977 |
| DNAS_isl_57_MT800888 | 28881GtoA | R | K | Nonsynonymous | 4104 |
| DNAS_isl_57_MT800888 | 7177AtoG | L | L | Synonymous | 3824 |
| DNAS_isl_57_MT800888 | 241CtoT | R | C | 5'UTR SNP | 2936 |
| DNAS_isl_57_MT800888 | 14408CtoT | P | L | Nonsynonymous | 3004 |
| DNAS_isl_57_MT800888 | 28882GtoA | R | K | Nonsynonymous | 4111 |
| DNAS_isl_57_MT800888 | 23604CtoG | P | R | Nonsynonymous | 7994 |
| DNAS_isl_57_MT800888 | 28883GtoC | G | R | Nonsynonymous | 4112 |
| DNAS_isl_57_MT800888 | 2731GtoA | K | K | Synonymous | 7940 |
| DNAS_isl_57_MT800888 | 434GtoA | E | K | Nonsynonymous | 7628 |
| DNAS_isl_57_MT800888 | 1163AtoT | I | F | Nonsynonymous | 5915 |
| DNAS_isl_58_MT800886 | 28904GtoT | A | S | Nonsynonymous | 7991 |
| DNAS_isl_58_MT800886 | 241CtoT | R | C | 5'UTR SNP | 1906 |
| DNAS_isl_58_MT800886 | 25997TtoC | V | A | Nonsynonymous | 3541 |
| DNAS_isl_58_MT800886 | 28883GtoC | G | R | Nonsynonymous | 4684 |
| DNAS_isl_58_MT800886 | 22017GtoT | W | L | Nonsynonymous | 2150 |
| DNAS_isl_58_MT800886 | 10377CtoT | P | L | Nonsynonymous | 5428 |
| DNAS_isl_58_MT800886 | 1163AtoT | I | F | Nonsynonymous | 6946 |
| DNAS_isl_58_MT800886 | 28881GtoA | R | K | Nonsynonymous | 4671 |
| DNAS_isl_58_MT800886 | 28882GtoA | R | K | Nonsynonymous | 4684 |
| DNAS_isl_58_MT800886 | 23403AtoG | D | G | Nonsynonymous | 6239 |
| DNAS_isl_58_MT800886 | 3037CtoT | F | F | Synonymous | 5224 |
| DNAS_isl_58_MT800886 | 26885CtoT | N | N | Synonymous | 4243 |
| DNAS_isl_58_MT800886 | 14408CtoT | P | L | Nonsynonymous | 1498 |
| DNAS_isl_58_MT800886 | 21989GtoT | V | F | Nonsynonymous | 1967 |
| DNAS_isl_59_MT800889 | 29118CtoT | T | I | Nonsynonymous | 4933 |
| DNAS_isl_59_MT800889 | 28178GtoT | L | F | Nonsynonymous | 7951 |
| DNAS_isl_59_MT800889 | 241CtoT | R | C | 5'UTR SNP | 1196 |
| DNAS_isl_59_MT800889 | 14408CtoT | P | L | Nonsynonymous | 3718 |
| DNAS_isl_59_MT800889 | 29262GtoT | W | L | Nonsynonymous | 7925 |
| DNAS_isl_59_MT800889 | 28883GtoC | G | R | Nonsynonymous | 3587 |
| DNAS_isl_59_MT800889 | 28882GtoA | R | K | Nonsynonymous | 3586 |
| DNAS_isl_59_MT800889 | 23403AtoG | D | G | Nonsynonymous | 7952 |
| DNAS_isl_59_MT800889 | 28881GtoA | R | K | Nonsynonymous | 3575 |
| DNAS_isl_59_MT800889 | 3037CtoT | F | F | Synonymous | 5185 |
| DNAS_isl_60_MT800887 | 241CtoT | R | C | 5'UTR SNP | 1101 |
| DNAS_isl_60_MT800887 | 28881GtoA | R | K | Nonsynonymous | 3714 |
| DNAS_isl_60_MT800887 | 28883GtoC | G | R | Nonsynonymous | 3721 |
| DNAS_isl_60_MT800887 | 23403AtoG | D | G | Nonsynonymous | 4587 |
| DNAS_isl_60_MT800887 | 28882GtoA | R | K | Nonsynonymous | 3721 |
| DNAS_isl_60_MT800887 | 12076CtoT | N | N | Synonymous | 2609 |
| DNAS_isl_60_MT800887 | 2040CtoT | T | I | Nonsynonymous | 1984 |
| DNAS_isl_60_MT800887 | 25504CtoG | Q | E | Nonsynonymous | 1669 |
| DNAS_isl_60_MT800887 | 14408CtoT | P | L | Nonsynonymous | 464 |
| DNAS_isl_60_MT800887 | 3037CtoT | F | F | Synonymous | 1806 |
| DNAS_isl_60_MT800887 | 25855GtoT | D | Y | Nonsynonymous | 1442 |
| DNAS_isl_60_MT800887 | 1163AtoT | I | F | Nonsynonymous | 2691 |
| DNAS_isl_60_MT800887 | 22648TtoA | V | V | Synonymous | 1168 |
| DNAS_isl_61_MT800875 | 14408CtoT | P | L | Nonsynonymous | 3294 |
| DNAS_isl_61_MT800875 | 6402CtoT | P | L | Nonsynonymous | 6345 |
| DNAS_isl_61_MT800875 | 1163AtoT | I | F | Nonsynonymous | 7596 |
| DNAS_isl_61_MT800875 | 23403AtoG | D | G | Nonsynonymous | 7986 |
| DNAS_isl_61_MT800875 | 25540GtoA | V | I | Nonsynonymous | 7986 |
| DNAS_isl_61_MT800875 | 3037CtoT | F | F | Synonymous | 5954 |
| DNAS_isl_61_MT800875 | 25904CtoT | S | L | Nonsynonymous | 3752 |
| DNAS_isl_61_MT800875 | 28082TtoC | D | D | Synonymous | 7979 |
| DNAS_isl_61_MT800875 | 6807CtoT | T | I | Nonsynonymous | 6077 |
| DNAS_isl_61_MT800875 | 28883GtoC | G | R | Nonsynonymous | 4016 |
| DNAS_isl_61_MT800875 | 241CtoT | R | C | 5'UTR SNP | 3045 |
| DNAS_isl_61_MT800875 | 28881GtoA | R | K | Nonsynonymous | 4006 |
| DNAS_isl_61_MT800875 | 28882GtoA | R | K | Nonsynonymous | 4016 |
| DNAS_isl_62_MT800894 | 26022CtoT | D | D | Synonymous | 5121 |
| DNAS_isl_62_MT800894 | 241CtoT | R | C | 5'UTR SNP | 4417 |
| DNAS_isl_62_MT800894 | 14408CtoT | P | L | Nonsynonymous | 412 |
| DNAS_isl_62_MT800894 | 3037CtoT | F | F | Synonymous | 7022 |
| DNAS_isl_62_MT800894 | 10323AtoG | K | R | Nonsynonymous | 8002 |
| DNAS_isl_62_MT800894 | 1163AtoT | I | F | Nonsynonymous | 4907 |
| DNAS_isl_62_MT800894 | 8692CtoT | Y | Y | Synonymous | 1796 |
| DNAS_isl_62_MT800894 | 29171CtoT | H | Y | Nonsynonymous | 5725 |
| DNAS_isl_62_MT800894 | 11083GtoT | L | F | Nonsynonymous | 817 |
| DNAS_isl_62_MT800894 | 23403AtoG | D | G | Nonsynonymous | 7953 |
| DNAS_isl_62_MT800894 | 6701CtoT | L | F | Nonsynonymous | 6450 |
| DNAS_isl_62_MT800894 | 28881GtoA | R | K | Nonsynonymous | 5501 |
| DNAS_isl_62_MT800894 | 28882GtoA | R | K | Nonsynonymous | 5509 |
| DNAS_isl_62_MT800894 | 28883GtoC | G | R | Nonsynonymous | 5510 |
| DNAS_isl_62_MT800894 | 13686TtoC | H | H | Synonymous | 3090 |
| DNAS_isl_63_MT800892 | 1457CtoT | R | C | Nonsynonymous | 3322 |
| DNAS_isl_63_MT800892 | 3037CtoT | F | F | Synonymous | 6861 |
| DNAS_isl_63_MT800892 | 23403AtoG | D | G | Nonsynonymous | 7948 |
| DNAS_isl_63_MT800892 | 2836CtoT | C | C | Synonymous | 7936 |
| DNAS_isl_63_MT800892 | 241CtoT | R | C | 5'UTR SNP | 3135 |
| DNAS_isl_63_MT800892 | 14408CtoT | P | L | Nonsynonymous | 324 |
| DNAS_isl_63_MT800892 | 28854CtoT | S | L | Nonsynonymous | 7963 |
| DNAS_isl_63_MT800892 | 25563GtoT | Q | H | Nonsynonymous | 7949 |
| DNAS_isl_63_MT800892 | 6367GtoT | Q | H | Nonsynonymous | 849 |
| DNAS_isl_63_MT800892 | 22444CtoT | D | D | Synonymous | 5177 |
| DNAS_isl_63_MT800892 | 26735CtoT | Y | Y | Synonymous | 1578 |
| DNAS_isl_63_MT800892 | 27703GtoT | V | F | Nonsynonymous | 138 |
| DNAS_isl_63_MT800892 | 18877CtoT | L | L | Synonymous | 7947 |
| DNAS_isl_64_MT800891 | 28854CtoT | S | L | Nonsynonymous | 7951 |
| DNAS_isl_64_MT800891 | 26735CtoT | Y | Y | Synonymous | 2137 |
| DNAS_isl_64_MT800891 | 27703GtoT | V | F | Nonsynonymous | 1302 |
| DNAS_isl_64_MT800891 | 22444CtoT | D | D | Synonymous | 2908 |
| DNAS_isl_64_MT800891 | 241CtoT | R | C | 5'UTR SNP | 1144 |
| DNAS_isl_64_MT800891 | 18877CtoT | L | L | Synonymous | 3013 |
| DNAS_isl_64_MT800891 | 3037CtoT | F | F | Synonymous | 2443 |
| DNAS_isl_64_MT800891 | 6367GtoT | Q | H | Nonsynonymous | 674 |
| DNAS_isl_64_MT800891 | 2836CtoT | C | C | Synonymous | 2526 |
| DNAS_isl_64_MT800891 | 23403AtoG | D | G | Nonsynonymous | 5139 |
| DNAS_isl_64_MT800891 | 25563GtoT | Q | H | Nonsynonymous | 4576 |
| DNAS_isl_64_MT800891 | 14408CtoT | P | L | Nonsynonymous | 659 |
| DNAS_isl_65_MT800877 | 3037CtoT | F | F | Synonymous | 5620 |
| DNAS_isl_65_MT800877 | 23403AtoG | D | G | Nonsynonymous | 7957 |
| DNAS_isl_65_MT800877 | 28882GtoA | R | K | Nonsynonymous | 3659 |
| DNAS_isl_65_MT800877 | 28881GtoA | R | K | Nonsynonymous | 3650 |
| DNAS_isl_65_MT800877 | 26211GtoT | V | V | Synonymous | 8018 |
| DNAS_isl_65_MT800877 | 14408CtoT | P | L | Nonsynonymous | 3445 |
| DNAS_isl_65_MT800877 | 25904CtoT | S | L | Nonsynonymous | 3770 |
| DNAS_isl_65_MT800877 | 21575CtoT | L | F | Nonsynonymous | 1136 |
| DNAS_isl_65_MT800877 | 1163AtoT | I | F | Nonsynonymous | 7461 |
| DNAS_isl_65_MT800877 | 20238GtoT | R | S | Nonsynonymous | 5085 |
| DNAS_isl_65_MT800877 | 28883GtoC | G | R | Nonsynonymous | 3660 |
| DNAS_isl_65_MT800877 | 241CtoT | R | C | 5'UTR SNP | 2571 |
| DNAS_isl_66_MT800879 | 17518CtoT | L | F | Nonsynonymous | 5099 |
| DNAS_isl_66_MT800879 | 241CtoT | R | C | 5'UTR SNP | 2283 |
| DNAS_isl_66_MT800879 | 1163AtoT | I | F | Nonsynonymous | 5340 |
| DNAS_isl_66_MT800879 | 3037CtoT | F | F | Synonymous | 4003 |
| DNAS_isl_66_MT800879 | 8146CtoT | D | D | Synonymous | 3086 |
| DNAS_isl_66_MT800879 | 19344TtoC | A | A | Synonymous | 2787 |
| DNAS_isl_66_MT800879 | 28882GtoA | R | K | Nonsynonymous | 4315 |
| DNAS_isl_66_MT800879 | 28881GtoA | R | K | Nonsynonymous | 4306 |
| DNAS_isl_66_MT800879 | 28883GtoC | G | R | Nonsynonymous | 4315 |
| DNAS_isl_66_MT800879 | 14408CtoT | P | L | Nonsynonymous | 828 |
| DNAS_isl_66_MT800879 | 23403AtoG | D | G | Nonsynonymous | 7569 |
| DNAS_isl_67_MT800880 | 3037CtoT | F | F | Synonymous | 5801 |
| DNAS_isl_67_MT800880 | 28881GtoA | R | K | Nonsynonymous | 4149 |
| DNAS_isl_67_MT800880 | 28882GtoA | R | K | Nonsynonymous | 4159 |
| DNAS_isl_67_MT800880 | 28883GtoC | G | R | Nonsynonymous | 4159 |
| DNAS_isl_67_MT800880 | 1163AtoT | I | F | Nonsynonymous | 7418 |
| DNAS_isl_67_MT800880 | 12488CtoT | P | S | Nonsynonymous | 5275 |
| DNAS_isl_67_MT800880 | 241CtoT | R | C | 5'UTR SNP | 2847 |
| DNAS_isl_67_MT800880 | 14292CtoT | D | D | Synonymous | 6829 |
| DNAS_isl_67_MT800880 | 23403AtoG | D | G | Nonsynonymous | 7951 |
| DNAS_isl_67_MT800880 | 14408CtoT | P | L | Nonsynonymous | 3360 |
| DNAS_isl_68_MT800878 | 23403AtoG | D | G | Nonsynonymous | 2388 |
| DNAS_isl_68_MT800878 | 1163AtoT | I | F | Nonsynonymous | 1562 |
| DNAS_isl_68_MT800878 | 27476CtoT | T | I | Nonsynonymous | 2666 |
| DNAS_isl_68_MT800878 | 241CtoT | R | C | 5'UTR SNP | 598 |
| DNAS_isl_68_MT800878 | 1191CtoT | P | L | Nonsynonymous | 1568 |
| DNAS_isl_68_MT800878 | 3037CtoT | F | F | Synonymous | 3203 |
| DNAS_isl_68_MT800878 | 14408CtoT | P | L | Nonsynonymous | 155 |
| DNAS_isl_68_MT800878 | 28882GtoA | R | K | Nonsynonymous | 3404 |
| DNAS_isl_68_MT800878 | 2144GtoT | V | F | Nonsynonymous | 1657 |
| DNAS_isl_68_MT800878 | 28883GtoC | G | R | Nonsynonymous | 3404 |
| DNAS_isl_68_MT800878 | 17944GtoT | V | L | Nonsynonymous | 1563 |
| DNAS_isl_68_MT800878 | 28881GtoA | R | K | Nonsynonymous | 3396 |
| DNAS_isl_69_MT800885 | 28881GtoA | R | K | Nonsynonymous | 5224 |
| DNAS_isl_69_MT800885 | 28882GtoA | R | K | Nonsynonymous | 5234 |
| DNAS_isl_69_MT800885 | 28883GtoC | G | R | Nonsynonymous | 5235 |
| DNAS_isl_69_MT800885 | 3037CtoT | F | F | Synonymous | 6066 |
| DNAS_isl_69_MT800885 | 23403AtoG | D | G | Nonsynonymous | 7940 |
| DNAS_isl_69_MT800885 | 18672TtoC | D | D | Synonymous | 1058 |
| DNAS_isl_69_MT800885 | 28888TtoC | T | T | Synonymous | 5437 |
| DNAS_isl_69_MT800885 | 241CtoT | R | C | 5'UTR SNP | 1967 |
| DNAS_isl_69_MT800885 | 4754CtoT | P | S | Nonsynonymous | 4594 |
| DNAS_isl_69_MT800885 | 14408CtoT | P | L | Nonsynonymous | 599 |
| DNAS_isl_69_MT800885 | 1163AtoT | I | F | Nonsynonymous | 7894 |
| DNAS_isl_69_MT800885 | 21707CtoT | H | Y | Nonsynonymous | 1689 |
| DNAS_isl_70_MT800890 | 6449CtoT | L | F | Nonsynonymous | 664 |
| DNAS_isl_70_MT800890 | 683CtoT | L | L | Synonymous | 2504 |
| DNAS_isl_70_MT800890 | 23403AtoG | D | G | Nonsynonymous | 2227 |
| DNAS_isl_70_MT800890 | 3037CtoT | F | F | Synonymous | 2821 |
| DNAS_isl_70_MT800890 | 28882GtoA | R | K | Nonsynonymous | 2104 |
| DNAS_isl_70_MT800890 | 28883GtoC | G | R | Nonsynonymous | 2104 |
| DNAS_isl_70_MT800890 | 241CtoT | R | C | 5'UTR SNP | 736 |
| DNAS_isl_70_MT800890 | 2939CtoT | P | S | Nonsynonymous | 2887 |
| DNAS_isl_70_MT800890 | 28881GtoA | R | K | Nonsynonymous | 2098 |
| DNAS_isl_71_MT800883 | 19032CtoT | D | D | Synonymous | 1783 |
| DNAS_isl_71_MT800883 | 5392CtoT | N | N | Synonymous | 5233 |
| DNAS_isl_71_MT800883 | 28881GtoA | R | K | Nonsynonymous | 3016 |
| DNAS_isl_71_MT800883 | 1163AtoT | I | F | Nonsynonymous | 6758 |
| DNAS_isl_71_MT800883 | 241CtoT | R | C | 5'UTR SNP | 2914 |
| DNAS_isl_71_MT800883 | 28883GtoC | G | R | Nonsynonymous | 3029 |
| DNAS_isl_71_MT800883 | 28882GtoA | R | K | Nonsynonymous | 3029 |
| DNAS_isl_71_MT800883 | 23403AtoG | D | G | Nonsynonymous | 7982 |
| DNAS_isl_71_MT800883 | 14408CtoT | P | L | Nonsynonymous | 1933 |
| DNAS_isl_71_MT800883 | 712TtoA | L | L | Synonymous | 3715 |
| DNAS_isl_71_MT800883 | 3037CtoT | F | F | Synonymous | 5348 |
| DNAS_isl_71_MT800883 | 12086TtoC | L | L | Synonymous | 7894 |
| DNAS_isl_72_MT775562 | 28881GtoA | R | K | Nonsynonymous | 3070 |
| DNAS_isl_72_MT775562 | 1163AtoT | I | F | Nonsynonymous | 7376 |
| DNAS_isl_72_MT775562 | 28882GtoA | R | K | Nonsynonymous | 3078 |
| DNAS_isl_72_MT775562 | 3053GtoT | D | Y | Nonsynonymous | 3462 |
| DNAS_isl_72_MT775562 | 28883GtoC | G | R | Nonsynonymous | 3078 |
| DNAS_isl_72_MT775562 | 23403AtoG | D | G | Nonsynonymous | 7951 |
| DNAS_isl_72_MT775562 | 28079GtoT | V | V | Synonymous | 7956 |
| DNAS_isl_72_MT775562 | 3037CtoT | F | F | Synonymous | 7671 |
| DNAS_isl_72_MT775562 | 241CtoT | R | C | 5'UTR SNP | 2434 |
| DNAS_isl_72_MT775562 | 14408CtoT | P | L | Nonsynonymous | 3923 |
| DNAS_isl_73_MT775565 | 10074CtoT | A | V | Nonsynonymous | 4296 |
| DNAS_isl_73_MT775565 | 3037CtoT | F | F | Synonymous | 6039 |
| DNAS_isl_73_MT775565 | 5497CtoT | C | C | Synonymous | 1725 |
| DNAS_isl_73_MT775565 | 22113GtoA | G | D | Nonsynonymous | 2733 |
| DNAS_isl_73_MT775565 | 23403AtoG | D | G | Nonsynonymous | 7767 |
| DNAS_isl_73_MT775565 | 14408CtoT | P | L | Nonsynonymous | 1964 |
| DNAS_isl_73_MT775565 | 21575CtoT | L | F | Nonsynonymous | 100 |
| DNAS_isl_73_MT775565 | 1163AtoT | I | F | Nonsynonymous | 4231 |
| DNAS_isl_73_MT775565 | 28883GtoC | G | R | Nonsynonymous | 4306 |
| DNAS_isl_73_MT775565 | 28882GtoA | R | K | Nonsynonymous | 4306 |
| DNAS_isl_73_MT775565 | 241CtoT | R | C | 5'UTR SNP | 951 |
| DNAS_isl_73_MT775565 | 28881GtoA | R | K | Nonsynonymous | 4296 |
| DNAS_isl_74_MT775564 | 3037CtoT | F | F | Synonymous | 6171 |
| DNAS_isl_74_MT775564 | 21575CtoT | L | F | Nonsynonymous | 1887 |
| DNAS_isl_74_MT775564 | 241CtoT | R | C | 5'UTR SNP | 5491 |
| DNAS_isl_74_MT775564 | 5497CtoT | C | C | Synonymous | 7418 |
| DNAS_isl_74_MT775564 | 28883GtoC | G | R | Nonsynonymous | 6152 |
| DNAS_isl_74_MT775564 | 28882GtoA | R | K | Nonsynonymous | 6152 |
| DNAS_isl_74_MT775564 | 23403AtoG | D | G | Nonsynonymous | 7950 |
| DNAS_isl_74_MT775564 | 1163AtoT | I | F | Nonsynonymous | 6783 |
| DNAS_isl_74_MT775564 | 28881GtoA | R | K | Nonsynonymous | 6147 |
| DNAS_isl_75_MT775558 | 14408CtoT | P | L | Nonsynonymous | 2650 |
| DNAS_isl_75_MT775558 | 241CtoT | R | C | 5'UTR SNP | 3852 |
| DNAS_isl_75_MT775558 | 3037CtoT | F | F | Synonymous | 5578 |
| DNAS_isl_75_MT775558 | 1163AtoT | I | F | Nonsynonymous | 7190 |
| DNAS_isl_75_MT775558 | 28882GtoA | R | K | Nonsynonymous | 5560 |
| DNAS_isl_75_MT775558 | 28881GtoA | R | K | Nonsynonymous | 5554 |
| DNAS_isl_75_MT775558 | 28883GtoC | G | R | Nonsynonymous | 5560 |
| DNAS_isl_75_MT775558 | 6445CtoT | D | D | Synonymous | 8004 |
| DNAS_isl_75_MT775558 | 28122GtoT | G | C | Nonsynonymous | 4211 |
| DNAS_isl_75_MT775558 | 23403AtoG | D | G | Nonsynonymous | 7975 |
| DNAS_isl_75_MT775558 | 21575CtoT | L | F | Nonsynonymous | 1411 |
| DNAS_isl_75_MT775558 | 22801GtoT | G | G | Synonymous | 8001 |
| DNAS_isl_76_MT775563 | 3961CtoT | I | I | Synonymous | 3172 |
| DNAS_isl_76_MT775563 | 3037CtoT | F | F | Synonymous | 5025 |
| DNAS_isl_76_MT775563 | 241CtoT | R | C | 5'UTR SNP | 3330 |
| DNAS_isl_76_MT775563 | 14408CtoT | P | L | Nonsynonymous | 2749 |
| DNAS_isl_76_MT775563 | 23403AtoG | D | G | Nonsynonymous | 7969 |
| DNAS_isl_76_MT775563 | 28883GtoC | G | R | Nonsynonymous | 2258 |
| DNAS_isl_76_MT775563 | 1163AtoT | I | F | Nonsynonymous | 7019 |
| DNAS_isl_76_MT775563 | 28882GtoA | R | K | Nonsynonymous | 2258 |
| DNAS_isl_76_MT775563 | 28881GtoA | R | K | Nonsynonymous | 2251 |
| DNAS_isl_76_MT775563 | 19600AtoC | N | H | Nonsynonymous | 5863 |
| DNAS_isl_77_MT775571 | 28881GtoA | R | K | Nonsynonymous | 3588 |
| DNAS_isl_77_MT775571 | 1163AtoT | I | F | Nonsynonymous | 7087 |
| DNAS_isl_77_MT775571 | 3037CtoT | F | F | Synonymous | 4663 |
| DNAS_isl_77_MT775571 | 23403AtoG | D | G | Nonsynonymous | 7949 |
| DNAS_isl_77_MT775571 | 28200TtoC | S | P | Nonsynonymous | 4940 |
| DNAS_isl_77_MT775571 | 12525CtoT | T | I | Nonsynonymous | 4656 |
| DNAS_isl_77_MT775571 | 19648GtoT | V | L | Nonsynonymous | 6149 |
| DNAS_isl_77_MT775571 | 241CtoT | R | C | 5'UTR SNP | 2574 |
| DNAS_isl_77_MT775571 | 28882GtoA | R | K | Nonsynonymous | 3596 |
| DNAS_isl_77_MT775571 | 14408CtoT | P | L | Nonsynonymous | 3138 |
| DNAS_isl_77_MT775571 | 28883GtoC | G | R | Nonsynonymous | 3598 |
| DNAS_isl_78_MT775570 | 241CtoT | R | C | 5'UTR SNP | 3252 |
| DNAS_isl_78_MT775570 | 3037CtoT | F | F | Synonymous | 5464 |
| DNAS_isl_78_MT775570 | 23403AtoG | D | G | Nonsynonymous | 7979 |
| DNAS_isl_78_MT775570 | 28200TtoC | S | P | Nonsynonymous | 6755 |
| DNAS_isl_78_MT775570 | 12525CtoT | T | I | Nonsynonymous | 2971 |
| DNAS_isl_78_MT775570 | 1163AtoT | I | F | Nonsynonymous | 6373 |
| DNAS_isl_78_MT775570 | 19648GtoT | V | L | Nonsynonymous | 4323 |
| DNAS_isl_78_MT775570 | 14408CtoT | P | L | Nonsynonymous | 1802 |
| DNAS_isl_78_MT775570 | 28881GtoA | R | K | Nonsynonymous | 4840 |
| DNAS_isl_78_MT775570 | 28883GtoC | G | R | Nonsynonymous | 4859 |
| DNAS_isl_78_MT775570 | 28882GtoA | R | K | Nonsynonymous | 4853 |
| DNAS_isl_79_MT775568 | 4802GtoA | D | N | Nonsynonymous | 5002 |
| DNAS_isl_79_MT775568 | 482CtoT | R | C | Nonsynonymous | 7964 |
| DNAS_isl_79_MT775568 | 3554CtoA | L | I | Nonsynonymous | 5921 |
| DNAS_isl_79_MT775568 | 14408CtoT | P | L | Nonsynonymous | 2347 |
| DNAS_isl_79_MT775568 | 28881GtoA | R | K | Nonsynonymous | 5015 |
| DNAS_isl_79_MT775568 | 241CtoT | R | C | 5'UTR SNP | 3763 |
| DNAS_isl_79_MT775568 | 26774GtoT | M | I | Nonsynonymous | 6824 |
| DNAS_isl_79_MT775568 | 28883GtoC | G | R | Nonsynonymous | 5027 |
| DNAS_isl_79_MT775568 | 23403AtoG | D | G | Nonsynonymous | 7976 |
| DNAS_isl_79_MT775568 | 1163AtoT | I | F | Nonsynonymous | 7270 |
| DNAS_isl_79_MT775568 | 28882GtoA | R | K | Nonsynonymous | 5027 |
| DNAS_isl_79_MT775568 | 26211GtoT | V | V | Synonymous | 7984 |
| DNAS_isl_79_MT775568 | 3037CtoT | F | F | Synonymous | 5745 |
| DNAS_isl_80_MT775560 | 28882GtoA | R | K | Nonsynonymous | 3605 |
| DNAS_isl_80_MT775560 | 28881GtoA | R | K | Nonsynonymous | 3598 |
| DNAS_isl_80_MT775560 | 4579TtoA | L | L | Synonymous | 7987 |
| DNAS_isl_80_MT775560 | 28883GtoC | G | R | Nonsynonymous | 3605 |
| DNAS_isl_80_MT775560 | 21575CtoT | L | F | Nonsynonymous | 439 |
| DNAS_isl_80_MT775560 | 23403AtoG | D | G | Nonsynonymous | 7990 |
| DNAS_isl_80_MT775560 | 23635CtoT | S | S | Synonymous | 8017 |
| DNAS_isl_80_MT775560 | 3037CtoT | F | F | Synonymous | 3605 |
| DNAS_isl_80_MT775560 | 14408CtoT | P | L | Nonsynonymous | 2303 |
| DNAS_isl_80_MT775560 | 241CtoT | R | C | 5'UTR SNP | 745 |
| DNAS_isl_80_MT775560 | 1163AtoT | I | F | Nonsynonymous | 7345 |
| DNAS_isl_81_MT775566 | 1163AtoT | I | F | Nonsynonymous | 7250 |
| DNAS_isl_81_MT775566 | 28882GtoA | R | K | Nonsynonymous | 4361 |
| DNAS_isl_81_MT775566 | 28881GtoA | R | K | Nonsynonymous | 4354 |
| DNAS_isl_81_MT775566 | 28883GtoC | G | R | Nonsynonymous | 4361 |
| DNAS_isl_81_MT775566 | 14408CtoT | P | L | Nonsynonymous | 2650 |
| DNAS_isl_81_MT775566 | 23403AtoG | D | G | Nonsynonymous | 7928 |
| DNAS_isl_81_MT775566 | 241CtoT | R | C | 5'UTR SNP | 2665 |
| DNAS_isl_81_MT775566 | 3037CtoT | F | F | Synonymous | 5746 |
| DNAS_isl_82_MT775567 | 26259TtoG | V | V | Synonymous | 8007 |
| DNAS_isl_82_MT775567 | 21123GtoT | G | G | Synonymous | 7905 |
| DNAS_isl_82_MT775567 | 23403AtoG | D | G | Nonsynonymous | 7950 |
| DNAS_isl_82_MT775567 | 14408CtoT | P | L | Nonsynonymous | 3050 |
| DNAS_isl_82_MT775567 | 20134GtoT | V | L | Nonsynonymous | 5168 |
| DNAS_isl_82_MT775567 | 28883GtoC | G | R | Nonsynonymous | 3164 |
| DNAS_isl_82_MT775567 | 3037CtoT | F | F | Synonymous | 6179 |
| DNAS_isl_82_MT775567 | 28882GtoA | R | K | Nonsynonymous | 3163 |
| DNAS_isl_82_MT775567 | 28881GtoA | R | K | Nonsynonymous | 3143 |
| DNAS_isl_82_MT775567 | 19858GtoT | A | S | Nonsynonymous | 6325 |
| DNAS_isl_82_MT775567 | 1163AtoT | I | F | Nonsynonymous | 7011 |
| DNAS_isl_82_MT775567 | 8650CtoT | V | V | Synonymous | 4759 |
| DNAS_isl_82_MT775567 | 241CtoT | R | C | 5'UTR SNP | 2426 |
| DNAS_isl_82_MT775567 | 25690GtoT | G | C | Nonsynonymous | 7777 |
| DNAS_isl_82_MT775567 | 22925TtoC | L | L | Synonymous | 7972 |
| DNAS_isl_83_MT775572 | 23403AtoG | D | G | Nonsynonymous | 1529 |
| DNAS_isl_83_MT775572 | 10761AtoG | K | R | Nonsynonymous | 1891 |
| DNAS_isl_83_MT775572 | 290AtoT | N | C | Nonsynonymous | 3 |
| DNAS_isl_83_MT775572 | 291AtoG | N | C | Nonsynonymous | 3 |
| DNAS_isl_83_MT775572 | 25135GtoT | K | N | Nonsynonymous | 775 |
| DNAS_isl_83_MT775572 | 3037CtoT | F | F | Synonymous | 1591 |
| DNAS_isl_83_MT775572 | 1163AtoT | I | F | Nonsynonymous | 1414 |
| DNAS_isl_83_MT775572 | 3308GtoA | E | K | Nonsynonymous | 269 |
| DNAS_isl_83_MT775572 | 241CtoT | R | C | 5'UTR SNP | 299 |
| DNAS_isl_83_MT775572 | 28883GtoC | G | R | Nonsynonymous | 1391 |
| DNAS_isl_83_MT775572 | 28882GtoA | R | K | Nonsynonymous | 1391 |
| DNAS_isl_83_MT775572 | 29296CtoT | D | D | Synonymous | 486 |
| DNAS_isl_83_MT775572 | 14408CtoT | P | L | Nonsynonymous | 244 |
| DNAS_isl_83_MT775572 | 254CtoT | T | I | 5'UTR SNP | 297 |
| DNAS_isl_83_MT775572 | 28881GtoA | R | K | Nonsynonymous | 1386 |
| DNAS_isl_84_MT775569 | 11521GtoT | M | I | Nonsynonymous | 3126 |
| DNAS_isl_84_MT775569 | 5284CtoT | N | N | Synonymous | 7693 |
| DNAS_isl_84_MT775569 | 3037CtoT | F | F | Synonymous | 6482 |
| DNAS_isl_84_MT775569 | 1079AtoC | N | H | Nonsynonymous | 5083 |
| DNAS_isl_84_MT775569 | 10268AtoG | N | D | Nonsynonymous | 7970 |
| DNAS_isl_84_MT775569 | 23403AtoG | D | G | Nonsynonymous | 7955 |
| DNAS_isl_84_MT775569 | 241CtoT | R | C | 5'UTR SNP | 2470 |
| DNAS_isl_84_MT775569 | 23563TtoC | G | G | Synonymous | 6482 |
| DNAS_isl_84_MT775569 | 28881GtoA | R | K | Nonsynonymous | 4248 |
| DNAS_isl_84_MT775569 | 28883GtoC | G | R | Nonsynonymous | 4254 |
| DNAS_isl_84_MT775569 | 14408CtoT | P | L | Nonsynonymous | 2427 |
| DNAS_isl_84_MT775569 | 28882GtoA | R | K | Nonsynonymous | 4254 |
| DNAS_isl_85_MT775559 | 3037CtoT | F | F | Synonymous | 1803 |
| DNAS_isl_85_MT775559 | 23403AtoG | D | G | Nonsynonymous | 2075 |
| DNAS_isl_85_MT775559 | 925CtoT | D | D | Synonymous | 1764 |
| DNAS_isl_85_MT775559 | 1163AtoT | I | F | Nonsynonymous | 1071 |
| DNAS_isl_85_MT775559 | 19344TtoC | A | A | Synonymous | 1188 |
| DNAS_isl_85_MT775559 | 28881GtoA | R | K | Nonsynonymous | 1119 |
| DNAS_isl_85_MT775559 | 28883GtoC | G | R | Nonsynonymous | 1124 |
| DNAS_isl_85_MT775559 | 8146CtoT | D | D | Synonymous | 1027 |
| DNAS_isl_85_MT775559 | 28882GtoA | R | K | Nonsynonymous | 1124 |
| DNAS_isl_85_MT775559 | 14408CtoT | P | L | Nonsynonymous | 195 |
| DNAS_isl_85_MT775559 | 17518CtoT | L | F | Nonsynonymous | 2953 |
| DNAS_isl_85_MT775559 | 241CtoT | R | C | 5'UTR SNP | 323 |
| DNAS_isl_86_MT775561 | 14585CtoT | A | V | Nonsynonymous | 3255 |
| DNAS_isl_86_MT775561 | 1163AtoT | I | F | Nonsynonymous | 7274 |
| DNAS_isl_86_MT775561 | 19185CtoT | C | C | Synonymous | 1454 |
| DNAS_isl_86_MT775561 | 241CtoT | R | C | 5'UTR SNP | 2913 |
| DNAS_isl_86_MT775561 | 3037CtoT | F | F | Synonymous | 5757 |
| DNAS_isl_86_MT775561 | 28881GtoA | R | K | Nonsynonymous | 3882 |
| DNAS_isl_86_MT775561 | 23403AtoG | D | G | Nonsynonymous | 7959 |
| DNAS_isl_86_MT775561 | 28253CtoT | F | F | Synonymous | 7905 |
| DNAS_isl_86_MT775561 | 3994GtoT | L | L | Synonymous | 1416 |
| DNAS_isl_86_MT775561 | 28882GtoA | R | K | Nonsynonymous | 3894 |
| DNAS_isl_86_MT775561 | 21010GtoT | V | L | Nonsynonymous | 4174 |
| DNAS_isl_86_MT775561 | 14408CtoT | P | L | Nonsynonymous | 2215 |
| DNAS_isl_86_MT775561 | 28883GtoC | G | R | Nonsynonymous | 3896 |
| DNAS_isl_86_MT775561 | 3961CtoT | I | I | Synonymous | 2203 |
| DNAS_isl_87_MT745761 | 28292CtoA | Q | K | Nonsynonymous | 5040 |
| DNAS_isl_87_MT745761 | 23403AtoG | D | G | Nonsynonymous | 3208 |
| DNAS_isl_87_MT745761 | 1163AtoT | I | F | Nonsynonymous | 1651 |
| DNAS_isl_87_MT745761 | 28881GtoA | R | K | Nonsynonymous | 1820 |
| DNAS_isl_87_MT745761 | 28883GtoC | G | R | Nonsynonymous | 1832 |
| DNAS_isl_87_MT745761 | 4300GtoT | V | V | Synonymous | 2315 |
| DNAS_isl_87_MT745761 | 14408CtoT | P | L | Nonsynonymous | 774 |
| DNAS_isl_87_MT745761 | 28882GtoA | R | K | Nonsynonymous | 1832 |
| DNAS_isl_87_MT745761 | 241CtoT | R | C | 5'UTR SNP | 684 |
| DNAS_isl_87_MT745761 | 3037CtoT | F | F | Synonymous | 3019 |
| DNAS_isl_87_MT745761 | 26526GtoT | A | S | Nonsynonymous | 1817 |
| DNAS_isl_87_MT745761 | 17193GtoT | E | D | Nonsynonymous | 5299 |
| DNAS_isl_88_MT745755 | 23403AtoG | D | G | Nonsynonymous | 3970 |
| DNAS_isl_88_MT745755 | 241CtoT | R | C | 5'UTR SNP | 792 |
| DNAS_isl_88_MT745755 | 10285TtoC | V | V | Synonymous | 4877 |
| DNAS_isl_88_MT745755 | 1163AtoT | I | F | Nonsynonymous | 2879 |
| DNAS_isl_88_MT745755 | 22711TtoG | S | S | Synonymous | 193 |
| DNAS_isl_88_MT745755 | 28882GtoA | R | K | Nonsynonymous | 1777 |
| DNAS_isl_88_MT745755 | 28881GtoA | R | K | Nonsynonymous | 1764 |
| DNAS_isl_88_MT745755 | 28883GtoC | G | R | Nonsynonymous | 1777 |
| DNAS_isl_88_MT745755 | 25954GtoT | G | C | Nonsynonymous | 2254 |
| DNAS_isl_88_MT745755 | 6262TtoC | N | N | Synonymous | 6593 |
| DNAS_isl_88_MT745755 | 14408CtoT | P | L | Nonsynonymous | 681 |
| DNAS_isl_88_MT745755 | 421CtoT | G | G | Synonymous | 2640 |
| DNAS_isl_88_MT745755 | 3037CtoT | F | F | Synonymous | 3926 |
| DNAS_isl_89_MT745754 | 23403AtoG | D | G | Nonsynonymous | 7964 |
| DNAS_isl_89_MT745754 | 10323AtoG | K | R | Nonsynonymous | 7993 |
| DNAS_isl_89_MT745754 | 1170CtoT | S | F | Nonsynonymous | 7645 |
| DNAS_isl_89_MT745754 | 8692CtoT | Y | Y | Synonymous | 2803 |
| DNAS_isl_89_MT745754 | 14408CtoT | P | L | Nonsynonymous | 3566 |
| DNAS_isl_89_MT745754 | 28882GtoA | R | K | Nonsynonymous | 3612 |
| DNAS_isl_89_MT745754 | 28883GtoC | G | R | Nonsynonymous | 3611 |
| DNAS_isl_89_MT745754 | 28808GtoA | G | S | Nonsynonymous | 7997 |
| DNAS_isl_89_MT745754 | 1163AtoT | I | F | Nonsynonymous | 7694 |
| DNAS_isl_89_MT745754 | 29632CtoT | N | N | Synonymous | 7903 |
| DNAS_isl_89_MT745754 | 28881GtoA | R | K | Nonsynonymous | 3599 |
| DNAS_isl_89_MT745754 | 3037CtoT | F | F | Synonymous | 5718 |
| DNAS_isl_89_MT745754 | 241CtoT | R | C | 5'UTR SNP | 1539 |
| DNAS_isl_90_MT745756 | 28882GtoA | R | K | Nonsynonymous | 5166 |
| DNAS_isl_90_MT745756 | 28881GtoA | R | K | Nonsynonymous | 5163 |
| DNAS_isl_90_MT745756 | 28883GtoC | G | R | Nonsynonymous | 5166 |
| DNAS_isl_90_MT745756 | 14408CtoT | P | L | Nonsynonymous | 1017 |
| DNAS_isl_90_MT745756 | 241CtoT | R | C | 5'UTR SNP | 3209 |
| DNAS_isl_90_MT745756 | 8128AtoC | A | A | Synonymous | 7987 |
| DNAS_isl_90_MT745756 | 27806GtoT | L | L | Synonymous | 7974 |
| DNAS_isl_90_MT745756 | 2967GtoT | S | I | Nonsynonymous | 7523 |
| DNAS_isl_90_MT745756 | 3037CtoT | F | F | Synonymous | 5584 |
| DNAS_isl_90_MT745756 | 23403AtoG | D | G | Nonsynonymous | 7964 |
| DNAS_isl_90_MT745756 | 1163AtoT | I | F | Nonsynonymous | 3223 |
| DNAS_isl_91_MT745765 | 241CtoT | R | C | 5'UTR SNP | 725 |
| DNAS_isl_91_MT745765 | 3037CtoT | F | F | Synonymous | 3887 |
| DNAS_isl_91_MT745765 | 3961CtoT | I | I | Synonymous | 2158 |
| DNAS_isl_91_MT745765 | 774CtoT | T | I | Nonsynonymous | 1220 |
| DNAS_isl_91_MT745765 | 13902TtoC | D | D | Synonymous | 1985 |
| DNAS_isl_91_MT745765 | 28882GtoA | R | K | Nonsynonymous | 2649 |
| DNAS_isl_91_MT745765 | 1163AtoT | I | F | Nonsynonymous | 2272 |
| DNAS_isl_91_MT745765 | 21123GtoT | G | G | Synonymous | 4932 |
| DNAS_isl_91_MT745765 | 28881GtoA | R | K | Nonsynonymous | 2636 |
| DNAS_isl_91_MT745765 | 23403AtoG | D | G | Nonsynonymous | 5315 |
| DNAS_isl_91_MT745765 | 22199GtoT | V | L | Nonsynonymous | 1524 |
| DNAS_isl_91_MT745765 | 14408CtoT | P | L | Nonsynonymous | 1041 |
| DNAS_isl_91_MT745765 | 28883GtoC | G | R | Nonsynonymous | 2652 |
| DNAS_isl_91_MT745765 | 19593CtoT | N | N | Synonymous | 4171 |
| DNAS_isl_91_MT745765 | 21597CtoT | S | F | Nonsynonymous | 977 |
| DNAS_isl_92_MT742762 | 774CtoT | T | I | Nonsynonymous | 3408 |
| DNAS_isl_92_MT742762 | 14408CtoT | P | L | Nonsynonymous | 3929 |
| DNAS_isl_92_MT742762 | 22199GtoT | V | L | Nonsynonymous | 4399 |
| DNAS_isl_92_MT742762 | 3037CtoT | F | F | Synonymous | 4802 |
| DNAS_isl_92_MT742762 | 28881GtoA | R | K | Nonsynonymous | 4424 |
| DNAS_isl_92_MT742762 | 28882GtoA | R | K | Nonsynonymous | 4436 |
| DNAS_isl_92_MT742762 | 28883GtoC | G | R | Nonsynonymous | 4437 |
| DNAS_isl_92_MT742762 | 21123GtoT | G | G | Synonymous | 7872 |
| DNAS_isl_92_MT742762 | 3961CtoT | I | I | Synonymous | 5553 |
| DNAS_isl_92_MT742762 | 21597CtoT | S | F | Nonsynonymous | 2746 |
| DNAS_isl_92_MT742762 | 1163AtoT | I | F | Nonsynonymous | 7739 |
| DNAS_isl_92_MT742762 | 19593CtoT | N | N | Synonymous | 7956 |
| DNAS_isl_92_MT742762 | 241CtoT | R | C | 5'UTR SNP | 3958 |
| DNAS_isl_92_MT742762 | 13902TtoC | D | D | Synonymous | 8017 |
| DNAS_isl_92_MT742762 | 23403AtoG | D | G | Nonsynonymous | 7935 |
| DNAS_isl_93_MT745757 | 23403AtoG | D | G | Nonsynonymous | 7972 |
| DNAS_isl_93_MT745757 | 241CtoT | R | C | 5'UTR SNP | 4265 |
| DNAS_isl_93_MT745757 | 3961CtoT | I | I | Synonymous | 5778 |
| DNAS_isl_93_MT745757 | 3037CtoT | F | F | Synonymous | 5390 |
| DNAS_isl_93_MT745757 | 19593CtoT | N | N | Synonymous | 7939 |
| DNAS_isl_93_MT745757 | 13902TtoC | D | D | Synonymous | 7998 |
| DNAS_isl_93_MT745757 | 28881GtoA | R | K | Nonsynonymous | 3567 |
| DNAS_isl_93_MT745757 | 22199GtoT | V | L | Nonsynonymous | 4723 |
| DNAS_isl_93_MT745757 | 28882GtoA | R | K | Nonsynonymous | 3572 |
| DNAS_isl_93_MT745757 | 28883GtoC | G | R | Nonsynonymous | 3572 |
| DNAS_isl_93_MT745757 | 14408CtoT | P | L | Nonsynonymous | 4269 |
| DNAS_isl_93_MT745757 | 774CtoT | T | I | Nonsynonymous | 2116 |
| DNAS_isl_93_MT745757 | 1163AtoT | I | F | Nonsynonymous | 5539 |
| DNAS_isl_93_MT745757 | 21123GtoT | G | G | Synonymous | 7847 |
| DNAS_isl_93_MT745757 | 21597CtoT | S | F | Nonsynonymous | 3287 |
| DNAS_isl_94_MT745760 | 241CtoT | R | C | 5'UTR SNP | 3117 |
| DNAS_isl_94_MT745760 | 1163AtoT | I | F | Nonsynonymous | 4687 |
| DNAS_isl_94_MT745760 | 28882GtoA | R | K | Nonsynonymous | 5217 |
| DNAS_isl_94_MT745760 | 28883GtoC | G | R | Nonsynonymous | 5218 |
| DNAS_isl_94_MT745760 | 28881GtoA | R | K | Nonsynonymous | 5209 |
| DNAS_isl_94_MT745760 | 23403AtoG | D | G | Nonsynonymous | 7976 |
| DNAS_isl_94_MT745760 | 14408CtoT | P | L | Nonsynonymous | 4205 |
| DNAS_isl_94_MT745760 | 3037CtoT | F | F | Synonymous | 4767 |
| DNAS_isl_95_MT745763 | 28882GtoA | R | K | Nonsynonymous | 1086 |
| DNAS_isl_95_MT745763 | 241CtoT | R | C | 5'UTR SNP | 1059 |
| DNAS_isl_95_MT745763 | 28883GtoC | G | R | Nonsynonymous | 1086 |
| DNAS_isl_95_MT745763 | 3037CtoT | F | F | Synonymous | 5820 |
| DNAS_isl_95_MT745763 | 28881GtoA | R | K | Nonsynonymous | 1082 |
| DNAS_isl_95_MT745763 | 14408CtoT | P | L | Nonsynonymous | 1489 |
| DNAS_isl_95_MT745763 | 1163AtoT | I | F | Nonsynonymous | 2751 |
| DNAS_isl_95_MT745763 | 23403AtoG | D | G | Nonsynonymous | 5954 |
| DNAS_isl_96_MT745753 | 1163AtoT | I | F | Nonsynonymous | 7538 |
| DNAS_isl_96_MT745753 | 22033CtoA | F | L | Nonsynonymous | 7656 |
| DNAS_isl_96_MT745753 | 28883GtoC | G | R | Nonsynonymous | 2774 |
| DNAS_isl_96_MT745753 | 28882GtoA | R | K | Nonsynonymous | 2773 |
| DNAS_isl_96_MT745753 | 28881GtoA | R | K | Nonsynonymous | 2769 |
| DNAS_isl_96_MT745753 | 25228TtoG | A | A | Synonymous | 5184 |
| DNAS_isl_96_MT745753 | 241CtoT | R | C | 5'UTR SNP | 1601 |
| DNAS_isl_96_MT745753 | 14408CtoT | P | L | Nonsynonymous | 4454 |
| DNAS_isl_96_MT745753 | 3037CtoT | F | F | Synonymous | 4575 |
| DNAS_isl_96_MT745753 | 23403AtoG | D | G | Nonsynonymous | 7942 |
| DNAS_isl_97_MT745752 | 1403CtoT | P | S | Nonsynonymous | 7867 |
| DNAS_isl_97_MT745752 | 8502CtoT | T | I | Nonsynonymous | 4739 |
| DNAS_isl_97_MT745752 | 3037CtoT | F | F | Synonymous | 4664 |
| DNAS_isl_97_MT745752 | 10153TtoC | D | D | Synonymous | 7243 |
| DNAS_isl_97_MT745752 | 28881GtoA | R | K | Nonsynonymous | 3097 |
| DNAS_isl_97_MT745752 | 18412GtoT | V | F | Nonsynonymous | 3923 |
| DNAS_isl_97_MT745752 | 1163AtoT | I | F | Nonsynonymous | 7641 |
| DNAS_isl_97_MT745752 | 23403AtoG | D | G | Nonsynonymous | 7966 |
| DNAS_isl_97_MT745752 | 28883GtoC | G | R | Nonsynonymous | 3101 |
| DNAS_isl_97_MT745752 | 14408CtoT | P | L | Nonsynonymous | 5107 |
| DNAS_isl_97_MT745752 | 28882GtoA | R | K | Nonsynonymous | 3101 |
| DNAS_isl_97_MT745752 | 241CtoT | R | C | 5'UTR SNP | 1435 |
| DNAS_isl_97_MT745752 | 16905TtoC | D | D | Synonymous | 7624 |
| DNAS_isl_98_MT745764 | 14408CtoT | P | L | Nonsynonymous | 2210 |
| DNAS_isl_98_MT745764 | 4503AtoT | E | V | Nonsynonymous | 2197 |
| DNAS_isl_98_MT745764 | 28727GtoT | A | S | Nonsynonymous | 7793 |
| DNAS_isl_98_MT745764 | 23403AtoG | D | G | Nonsynonymous | 7976 |
| DNAS_isl_98_MT745764 | 28881GtoA | R | K | Nonsynonymous | 4497 |
| DNAS_isl_98_MT745764 | 1163AtoT | I | F | Nonsynonymous | 4382 |
| DNAS_isl_98_MT745764 | 28882GtoA | R | K | Nonsynonymous | 4511 |
| DNAS_isl_98_MT745764 | 241CtoT | R | C | 5'UTR SNP | 1825 |
| DNAS_isl_98_MT745764 | 28883GtoC | G | Q | Nonsynonymous | 4514 |
| DNAS_isl_98_MT745764 | 3037CtoT | F | F | Synonymous | 5285 |
| DNAS_isl_98_MT745764 | 28884GtoA | G | Q | Nonsynonymous | 4514 |
| DNAS_isl_99_MT745759 | 14408CtoT | P | L | Nonsynonymous | 595 |
| DNAS_isl_99_MT745759 | 3037CtoT | F | F | Synonymous | 1918 |
| DNAS_isl_99_MT745759 | 241CtoT | R | C | 5'UTR SNP | 490 |
| DNAS_isl_99_MT745759 | 6154AtoG | L | L | Synonymous | 3352 |
| DNAS_isl_99_MT745759 | 28882GtoA | R | K | Nonsynonymous | 1668 |
| DNAS_isl_99_MT745759 | 28883GtoC | G | R | Nonsynonymous | 1669 |
| DNAS_isl_99_MT745759 | 1163AtoT | I | F | Nonsynonymous | 2332 |
| DNAS_isl_99_MT745759 | 23403AtoG | D | G | Nonsynonymous | 2447 |
| DNAS_isl_99_MT745759 | 28881GtoA | R | K | Nonsynonymous | 1653 |
| DNAS_isl_100_MT745762 | 1163AtoT | I | F | Nonsynonymous | 1412 |
| DNAS_isl_100_MT745762 | 14408CtoT | P | L | Nonsynonymous | 343 |
| DNAS_isl_100_MT745762 | 6154AtoG | L | L | Synonymous | 2546 |
| DNAS_isl_100_MT745762 | 3037CtoT | F | F | Synonymous | 1629 |
| DNAS_isl_100_MT745762 | 241CtoT | R | C | 5'UTR SNP | 582 |
| DNAS_isl_100_MT745762 | 23403AtoG | D | G | Nonsynonymous | 2154 |
| DNAS_isl_100_MT745762 | 28882GtoA | R | K | Nonsynonymous | 1400 |
| DNAS_isl_100_MT745762 | 28883GtoC | G | R | Nonsynonymous | 1400 |
| DNAS_isl_100_MT745762 | 28881GtoA | R | K | Nonsynonymous | 1390 |
| DNAS_isl_101_MT742761 | 14408CtoT | P | L | Nonsynonymous | 307 |
| DNAS_isl_101_MT742761 | 16943GtoT | S | I | Nonsynonymous | 2037 |
| DNAS_isl_101_MT742761 | 27972CtoT | Q | * | Nonsynonymous | 3937 |
| DNAS_isl_101_MT742761 | 1163AtoT | I | F | Nonsynonymous | 1050 |
| DNAS_isl_101_MT742761 | 23403AtoG | D | G | Nonsynonymous | 1371 |
| DNAS_isl_101_MT742761 | 3961CtoT | I | I | Synonymous | 653 |
| DNAS_isl_101_MT742761 | 3037CtoT | F | F | Synonymous | 1404 |
| DNAS_isl_101_MT742761 | 28883GtoC | G | R | Nonsynonymous | 1176 |
| DNAS_isl_101_MT742761 | 28882GtoA | R | K | Nonsynonymous | 1176 |
| DNAS_isl_101_MT742761 | 241CtoT | R | C | 5'UTR SNP | 369 |
| DNAS_isl_101_MT742761 | 28881GtoA | R | K | Nonsynonymous | 1174 |
| DNAS_isl_101_MT742761 | 29659CtoT | N | N | Synonymous | 4111 |
| DNAS_isl_102_MT745750 | 3037CtoT | F | F | Synonymous | 5299 |
| DNAS_isl_102_MT745750 | 1163AtoT | I | F | Nonsynonymous | 5526 |
| DNAS_isl_102_MT745750 | 19525GtoT | D | Y | Nonsynonymous | 4267 |
| DNAS_isl_102_MT745750 | 3961CtoT | I | I | Synonymous | 4012 |
| DNAS_isl_102_MT745750 | 23403AtoG | D | G | Nonsynonymous | 7961 |
| DNAS_isl_102_MT745750 | 28882GtoA | R | K | Nonsynonymous | 3809 |
| DNAS_isl_102_MT745750 | 241CtoT | R | C | 5'UTR SNP | 3089 |
| DNAS_isl_102_MT745750 | 28881GtoA | R | K | Nonsynonymous | 3801 |
| DNAS_isl_102_MT745750 | 28883GtoC | G | R | Nonsynonymous | 3809 |
| DNAS_isl_102_MT745750 | 14408CtoT | P | L | Nonsynonymous | 3626 |
| DNAS_isl_102_MT745750 | 26110CtoT | P | S | Nonsynonymous | 8005 |
| DNAS_isl_102_MT745750 | 1722CtoT | A | V | Nonsynonymous | 1678 |
| DNAS_isl_103_MT745751 | 1163AtoT | I | F | Nonsynonymous | 7442 |
| DNAS_isl_103_MT745751 | 28883GtoC | G | R | Nonsynonymous | 2029 |
| DNAS_isl_103_MT745751 | 21897CtoT | S | L | Nonsynonymous | 7973 |
| DNAS_isl_103_MT745751 | 3037CtoT | F | F | Synonymous | 4686 |
| DNAS_isl_103_MT745751 | 12301GtoA | M | I | Nonsynonymous | 7997 |
| DNAS_isl_103_MT745751 | 23403AtoG | D | G | Nonsynonymous | 7964 |
| DNAS_isl_103_MT745751 | 21641GtoT | A | S | Nonsynonymous | 7854 |
| DNAS_isl_103_MT745751 | 28881GtoA | R | K | Nonsynonymous | 2012 |
| DNAS_isl_103_MT745751 | 14408CtoT | P | L | Nonsynonymous | 4197 |
| DNAS_isl_103_MT745751 | 28882GtoA | R | K | Nonsynonymous | 2029 |
| DNAS_isl_103_MT745751 | 3961CtoT | I | I | Synonymous | 7680 |
| DNAS_isl_103_MT745751 | 25311GtoT | C | F | Nonsynonymous | 7836 |
| DNAS_isl_103_MT745751 | 8860CtoT | V | V | Synonymous | 7963 |
| DNAS_isl_103_MT745751 | 241CtoT | R | C | 5'UTR SNP | 1453 |
| DNAS_isl_104_MT745758 | 241CtoT | R | C | 5'UTR SNP | 213 |
| DNAS_isl_104_MT745758 | 28883GtoC | G | R | Nonsynonymous | 530 |
| DNAS_isl_104_MT745758 | 28882GtoA | R | K | Nonsynonymous | 530 |
| DNAS_isl_104_MT745758 | 28881GtoA | R | K | Nonsynonymous | 523 |
| DNAS_isl_104_MT745758 | 28200TtoC | S | P | Nonsynonymous | 486 |
| DNAS_isl_104_MT745758 | 19648GtoT | V | L | Nonsynonymous | 674 |
| DNAS_isl_104_MT745758 | 23403AtoG | D | G | Nonsynonymous | 932 |
| DNAS_isl_104_MT745758 | 23868GtoT | G | V | Nonsynonymous | 330 |
| DNAS_isl_104_MT745758 | 3037CtoT | F | F | Synonymous | 923 |
| DNAS_isl_104_MT745758 | 1163AtoT | I | F | Nonsynonymous | 523 |
| DNAS_isl_104_MT745758 | 26049GtoT | L | F | Nonsynonymous | 213 |
| DNAS_isl_104_MT745758 | 14408CtoT | P | L | Nonsynonymous | 123 |
| DNAS_isl_105_MT676415 | 26094TtoC | N | N | Synonymous | 4297 |
| DNAS_isl_105_MT676415 | 22432CtoT | D | D | Synonymous | 1127 |
| DNAS_isl_105_MT676415 | 25844CtoT | T | I | Nonsynonymous | 859 |
| DNAS_isl_105_MT676415 | 22801GtoT | G | G | Synonymous | 1073 |
| DNAS_isl_105_MT676415 | 241CtoT | R | C | 5'UTR SNP | 562 |
| DNAS_isl_105_MT676415 | 28881GtoA | R | K | Nonsynonymous | 2253 |
| DNAS_isl_105_MT676415 | 1163AtoT | I | F | Nonsynonymous | 2022 |
| DNAS_isl_105_MT676415 | 28882GtoA | R | K | Nonsynonymous | 2260 |
| DNAS_isl_105_MT676415 | 28883GtoC | G | R | Nonsynonymous | 2260 |
| DNAS_isl_105_MT676415 | 3037CtoT | F | F | Synonymous | 1604 |
| DNAS_isl_105_MT676415 | 23403AtoG | D | G | Nonsynonymous | 2498 |
| DNAS_isl_106_MT676414 | 28883GtoC | G | R | Nonsynonymous | 4057 |
| DNAS_isl_106_MT676414 | 28881GtoA | R | K | Nonsynonymous | 4052 |
| DNAS_isl_106_MT676414 | 28882GtoA | R | K | Nonsynonymous | 4057 |
| DNAS_isl_106_MT676414 | 1163AtoT | I | F | Nonsynonymous | 6800 |
| DNAS_isl_106_MT676414 | 241CtoT | R | C | 5'UTR SNP | 1195 |
| DNAS_isl_106_MT676414 | 3037CtoT | F | F | Synonymous | 5280 |
| DNAS_isl_106_MT676414 | 14408CtoT | P | L | Nonsynonymous | 4072 |
| DNAS_isl_106_MT676414 | 23403AtoG | D | G | Nonsynonymous | 7972 |
| DNAS_isl_107_MT676413 | 20438GtoT | S | I | Nonsynonymous | 7982 |
| DNAS_isl_107_MT676413 | 23403AtoG | D | G | Nonsynonymous | 7988 |
| DNAS_isl_107_MT676413 | 14408CtoT | P | L | Nonsynonymous | 3032 |
| DNAS_isl_107_MT676413 | 241CtoT | R | C | 5'UTR SNP | 2012 |
| DNAS_isl_107_MT676413 | 4201GtoT | M | I | Nonsynonymous | 5812 |
| DNAS_isl_107_MT676413 | 3037CtoT | F | F | Synonymous | 6131 |
| DNAS_isl_107_MT676413 | 12369CtoT | T | I | Nonsynonymous | 5139 |
| DNAS_isl_107_MT676413 | 26527CtoT | A | V | Nonsynonymous | 3737 |
| DNAS_isl_108_MT676418 | 28881GtoA | R | K | Nonsynonymous | 4974 |
| DNAS_isl_108_MT676418 | 28882GtoA | R | K | Nonsynonymous | 4979 |
| DNAS_isl_108_MT676418 | 28883GtoC | G | R | Nonsynonymous | 4979 |
| DNAS_isl_108_MT676418 | 241CtoT | R | C | 5'UTR SNP | 1383 |
| DNAS_isl_108_MT676418 | 1163AtoT | I | F | Nonsynonymous | 5472 |
| DNAS_isl_108_MT676418 | 4331CtoT | L | L | Synonymous | 4741 |
| DNAS_isl_108_MT676418 | 3037CtoT | F | F | Synonymous | 4782 |
| DNAS_isl_108_MT676418 | 23403AtoG | D | G | Nonsynonymous | 7924 |
| DNAS_isl_108_MT676418 | 19101GtoA | Q | Q | Synonymous | 5570 |
| DNAS_isl_109_MT676420 | 14408CtoT | P | L | Nonsynonymous | 5242 |
| DNAS_isl_109_MT676420 | 1163AtoT | I | F | Nonsynonymous | 7097 |
| DNAS_isl_109_MT676420 | 19101GtoA | Q | Q | Synonymous | 7912 |
| DNAS_isl_109_MT676420 | 241CtoT | R | C | 5'UTR SNP | 1403 |
| DNAS_isl_109_MT676420 | 28881GtoA | R | K | Nonsynonymous | 3167 |
| DNAS_isl_109_MT676420 | 28883GtoC | G | R | Nonsynonymous | 3175 |
| DNAS_isl_109_MT676420 | 23403AtoG | D | G | Nonsynonymous | 8015 |
| DNAS_isl_109_MT676420 | 28882GtoA | R | K | Nonsynonymous | 3175 |
| DNAS_isl_109_MT676420 | 3037CtoT | F | F | Synonymous | 4811 |
| DNAS_isl_110_MT676416 | 23403AtoG | D | G | Nonsynonymous | 7983 |
| DNAS_isl_110_MT676416 | 8560TtoC | I | I | Synonymous | 7994 |
| DNAS_isl_110_MT676416 | 3037CtoT | F | F | Synonymous | 6136 |
| DNAS_isl_110_MT676416 | 1163AtoT | I | F | Nonsynonymous | 6365 |
| DNAS_isl_110_MT676416 | 29421CtoT | P | L | Nonsynonymous | 7859 |
| DNAS_isl_110_MT676416 | 241CtoT | R | C | 5'UTR SNP | 1131 |
| DNAS_isl_110_MT676416 | 28881GtoA | R | K | Nonsynonymous | 3509 |
| DNAS_isl_110_MT676416 | 5800GtoT | L | F | Nonsynonymous | 7917 |
| DNAS_isl_110_MT676416 | 28883GtoC | G | R | Nonsynonymous | 3517 |
| DNAS_isl_110_MT676416 | 28882GtoA | R | K | Nonsynonymous | 3516 |
| DNAS_isl_110_MT676416 | 14408CtoT | P | L | Nonsynonymous | 3728 |
| DNAS_isl_111_MT676412 | 23403AtoG | D | G | Nonsynonymous | 7925 |
| DNAS_isl_111_MT676412 | 19101GtoA | Q | Q | Synonymous | 7905 |
| DNAS_isl_111_MT676412 | 1163AtoT | I | F | Nonsynonymous | 6938 |
| DNAS_isl_111_MT676412 | 241CtoT | R | C | 5'UTR SNP | 1151 |
| DNAS_isl_111_MT676412 | 3037CtoT | F | F | Synonymous | 5285 |
| DNAS_isl_111_MT676412 | 28881GtoA | R | K | Nonsynonymous | 3573 |
| DNAS_isl_111_MT676412 | 14408CtoT | P | L | Nonsynonymous | 4429 |
| DNAS_isl_111_MT676412 | 28883GtoC | G | R | Nonsynonymous | 3580 |
| DNAS_isl_111_MT676412 | 28882GtoA | R | K | Nonsynonymous | 3580 |
| DNAS_isl_112_MT676419 | 28882GtoA | R | K | Nonsynonymous | 4809 |
| DNAS_isl_112_MT676419 | 28881GtoA | R | K | Nonsynonymous | 4799 |
| DNAS_isl_112_MT676419 | 3037CtoT | F | F | Synonymous | 4485 |
| DNAS_isl_112_MT676419 | 241CtoT | R | C | 5'UTR SNP | 1471 |
| DNAS_isl_112_MT676419 | 28883GtoC | G | R | Nonsynonymous | 4809 |
| DNAS_isl_112_MT676419 | 23403AtoG | D | G | Nonsynonymous | 7854 |
| DNAS_isl_112_MT676419 | 1163AtoT | I | F | Nonsynonymous | 5454 |
| DNAS_isl_112_MT676419 | 24764GtoT | V | F | Nonsynonymous | 5059 |
| DNAS_isl_112_MT676419 | 28378GtoC | A | A | Synonymous | 5250 |
| DNAS_isl_112_MT676419 | 27769CtoT | S | L | Nonsynonymous | 3459 |
| DNAS_isl_113_MT676421 | 21691CtoT | F | F | Synonymous | 5501 |
| DNAS_isl_113_MT676421 | 2036GtoT | A | S | Nonsynonymous | 7970 |
| DNAS_isl_113_MT676421 | 1163AtoT | I | F | Nonsynonymous | 5789 |
| DNAS_isl_113_MT676421 | 3037CtoT | F | F | Synonymous | 4784 |
| DNAS_isl_113_MT676421 | 14408CtoT | P | L | Nonsynonymous | 5383 |
| DNAS_isl_113_MT676421 | 22000CtoG | H | Q | Nonsynonymous | 5476 |
| DNAS_isl_113_MT676421 | 28881GtoA | R | K | Nonsynonymous | 2942 |
| DNAS_isl_113_MT676421 | 12084CtoT | T | I | Nonsynonymous | 7571 |
| DNAS_isl_113_MT676421 | 241CtoT | R | C | 5'UTR SNP | 1251 |
| DNAS_isl_113_MT676421 | 23403AtoG | D | G | Nonsynonymous | 7937 |
| DNAS_isl_113_MT676421 | 28882GtoA | R | K | Nonsynonymous | 2950 |
| DNAS_isl_113_MT676421 | 28883GtoC | G | R | Nonsynonymous | 2951 |
| DNAS_isl_114_MT676417 | 3037CtoT | F | F | Synonymous | 6503 |
| DNAS_isl_114_MT676417 | 1163AtoT | I | F | Nonsynonymous | 7341 |
| DNAS_isl_114_MT676417 | 23403AtoG | D | G | Nonsynonymous | 7986 |
| DNAS_isl_114_MT676417 | 241CtoT | R | C | 5'UTR SNP | 2900 |
| DNAS_isl_114_MT676417 | 28881GtoA | R | K | Nonsynonymous | 3500 |
| DNAS_isl_114_MT676417 | 28882GtoA | R | K | Nonsynonymous | 3505 |
| DNAS_isl_114_MT676417 | 14408CtoT | P | L | Nonsynonymous | 3572 |
| DNAS_isl_114_MT676417 | 28883GtoC | G | R | Nonsynonymous | 3505 |
| DNAS_isl_115_MT607976 | 241CtoT | R | C | 5'UTR SNP | 1047 |
| DNAS_isl_115_MT607976 | 28883GtoC | G | R | Nonsynonymous | 4967 |
| DNAS_isl_115_MT607976 | 1288CtoT | C | C | Synonymous | 4462 |
| DNAS_isl_115_MT607976 | 13255CtoT | C | C | Synonymous | 3440 |
| DNAS_isl_115_MT607976 | 28881GtoA | R | K | Nonsynonymous | 4939 |
| DNAS_isl_115_MT607976 | 28882GtoA | R | K | Nonsynonymous | 4967 |
| DNAS_isl_115_MT607976 | 23403AtoG | D | G | Nonsynonymous | 5531 |
| DNAS_isl_115_MT607976 | 3037CtoT | F | F | Synonymous | 3613 |
| DNAS_isl_115_MT607976 | 2048CtoT | L | L | Synonymous | 2452 |
| DNAS_isl_115_MT607976 | 22193AtoT | N | Y | Nonsynonymous | 1688 |
| DNAS_isl_115_MT607976 | 4752CtoT | T | I | Nonsynonymous | 2948 |
| DNAS_isl_116_MT607972 | 23403AtoG | D | G | Nonsynonymous | 7931 |
| DNAS_isl_116_MT607972 | 28883GtoC | G | R | Nonsynonymous | 3676 |
| DNAS_isl_116_MT607972 | 1163AtoT | I | F | Nonsynonymous | 7656 |
| DNAS_isl_116_MT607972 | 21987GtoT | G | V | Nonsynonymous | 2793 |
| DNAS_isl_116_MT607972 | 2644CtoT | I | I | Synonymous | 6747 |
| DNAS_isl_116_MT607972 | 28881GtoA | R | K | Nonsynonymous | 3665 |
| DNAS_isl_116_MT607972 | 14408CtoT | P | L | Nonsynonymous | 1939 |
| DNAS_isl_116_MT607972 | 28882GtoA | R | K | Nonsynonymous | 3677 |
| DNAS_isl_116_MT607972 | 241CtoT | R | C | 5'UTR SNP | 2127 |
| DNAS_isl_116_MT607972 | 27735CtoT | F | F | Synonymous | 1399 |
| DNAS_isl_116_MT607972 | 7173CtoT | S | F | Nonsynonymous | 1527 |
| DNAS_isl_116_MT607972 | 3037CtoT | F | F | Synonymous | 5470 |
| DNAS_isl_117_MT607975 | 16194TtoA | P | P | Synonymous | 7998 |
| DNAS_isl_117_MT607975 | 28883GtoC | G | R | Nonsynonymous | 3320 |
| DNAS_isl_117_MT607975 | 28882GtoA | R | K | Nonsynonymous | 3320 |
| DNAS_isl_117_MT607975 | 5175CtoT | T | I | Nonsynonymous | 7975 |
| DNAS_isl_117_MT607975 | 14408CtoT | P | L | Nonsynonymous | 4061 |
| DNAS_isl_117_MT607975 | 28881GtoA | R | K | Nonsynonymous | 3313 |
| DNAS_isl_117_MT607975 | 25617GtoT | K | N | Nonsynonymous | 7990 |
| DNAS_isl_117_MT607975 | 3037CtoT | F | F | Synonymous | 5508 |
| DNAS_isl_117_MT607975 | 1163AtoT | I | F | Nonsynonymous | 7145 |
| DNAS_isl_117_MT607975 | 241CtoT | R | C | 5'UTR SNP | 1422 |
| DNAS_isl_117_MT607975 | 19664TtoC | F | S | Nonsynonymous | 7987 |
| DNAS_isl_117_MT607975 | 23403AtoG | D | G | Nonsynonymous | 7979 |
| DNAS_isl_118_MT607974 | 14307TtoC | Y | Y | Synonymous | 7944 |
| DNAS_isl_118_MT607974 | 23403AtoG | D | G | Nonsynonymous | 7955 |
| DNAS_isl_118_MT607974 | 241CtoT | R | C | 5'UTR SNP | 2765 |
| DNAS_isl_118_MT607974 | 28881GtoA | R | K | Nonsynonymous | 3819 |
| DNAS_isl_118_MT607974 | 28882GtoA | R | K | Nonsynonymous | 3830 |
| DNAS_isl_118_MT607974 | 14408CtoT | P | L | Nonsynonymous | 4029 |
| DNAS_isl_118_MT607974 | 28883GtoC | G | R | Nonsynonymous | 3830 |
| DNAS_isl_118_MT607974 | 3037CtoT | F | F | Synonymous | 5824 |
| DNAS_isl_118_MT607974 | 5497CtoT | C | C | Synonymous | 7889 |
| DNAS_isl_118_MT607974 | 1288CtoT | C | C | Synonymous | 7896 |
| DNAS_isl_118_MT607974 | 22269CtoG | T | S | Nonsynonymous | 7936 |
| DNAS_isl_118_MT607974 | 2048CtoT | L | L | Synonymous | 7793 |
| DNAS_isl_118_MT607974 | 22193AtoT | N | Y | Nonsynonymous | 7771 |
| DNAS_isl_118_MT607974 | 13255CtoT | C | C | Synonymous | 4488 |
| DNAS_isl_119_MT607973 | 4201GtoT | M | I | Nonsynonymous | 1537 |
| DNAS_isl_119_MT607973 | 23403AtoG | D | G | Nonsynonymous | 3542 |
| DNAS_isl_119_MT607973 | 28391CtoT | R | C | Nonsynonymous | 2459 |
| DNAS_isl_119_MT607973 | 241CtoT | R | C | 5'UTR SNP | 635 |
| DNAS_isl_119_MT607973 | 26527CtoT | A | V | Nonsynonymous | 2008 |
| DNAS_isl_119_MT607973 | 17561GtoA | R | Q | Nonsynonymous | 4217 |
| DNAS_isl_119_MT607973 | 13923CtoT | D | D | Synonymous | 3131 |
| DNAS_isl_119_MT607973 | 3037CtoT | F | F | Synonymous | 2126 |
| DNAS_isl_119_MT607973 | 18756GtoT | P | P | Synonymous | 1042 |
| DNAS_isl_119_MT607973 | 17138CtoT | S | F | Nonsynonymous | 4812 |
| DNAS_isl_120_MT581414 | 14408CtoT | P | L | Nonsynonymous | 2807 |
| DNAS_isl_120_MT581414 | 28881GtoA | R | K | Nonsynonymous | 2283 |
| DNAS_isl_120_MT581414 | 28882GtoA | R | K | Nonsynonymous | 2288 |
| DNAS_isl_120_MT581414 | 241CtoT | R | C | 5'UTR SNP | 2286 |
| DNAS_isl_120_MT581414 | 28883GtoC | G | R | Nonsynonymous | 2288 |
| DNAS_isl_120_MT581414 | 27945CtoG | Q | E | Nonsynonymous | 4008 |
| DNAS_isl_120_MT581414 | 1163AtoT | I | F | Nonsynonymous | 5378 |
| DNAS_isl_120_MT581414 | 23403AtoG | D | G | Nonsynonymous | 7989 |
| DNAS_isl_120_MT581414 | 3037CtoT | F | F | Synonymous | 5907 |
| DNAS_isl_120_MT581414 | 2210GtoT | V | F | Nonsynonymous | 4215 |
| DNAS_isl_121_MT581434 | 8717CtoA | R | S | Nonsynonymous | 10 |
| DNAS_isl_121_MT581434 | 15324CtoT | N | N | Synonymous | 8015 |
| DNAS_isl_121_MT581434 | 3688CtoT | H | H | Synonymous | 4655 |
| DNAS_isl_121_MT581434 | 20896CtoG | P | A | Nonsynonymous | 8 |
| DNAS_isl_121_MT581434 | 23403AtoG | D | G | Nonsynonymous | 6098 |
| DNAS_isl_121_MT581434 | 3037CtoT | F | F | Synonymous | 4802 |
| DNAS_isl_121_MT581434 | 14408CtoT | P | L | Nonsynonymous | 6164 |
| DNAS_isl_121_MT581434 | 3077GtoA | E | K | Nonsynonymous | 4392 |
| DNAS_isl_121_MT581434 | 8716TtoG | T | T | Synonymous | 10 |
| DNAS_isl_121_MT581434 | 21551AtoT | N | I | Nonsynonymous | 4 |
| DNAS_isl_122_MT581413 | 15540CtoT | V | V | Synonymous | 4581 |
| DNAS_isl_122_MT581413 | 1163AtoT | I | F | Nonsynonymous | 4081 |
| DNAS_isl_122_MT581413 | 23403AtoG | D | G | Nonsynonymous | 6141 |
| DNAS_isl_122_MT581413 | 28883GtoC | G | R | Nonsynonymous | 4390 |
| DNAS_isl_122_MT581413 | 241CtoT | R | C | 5'UTR SNP | 766 |
| DNAS_isl_122_MT581413 | 3961CtoT | I | I | Synonymous | 1218 |
| DNAS_isl_122_MT581413 | 10252CtoT | F | F | Synonymous | 6316 |
| DNAS_isl_122_MT581413 | 28881GtoA | R | K | Nonsynonymous | 4386 |
| DNAS_isl_122_MT581413 | 28882GtoA | R | K | Nonsynonymous | 4390 |
| DNAS_isl_122_MT581413 | 22199GtoT | V | L | Nonsynonymous | 531 |
| DNAS_isl_122_MT581413 | 3037CtoT | F | F | Synonymous | 6450 |
| DNAS_isl_123_MT581425 | 3037CtoT | F | F | Synonymous | 3101 |
| DNAS_isl_123_MT581425 | 8371GtoT | Q | H | Nonsynonymous | 6945 |
| DNAS_isl_123_MT581425 | 25906GtoT | G | C | Nonsynonymous | 623 |
| DNAS_isl_123_MT581425 | 23541AtoT | Y | F | Nonsynonymous | 4843 |
| DNAS_isl_123_MT581425 | 27671TtoG | V | G | Nonsynonymous | 7 |
| DNAS_isl_123_MT581425 | 23403AtoG | D | G | Nonsynonymous | 1853 |
| DNAS_isl_123_MT581425 | 14408CtoT | P | L | Nonsynonymous | 1216 |
| DNAS_isl_123_MT581425 | 28882GtoA | R | K | Nonsynonymous | 1164 |
| DNAS_isl_123_MT581425 | 28883GtoC | G | R | Nonsynonymous | 1164 |
| DNAS_isl_123_MT581425 | 25714CtoT | L | F | Nonsynonymous | 411 |
| DNAS_isl_123_MT581425 | 28881GtoA | R | K | Nonsynonymous | 1159 |
| DNAS_isl_123_MT581425 | 4444GtoT | V | V | Synonymous | 3320 |
| DNAS_isl_123_MT581425 | 241CtoT | R | C | 5'UTR SNP | 119 |
| DNAS_isl_123_MT581425 | 29403AtoG | D | G | Nonsynonymous | 6339 |
| DNAS_isl_123_MT581425 | 1163AtoT | I | F | Nonsynonymous | 1603 |
| DNAS_isl_124_MT581422 | 28883GtoC | G | R | Nonsynonymous | 5620 |
| DNAS_isl_124_MT581422 | 28882GtoA | R | K | Nonsynonymous | 5620 |
| DNAS_isl_124_MT581422 | 28881GtoA | R | K | Nonsynonymous | 5611 |
| DNAS_isl_124_MT581422 | 23403AtoG | D | G | Nonsynonymous | 7995 |
| DNAS_isl_124_MT581422 | 4444GtoT | V | V | Synonymous | 3441 |
| DNAS_isl_124_MT581422 | 241CtoT | R | C | 5'UTR SNP | 614 |
| DNAS_isl_124_MT581422 | 3037CtoT | F | F | Synonymous | 5872 |
| DNAS_isl_124_MT581422 | 1163AtoT | I | F | Nonsynonymous | 4989 |
| DNAS_isl_124_MT581422 | 8371GtoT | Q | H | Nonsynonymous | 8001 |
| DNAS_isl_124_MT581422 | 25906GtoT | G | C | Nonsynonymous | 1921 |
| DNAS_isl_124_MT581422 | 29403AtoG | D | G | Nonsynonymous | 7713 |
| DNAS_isl_124_MT581422 | 14408CtoT | P | L | Nonsynonymous | 1125 |
| DNAS_isl_125_MT581423 | 28883GtoC | G | R | Nonsynonymous | 1930 |
| DNAS_isl_125_MT581423 | 8371GtoT | Q | H | Nonsynonymous | 7999 |
| DNAS_isl_125_MT581423 | 14408CtoT | P | L | Nonsynonymous | 1769 |
| DNAS_isl_125_MT581423 | 28882GtoA | R | K | Nonsynonymous | 1930 |
| DNAS_isl_125_MT581423 | 3037CtoT | F | F | Synonymous | 6337 |
| DNAS_isl_125_MT581423 | 28881GtoA | R | K | Nonsynonymous | 1928 |
| DNAS_isl_125_MT581423 | 29403AtoG | D | G | Nonsynonymous | 7825 |
| DNAS_isl_125_MT581423 | 241CtoT | R | C | 5'UTR SNP | 251 |
| DNAS_isl_125_MT581423 | 4444GtoT | V | V | Synonymous | 3517 |
| DNAS_isl_125_MT581423 | 1163AtoT | I | F | Nonsynonymous | 2726 |
| DNAS_isl_125_MT581423 | 23403AtoG | D | G | Nonsynonymous | 8017 |
| DNAS_isl_126_MT581412 | 28882GtoA | R | K | Nonsynonymous | 3716 |
| DNAS_isl_126_MT581412 | 28881GtoA | R | K | Nonsynonymous | 3712 |
| DNAS_isl_126_MT581412 | 14408CtoT | P | L | Nonsynonymous | 3476 |
| DNAS_isl_126_MT581412 | 23403AtoG | D | G | Nonsynonymous | 7735 |
| DNAS_isl_126_MT581412 | 20774GtoA | G | D | Nonsynonymous | 6214 |
| DNAS_isl_126_MT581412 | 28883GtoC | G | R | Nonsynonymous | 3716 |
| DNAS_isl_126_MT581412 | 3037CtoT | F | F | Synonymous | 5623 |
| DNAS_isl_126_MT581412 | 241CtoT | R | C | 5'UTR SNP | 596 |
| DNAS_isl_127_MT581433 | 7133GtoA | V | I | Nonsynonymous | 9 |
| DNAS_isl_127_MT581433 | 1288CtoT | C | C | Synonymous | 8017 |
| DNAS_isl_127_MT581433 | 28882GtoA | R | K | Nonsynonymous | 15 |
| DNAS_isl_127_MT581433 | 14408CtoT | P | L | Nonsynonymous | 720 |
| DNAS_isl_127_MT581433 | 28881GtoA | R | K | Nonsynonymous | 13 |
| DNAS_isl_127_MT581433 | 3037CtoT | F | F | Synonymous | 2673 |
| DNAS_isl_127_MT581433 | 28883GtoC | G | R | Nonsynonymous | 15 |
| DNAS_isl_127_MT581433 | 2048CtoT | L | L | Synonymous | 5004 |
| DNAS_isl_127_MT581433 | 23403AtoG | D | G | Nonsynonymous | 7980 |
| DNAS_isl_127_MT581433 | 13255CtoT | C | C | Synonymous | 7020 |
| DNAS_isl_127_MT581433 | 22193AtoT | N | Y | Nonsynonymous | 3218 |
| DNAS_isl_127_MT581433 | 4752CtoT | T | I | Nonsynonymous | 199 |
| DNAS_isl_128_MT581432 | 22193AtoT | N | Y | Nonsynonymous | 843 |
| DNAS_isl_128_MT581432 | 13255CtoT | C | C | Synonymous | 107 |
| DNAS_isl_128_MT581432 | 4752CtoT | T | I | Nonsynonymous | 7224 |
| DNAS_isl_128_MT581432 | 28881GtoA | R | K | Nonsynonymous | 6704 |
| DNAS_isl_128_MT581432 | 1288CtoT | C | C | Synonymous | 5004 |
| DNAS_isl_128_MT581432 | 3037CtoT | F | F | Synonymous | 7137 |
| DNAS_isl_128_MT581432 | 2048CtoT | L | L | Synonymous | 4806 |
| DNAS_isl_128_MT581432 | 28883GtoC | G | R | Nonsynonymous | 6713 |
| DNAS_isl_128_MT581432 | 28882GtoA | R | K | Nonsynonymous | 6713 |
| DNAS_isl_128_MT581432 | 23403AtoG | D | G | Nonsynonymous | 3301 |
| DNAS_isl_128_MT581432 | 241CtoT | R | C | 5'UTR SNP | 2926 |
| DNAS_isl_129_MT581427 | 2036GtoT | A | S | Nonsynonymous | 7960 |
| DNAS_isl_129_MT581427 | 28883GtoC | G | R | Nonsynonymous | 1721 |
| DNAS_isl_129_MT581427 | 1163AtoT | I | F | Nonsynonymous | 6529 |
| DNAS_isl_129_MT581427 | 28882GtoA | R | K | Nonsynonymous | 1721 |
| DNAS_isl_129_MT581427 | 14408CtoT | P | L | Nonsynonymous | 5144 |
| DNAS_isl_129_MT581427 | 28881GtoA | R | K | Nonsynonymous | 1717 |
| DNAS_isl_129_MT581427 | 3037CtoT | F | F | Synonymous | 5147 |
| DNAS_isl_129_MT581427 | 21691CtoT | F | F | Synonymous | 6656 |
| DNAS_isl_129_MT581427 | 23403AtoG | D | G | Nonsynonymous | 7966 |
| DNAS_isl_129_MT581427 | 241CtoT | R | C | 5'UTR SNP | 666 |
| DNAS_isl_129_MT581427 | 22000CtoG | H | Q | Nonsynonymous | 6653 |
| DNAS_isl_130_MT581428 | 1163AtoT | I | F | Nonsynonymous | 5381 |
| DNAS_isl_130_MT581428 | 2036GtoT | A | S | Nonsynonymous | 4163 |
| DNAS_isl_130_MT581428 | 3037CtoT | F | F | Synonymous | 7675 |
| DNAS_isl_130_MT581428 | 23403AtoG | D | G | Nonsynonymous | 7323 |
| DNAS_isl_130_MT581428 | 1912CtoT | S | S | Synonymous | 3920 |
| DNAS_isl_130_MT581428 | 241CtoT | R | C | 5'UTR SNP | 2736 |
| DNAS_isl_130_MT581428 | 28881GtoA | R | K | Nonsynonymous | 7551 |
| DNAS_isl_130_MT581428 | 28882GtoA | R | K | Nonsynonymous | 7562 |
| DNAS_isl_130_MT581428 | 28883GtoC | G | R | Nonsynonymous | 7562 |
| DNAS_isl_131_MT581436 | 23403AtoG | D | G | Nonsynonymous | 4093 |
| DNAS_isl_131_MT581436 | 3037CtoT | F | F | Synonymous | 7641 |
| DNAS_isl_131_MT581436 | 19972CtoT | P | S | Nonsynonymous | 72 |
| DNAS_isl_131_MT581436 | 28881GtoA | R | K | Nonsynonymous | 7980 |
| DNAS_isl_131_MT581436 | 28882GtoA | R | K | Nonsynonymous | 7992 |
| DNAS_isl_131_MT581436 | 2048CtoT | L | L | Synonymous | 7911 |
| DNAS_isl_131_MT581436 | 28883GtoC | G | R | Nonsynonymous | 7993 |
| DNAS_isl_131_MT581436 | 4752CtoT | T | I | Nonsynonymous | 8039 |
| DNAS_isl_131_MT581436 | 1288CtoT | C | C | Synonymous | 7917 |
| DNAS_isl_131_MT581436 | 241CtoT | R | C | 5'UTR SNP | 4226 |
| DNAS_isl_131_MT581436 | 22193AtoT | N | Y | Nonsynonymous | 1130 |
| DNAS_isl_132_MT581435 | 7036AtoG | L | L | Synonymous | 3 |
| DNAS_isl_132_MT581435 | 28882GtoA | R | K | Nonsynonymous | 7962 |
| DNAS_isl_132_MT581435 | 6471CtoT | T | I | Nonsynonymous | 4453 |
| DNAS_isl_132_MT581435 | 28883GtoC | G | R | Nonsynonymous | 7963 |
| DNAS_isl_132_MT581435 | 4892AtoG | T | A | Nonsynonymous | 45 |
| DNAS_isl_132_MT581435 | 11719GtoA | Q | Q | Synonymous | 3324 |
| DNAS_isl_132_MT581435 | 5194GtoT | L | L | Synonymous | 3314 |
| DNAS_isl_132_MT581435 | 23403AtoG | D | G | Nonsynonymous | 3919 |
| DNAS_isl_132_MT581435 | 28881GtoA | R | K | Nonsynonymous | 7943 |
| DNAS_isl_132_MT581435 | 22329CtoT | S | L | Nonsynonymous | 5980 |
| DNAS_isl_132_MT581435 | 241CtoT | R | C | 5'UTR SNP | 1893 |
| DNAS_isl_132_MT581435 | 3037CtoT | F | F | Synonymous | 7584 |
| DNAS_isl_133_MT581426 | 1163AtoT | I | F | Nonsynonymous | 432 |
| DNAS_isl_133_MT581426 | 25062GtoT | G | V | Nonsynonymous | 520 |
| DNAS_isl_133_MT581426 | 3037CtoT | F | F | Synonymous | 777 |
| DNAS_isl_133_MT581426 | 241CtoT | R | C | 5'UTR SNP | 206 |
| DNAS_isl_133_MT581426 | 28883GtoC | G | R | Nonsynonymous | 466 |
| DNAS_isl_133_MT581426 | 28881GtoA | R | K | Nonsynonymous | 464 |
| DNAS_isl_133_MT581426 | 28882GtoA | R | K | Nonsynonymous | 466 |
| DNAS_isl_133_MT581426 | 23403AtoG | D | G | Nonsynonymous | 1126 |
| DNAS_isl_133_MT581426 | 28253CtoT | F | F | Synonymous | 1188 |
| DNAS_isl_133_MT581426 | 18688TtoA | F | I | Nonsynonymous | 184 |
| DNAS_isl_133_MT581426 | 14408CtoT | P | L | Nonsynonymous | 204 |
| DNAS_isl_134_MT581424 | 28881GtoA | R | K | Nonsynonymous | 5855 |
| DNAS_isl_134_MT581424 | 28882GtoA | R | K | Nonsynonymous | 5865 |
| DNAS_isl_134_MT581424 | 27671TtoG | V | G | Nonsynonymous | 7 |
| DNAS_isl_134_MT581424 | 28883GtoC | G | R | Nonsynonymous | 5865 |
| DNAS_isl_134_MT581424 | 23403AtoG | D | G | Nonsynonymous | 7991 |
| DNAS_isl_134_MT581424 | 3037CtoT | F | F | Synonymous | 6225 |
| DNAS_isl_134_MT581424 | 241CtoT | R | C | 5'UTR SNP | 1239 |
| DNAS_isl_134_MT581424 | 14408CtoT | P | L | Nonsynonymous | 2430 |
| DNAS_isl_134_MT581424 | 1163AtoT | I | F | Nonsynonymous | 7744 |
| DNAS_isl_134_MT581424 | 4444GtoT | V | V | Synonymous | 4624 |
| DNAS_isl_134_MT581424 | 27479TtoA | V | E | Nonsynonymous | 17 |
| DNAS_isl_134_MT581424 | 8371GtoT | Q | H | Nonsynonymous | 7995 |
| DNAS_isl_134_MT581424 | 29403AtoG | D | G | Nonsynonymous | 7897 |
| DNAS_isl_134_MT581424 | 25906GtoT | G | C | Nonsynonymous | 2614 |
| DNAS_isl_135_MT581421 | 23403AtoG | D | G | Nonsynonymous | 7988 |
| DNAS_isl_135_MT581421 | 28883GtoC | G | R | Nonsynonymous | 4650 |
| DNAS_isl_135_MT581421 | 8371GtoT | Q | H | Nonsynonymous | 7998 |
| DNAS_isl_135_MT581421 | 28881GtoA | R | K | Nonsynonymous | 4640 |
| DNAS_isl_135_MT581421 | 28882GtoA | R | K | Nonsynonymous | 4650 |
| DNAS_isl_135_MT581421 | 1163AtoT | I | F | Nonsynonymous | 7205 |
| DNAS_isl_135_MT581421 | 3037CtoT | F | F | Synonymous | 5646 |
| DNAS_isl_135_MT581421 | 241CtoT | R | C | 5'UTR SNP | 834 |
| DNAS_isl_135_MT581421 | 14408CtoT | P | L | Nonsynonymous | 3704 |
| DNAS_isl_135_MT581421 | 4444GtoT | V | V | Synonymous | 5248 |
| DNAS_isl_135_MT581421 | 29403AtoG | D | G | Nonsynonymous | 7923 |
| DNAS_isl_136_MT581417 | 23403AtoG | D | G | Nonsynonymous | 7963 |
| DNAS_isl_136_MT581417 | 3688CtoT | H | H | Synonymous | 7880 |
| DNAS_isl_136_MT581417 | 15324CtoT | N | N | Synonymous | 7991 |
| DNAS_isl_136_MT581417 | 14408CtoT | P | L | Nonsynonymous | 931 |
| DNAS_isl_136_MT581417 | 241CtoT | R | C | 5'UTR SNP | 1235 |
| DNAS_isl_136_MT581417 | 3037CtoT | F | F | Synonymous | 5770 |
| DNAS_isl_137_MT581419 | 23403AtoG | D | G | Nonsynonymous | 7999 |
| DNAS_isl_137_MT581419 | 14408CtoT | P | L | Nonsynonymous | 1334 |
| DNAS_isl_137_MT581419 | 241CtoT | R | C | 5'UTR SNP | 878 |
| DNAS_isl_137_MT581419 | 15324CtoT | N | N | Synonymous | 7998 |
| DNAS_isl_137_MT581419 | 3688CtoT | H | H | Synonymous | 7849 |
| DNAS_isl_137_MT581419 | 7528CtoT | V | V | Synonymous | 7980 |
| DNAS_isl_137_MT581419 | 3037CtoT | F | F | Synonymous | 6767 |
| DNAS_isl_138_MT581416 | 6578CtoT | L | F | Nonsynonymous | 4731 |
| DNAS_isl_138_MT581416 | 23403AtoG | D | G | Nonsynonymous | 6657 |
| DNAS_isl_138_MT581416 | 15324CtoT | N | N | Synonymous | 7996 |
| DNAS_isl_138_MT581416 | 241CtoT | R | C | 5'UTR SNP | 521 |
| DNAS_isl_138_MT581416 | 3037CtoT | F | F | Synonymous | 6647 |
| DNAS_isl_138_MT581416 | 3688CtoT | H | H | Synonymous | 7948 |
| DNAS_isl_139_MT581418 | 3037CtoT | F | F | Synonymous | 4915 |
| DNAS_isl_139_MT581418 | 23268TtoG | I | S | Nonsynonymous | 3614 |
| DNAS_isl_139_MT581418 | 241CtoT | R | C | 5'UTR SNP | 503 |
| DNAS_isl_139_MT581418 | 15324CtoT | N | N | Synonymous | 7998 |
| DNAS_isl_139_MT581418 | 23230CtoT | N | N | Synonymous | 3467 |
| DNAS_isl_139_MT581418 | 23403AtoG | D | G | Nonsynonymous | 4742 |
| DNAS_isl_139_MT581418 | 3688CtoT | H | H | Synonymous | 7904 |
| DNAS_isl_139_MT581418 | 14408CtoT | P | L | Nonsynonymous | 2554 |
| DNAS_isl_140_MT581411 | 3037CtoT | F | F | Synonymous | 6651 |
| DNAS_isl_140_MT581411 | 1163AtoT | I | F | Nonsynonymous | 7283 |
| DNAS_isl_140_MT581411 | 14408CtoT | P | L | Nonsynonymous | 5823 |
| DNAS_isl_140_MT581411 | 28883GtoC | G | R | Nonsynonymous | 4289 |
| DNAS_isl_140_MT581411 | 28687CtoT | A | A | Synonymous | 8011 |
| DNAS_isl_140_MT581411 | 28882GtoA | R | K | Nonsynonymous | 4290 |
| DNAS_isl_140_MT581411 | 23403AtoG | D | G | Nonsynonymous | 7984 |
| DNAS_isl_140_MT581411 | 28881GtoA | R | K | Nonsynonymous | 4281 |
| DNAS_isl_140_MT581411 | 241CtoT | R | C | 5'UTR SNP | 875 |
| DNAS_isl_140_MT581411 | 3871GtoT | K | N | Nonsynonymous | 3678 |
| DNAS_isl_141_MT581430 | 20438GtoT | S | I | Nonsynonymous | 7905 |
| DNAS_isl_141_MT581430 | 4201GtoT | M | I | Nonsynonymous | 1084 |
| DNAS_isl_141_MT581430 | 24208CtoT | I | I | Synonymous | 1481 |
| DNAS_isl_141_MT581430 | 1188CtoT | S | L | Nonsynonymous | 4500 |
| DNAS_isl_141_MT581430 | 3037CtoT | F | F | Synonymous | 7098 |
| DNAS_isl_141_MT581430 | 23403AtoG | D | G | Nonsynonymous | 7978 |
| DNAS_isl_141_MT581430 | 12369CtoT | T | I | Nonsynonymous | 7973 |
| DNAS_isl_141_MT581430 | 241CtoT | R | C | 5'UTR SNP | 707 |
| DNAS_isl_141_MT581430 | 26527CtoT | A | V | Nonsynonymous | 2730 |
| DNAS_isl_142_MT581431 | 3037CtoT | F | F | Synonymous | 6498 |
| DNAS_isl_142_MT581431 | 241CtoT | R | C | 5'UTR SNP | 609 |
| DNAS_isl_142_MT581431 | 8389CtoT | N | N | Synonymous | 7973 |
| DNAS_isl_142_MT581431 | 15324CtoT | N | N | Synonymous | 7993 |
| DNAS_isl_142_MT581431 | 23868GtoT | G | V | Nonsynonymous | 2254 |
| DNAS_isl_142_MT581431 | 18412GtoT | V | F | Nonsynonymous | 2675 |
| DNAS_isl_142_MT581431 | 23403AtoG | D | G | Nonsynonymous | 7981 |
| DNAS_isl_142_MT581431 | 3217TtoC | S | S | Synonymous | 3835 |
| DNAS_isl_142_MT581431 | 15850GtoA | D | N | Nonsynonymous | 1740 |
| DNAS_isl_143_MT581429 | 12369CtoT | T | I | Nonsynonymous | 7965 |
| DNAS_isl_143_MT581429 | 4201GtoT | M | I | Nonsynonymous | 4679 |
| DNAS_isl_143_MT581429 | 26527CtoT | A | V | Nonsynonymous | 3536 |
| DNAS_isl_143_MT581429 | 20438GtoT | S | I | Nonsynonymous | 7972 |
| DNAS_isl_143_MT581429 | 3037CtoT | F | F | Synonymous | 5741 |
| DNAS_isl_143_MT581429 | 29587GtoA | P | P | Synonymous | 7890 |
| DNAS_isl_143_MT581429 | 23403AtoG | D | G | Nonsynonymous | 7756 |
| DNAS_isl_143_MT581429 | 241CtoT | R | C | 5'UTR SNP | 489 |
| DNAS_isl_144_MT581420 | 8371GtoT | Q | H | Nonsynonymous | 8000 |
| DNAS_isl_144_MT581420 | 28883GtoC | G | R | Nonsynonymous | 4350 |
| DNAS_isl_144_MT581420 | 28882GtoA | R | K | Nonsynonymous | 4350 |
| DNAS_isl_144_MT581420 | 4444GtoT | V | V | Synonymous | 6755 |
| DNAS_isl_144_MT581420 | 29403AtoG | D | G | Nonsynonymous | 7994 |
| DNAS_isl_144_MT581420 | 1163AtoT | I | F | Nonsynonymous | 7567 |
| DNAS_isl_144_MT581420 | 241CtoT | R | C | 5'UTR SNP | 700 |
| DNAS_isl_144_MT581420 | 28881GtoA | R | K | Nonsynonymous | 4340 |
| DNAS_isl_144_MT581420 | 23403AtoG | D | G | Nonsynonymous | 7997 |
| DNAS_isl_144_MT581420 | 14408CtoT | P | L | Nonsynonymous | 4050 |
| DNAS_isl_144_MT581420 | 3037CtoT | F | F | Synonymous | 5279 |
| DNAS_isl_145_MT581415 | 14408CtoT | P | L | Nonsynonymous | 4068 |
| DNAS_isl_145_MT581415 | 4298GtoT | V | L | Nonsynonymous | 5804 |
| DNAS_isl_145_MT581415 | 28903GtoT | M | I | Nonsynonymous | 7993 |
| DNAS_isl_145_MT581415 | 23403AtoG | D | G | Nonsynonymous | 7995 |
| DNAS_isl_145_MT581415 | 3037CtoT | F | F | Synonymous | 6163 |
| DNAS_isl_145_MT581415 | 3688CtoT | H | H | Synonymous | 7949 |
| DNAS_isl_145_MT581415 | 241CtoT | R | C | 5'UTR SNP | 1002 |
| DNAS_isl_145_MT581415 | 15324CtoT | N | N | Synonymous | 8001 |
| DNAS_isl_146_MT581410 | 3037CtoT | F | F | Synonymous | 5434 |
| DNAS_isl_146_MT581410 | 3961CtoT | I | I | Synonymous | 7968 |
| DNAS_isl_146_MT581410 | 23202CtoT | T | I | Nonsynonymous | 7945 |
| DNAS_isl_146_MT581410 | 28882GtoA | R | K | Nonsynonymous | 5404 |
| DNAS_isl_146_MT581410 | 241CtoT | R | C | 5'UTR SNP | 549 |
| DNAS_isl_146_MT581410 | 28883GtoC | G | R | Nonsynonymous | 5405 |
| DNAS_isl_146_MT581410 | 28881GtoA | R | K | Nonsynonymous | 5401 |
| DNAS_isl_146_MT581410 | 1163AtoT | I | F | Nonsynonymous | 6631 |
| DNAS_isl_146_MT581410 | 23403AtoG | D | G | Nonsynonymous | 7970 |
| DNAS_isl_147_MT566438 | 29403AtoG | D | G | Nonsynonymous | 7983 |
| DNAS_isl_147_MT566438 | 28882GtoA | R | K | Nonsynonymous | 5513 |
| DNAS_isl_147_MT566438 | 28883GtoC | G | R | Nonsynonymous | 5513 |
| DNAS_isl_147_MT566438 | 241CtoT | R | C | 5'UTR SNP | 662 |
| DNAS_isl_147_MT566438 | 27671TtoG | V | G | Nonsynonymous | 5 |
| DNAS_isl_147_MT566438 | 28881GtoA | R | K | Nonsynonymous | 5507 |
| DNAS_isl_147_MT566438 | 3037CtoT | F | F | Synonymous | 5505 |
| DNAS_isl_147_MT566438 | 23403AtoG | D | G | Nonsynonymous | 8005 |
| DNAS_isl_147_MT566438 | 14408CtoT | P | L | Nonsynonymous | 2772 |
| DNAS_isl_147_MT566438 | 25906GtoT | G | C | Nonsynonymous | 672 |
| DNAS_isl_147_MT566438 | 4444GtoT | V | V | Synonymous | 2882 |
| DNAS_isl_147_MT566438 | 8371GtoT | Q | H | Nonsynonymous | 7999 |
| DNAS_isl_147_MT566438 | 1163AtoT | I | F | Nonsynonymous | 6733 |
| DNAS_isl_148_MT566437 | 3688CtoT | H | H | Synonymous | 856 |
| DNAS_isl_148_MT566437 | 15324CtoT | N | N | Synonymous | 3218 |
| DNAS_isl_148_MT566437 | 241CtoT | R | C | 5'UTR SNP | 195 |
| DNAS_isl_148_MT566437 | 23403AtoG | D | G | Nonsynonymous | 3920 |
| DNAS_isl_148_MT566437 | 14408CtoT | P | L | Nonsynonymous | 813 |
| DNAS_isl_148_MT566437 | 3037CtoT | F | F | Synonymous | 1150 |
| DNAS_isl_149_MT566436 | 29403AtoG | D | G | Nonsynonymous | 7986 |
| DNAS_isl_149_MT566436 | 15895CtoT | L | L | Synonymous | 4555 |
| DNAS_isl_149_MT566436 | 28881GtoA | R | K | Nonsynonymous | 4243 |
| DNAS_isl_149_MT566436 | 3037CtoT | F | F | Synonymous | 5388 |
| DNAS_isl_149_MT566436 | 23403AtoG | D | G | Nonsynonymous | 7983 |
| DNAS_isl_149_MT566436 | 28882GtoA | R | K | Nonsynonymous | 4249 |
| DNAS_isl_149_MT566436 | 241CtoT | R | C | 5'UTR SNP | 1528 |
| DNAS_isl_149_MT566436 | 28883GtoC | G | R | Nonsynonymous | 4249 |
| DNAS_isl_149_MT566436 | 8371GtoT | Q | H | Nonsynonymous | 7999 |
| DNAS_isl_149_MT566436 | 25906GtoT | G | C | Nonsynonymous | 2485 |
| DNAS_isl_149_MT566436 | 4444GtoT | V | V | Synonymous | 6377 |
| DNAS_isl_149_MT566436 | 14408CtoT | P | L | Nonsynonymous | 3635 |
| DNAS_isl_149_MT566436 | 1163AtoT | I | F | Nonsynonymous | 7091 |
| DNAS_isl_150_MT566435 | 29403AtoG | D | G | Nonsynonymous | 7948 |
| DNAS_isl_150_MT566435 | 1163AtoT | I | F | Nonsynonymous | 5377 |
| DNAS_isl_150_MT566435 | 24887TtoC | F | L | Nonsynonymous | 4947 |
| DNAS_isl_150_MT566435 | 28881GtoA | R | K | Nonsynonymous | 5190 |
| DNAS_isl_150_MT566435 | 28882GtoA | R | K | Nonsynonymous | 5196 |
| DNAS_isl_150_MT566435 | 2041TtoC | T | T | Synonymous | 6537 |
| DNAS_isl_150_MT566435 | 28883GtoC | G | R | Nonsynonymous | 5196 |
| DNAS_isl_150_MT566435 | 4444GtoT | V | V | Synonymous | 4656 |
| DNAS_isl_150_MT566435 | 3037CtoT | F | F | Synonymous | 6262 |
| DNAS_isl_150_MT566435 | 14408CtoT | P | L | Nonsynonymous | 2371 |
| DNAS_isl_150_MT566435 | 8371GtoT | Q | H | Nonsynonymous | 8004 |
| DNAS_isl_150_MT566435 | 23403AtoG | D | G | Nonsynonymous | 7982 |
| DNAS_isl_150_MT566435 | 241CtoT | R | C | 5'UTR SNP | 773 |
| DNAS_isl_151_MT566434 | 3037CtoT | F | F | Synonymous | 5533 |
| DNAS_isl_151_MT566434 | 25906GtoT | G | C | Nonsynonymous | 2728 |
| DNAS_isl_151_MT566434 | 29403AtoG | D | G | Nonsynonymous | 7974 |
| DNAS_isl_151_MT566434 | 241CtoT | R | C | 5'UTR SNP | 1703 |
| DNAS_isl_151_MT566434 | 10979GtoA | V | M | Nonsynonymous | 2434 |
| DNAS_isl_151_MT566434 | 28882GtoA | R | K | Nonsynonymous | 5162 |
| DNAS_isl_151_MT566434 | 4444GtoT | V | V | Synonymous | 6734 |
| DNAS_isl_151_MT566434 | 28883GtoC | G | R | Nonsynonymous | 5162 |
| DNAS_isl_151_MT566434 | 8371GtoT | Q | H | Nonsynonymous | 7998 |
| DNAS_isl_151_MT566434 | 14408CtoT | P | L | Nonsynonymous | 3428 |
| DNAS_isl_151_MT566434 | 1163AtoT | I | F | Nonsynonymous | 7094 |
| DNAS_isl_151_MT566434 | 23403AtoG | D | G | Nonsynonymous | 7984 |
| DNAS_isl_151_MT566434 | 27671TtoG | V | G | Nonsynonymous | 32 |
| DNAS_isl_151_MT566434 | 28881GtoA | R | K | Nonsynonymous | 5155 |
