## Supplemental Table 3 for "Large scale genomic and evolutionary study reveals SARS-CoV-2 virus isolates from Bangladesh strongly correlate with European origin and not with China"

**Supplementary Table 03.: Summary of variants associated with clinical parameters.**

A total of 37 variants returned p values less than 0.05 for various parameters. Variant 3961 C to T was associated with developing sore throat and diarrhoea. Variant 14408 C to T was inversely associated with coughing. Variants 22199 G to T, 19593 C to T, 13902 T to C, 774 C to T, and 21597 C to T were significantly associated with the development of sore throat, 28881 G to A, 28882 G to A, and 28883 G to C showed association with development of chest pain, 21123 G to T was associated with the development of anorexia and 29118 C to T, 28178 G to T, 29262 G to T were associated with an individual suffering from pneumonia. Variants 4105 G to T, 3456 A to G, 28305 A to G, 26051 G to A, 24685 T to C were associated with protection from a loss of taste and smell. 28292 C to A, 4300 G to T, 26526 G to T, 17193 G to T, 2731 G to A, 98264 A to G, 12025 C to T, 8311 C to T, 714 G to A, 8366 G to A, 18859 G to T, 9416 G to A, 20808 G to A were all significantly related to the onset or protection from skin rash. 28079 G to T and 3053 G to T were associated with itching and redness of eyes. Variants 28580 G to A, 2363 C to T, and the 3871 G to T variants significantly correlated with individuals who were asymptomatic.

| **Variant(s)** | **Variant Type** | **Associated Clinical Symptom** | **Effect on Disease Phenotype** | **Chi Squared P-Value of Significance** | **Fisher's Test P-value of Significance** | **Remarks** |
| --- | --- | --- | --- | --- | --- | --- |
| 14408CtoT | SNV | Cough | No Cough | 0.035140609 | 0.02691376 |  |
| 12525CtoT, 26828GtoT, 29578CtoA | Co-Variants (SNVs) | Fever | Unclear | 0.029428639 | 0.02856609 | Co-variants |
| 3961CtoT | SNV | Sore Throat | Causes Sore Throat | 0.0005889 | 0.000570637 |  |
|  | Co-Variants (SNVs) | Diarrhea | Protects from Diarrhea | NA | 0.03658523 |  |
| 22199GtoT, 19593CtoT, 13902TtoC, 774CtoT, 21597CtoT | MNV | Sore Throat | Causes Sore Throat | 0.025402255 | 0.01798972 | Co-variants |
| 28881GtoA, 28882GtoA, 28883GtoC | SNV | Chest Pain | Causes Chest Pain | 0.025142178 | 0.02478889 | Co-variants |
| 21123GtoT | SNV | Anorexia | Causes Anorexia | NA | 0.03548946 | Only four isolates contained variant |
| 4105GtoT, 3456AtoG, 28305AtoG, 26051GtoA, 24685TtoC | Co-Variants (SNVs) | Loss of taste/smell | Does not cause loss of taste | NA | 0.04632518 | Co-variants |
| 29118CtoT, 28178GtoT, 29262GtoT | Co-Variants (SNVs) | Pneumonia | Causes Pneumonia | 4.43E-07 | 0.009615385 | Co-variants |
| 28292CtoA, 4300GtoT, 26526GtoT, 17193GtoT | Co-Variants (SNVs) | Skin Rash | Unclear | 0.004675087 | 0.02884615 | Co-variants |
| 2731GtoA | SNV | Skin Rash | Unclear | 0.004675087 | 0.02884615 |  |
| 98264AtoG, 12025CtoT, 8311CtoT, 714GtoA, 8366GtoA, 18859GtoT, X9416GtoA, 20808GtoA | Co-Variants (SNVs) | Skin Rash | Unclear | 0.004675087 | 0.02884615 | Co-variants |
| 28079GtoT | SNV | Redness and Itching in Eyes | Unclear | NA | 0.03846154 |  |
| 3053GtoT | SNV | Redness and Itching in Eyes | Unclear | NA | 0.01923077 |  |
| 3871GtoT | SNV | Symptomatic Status | Asymptomatoc | 0.04531751 | 0.02723718 |  |
| 28580GtoA, 2363CtoT | Co-Variants (SNVs) | Symptomatic Status | Asymptomatoc | 0.04531751 | 0.02723718 | Co-variants |
